# Human SerRS/SIRT2 complex structure reveals cross regulation between translation and NAD^+^ metabolism

**DOI:** 10.1101/2025.09.23.677902

**Authors:** Qian Zhang, Huimin Zhang, Marscha Hirschi, Sheng Li, Gabriel C. Lander, Jie Yang, Xiang-Lei Yang

## Abstract

Life at the cellular level depends on effective coordination between diverse processes. Here we uncover a novel cross-regulation between metabolism and translation through a 3.2 Å cryo-EM structure of human cytosolic seryl-tRNA synthetase (SerRS) bound to sirtuin-2 (SIRT2), an NAD^+^-dependent deacetylase. This interaction, naturally triggered by the NAD^+^ metabolite ADP-ribose (ADPR), resembles substrate binding and block SIRT2’s active site. Interestingly, SerRS acetylation is not required for this interaction. SIRT2 binding sterically and allosterically impedes tRNA binding to SerRS, lowering charged tRNA^Ser^ level and protein synthesis activity. Key interaction residues in both proteins emerged simultaneously in vertebrates, suggesting co-evolution for cross-regulation. Given ADPR’s accumulation under stress, the ADPR-induced SerRS/SIRT2 interaction likely serves as a cell-protective response.

## Introduction

Aminoacyl-tRNA synthetases (aaRSs) are a family of evolutionarily conserved enzymes that catalyze the ligation of amino acid (aa) to their cognate tRNAs. Each proteinogenic aa requires a specific aaRS for the ligation reaction. Using serine (Ser) as an example, the aminoacylation reaction starts with the aa being selected by seryl-tRNA synthetase (SerRS, encoded by SARS1) and activated with energy released from ATP hydrolysis to form the reaction intermediate seryl-adenylate (Ser-AMP). Subsequently, the seryl group is transferred from Ser-AMP to the 3’-end of either serine tRNA (tRNA^Ser^) or selenocysteine tRNA (tRNA^Sec^), resulting in serylated-tRNA^Ser/Sec^ (Fig. 1a). Collectively, these aaRSs-catalyzed reactions create a pool of matched aa-tRNAs, essential for decoding mRNA while providing aa building blocks for ribosomal protein synthesis. In addition, many aaRSs, especially those in higher organisms, have been reported to play regulatory roles in a broad range of biological processes, including metabolism^1, 2, 3^.

**Figure 1.**
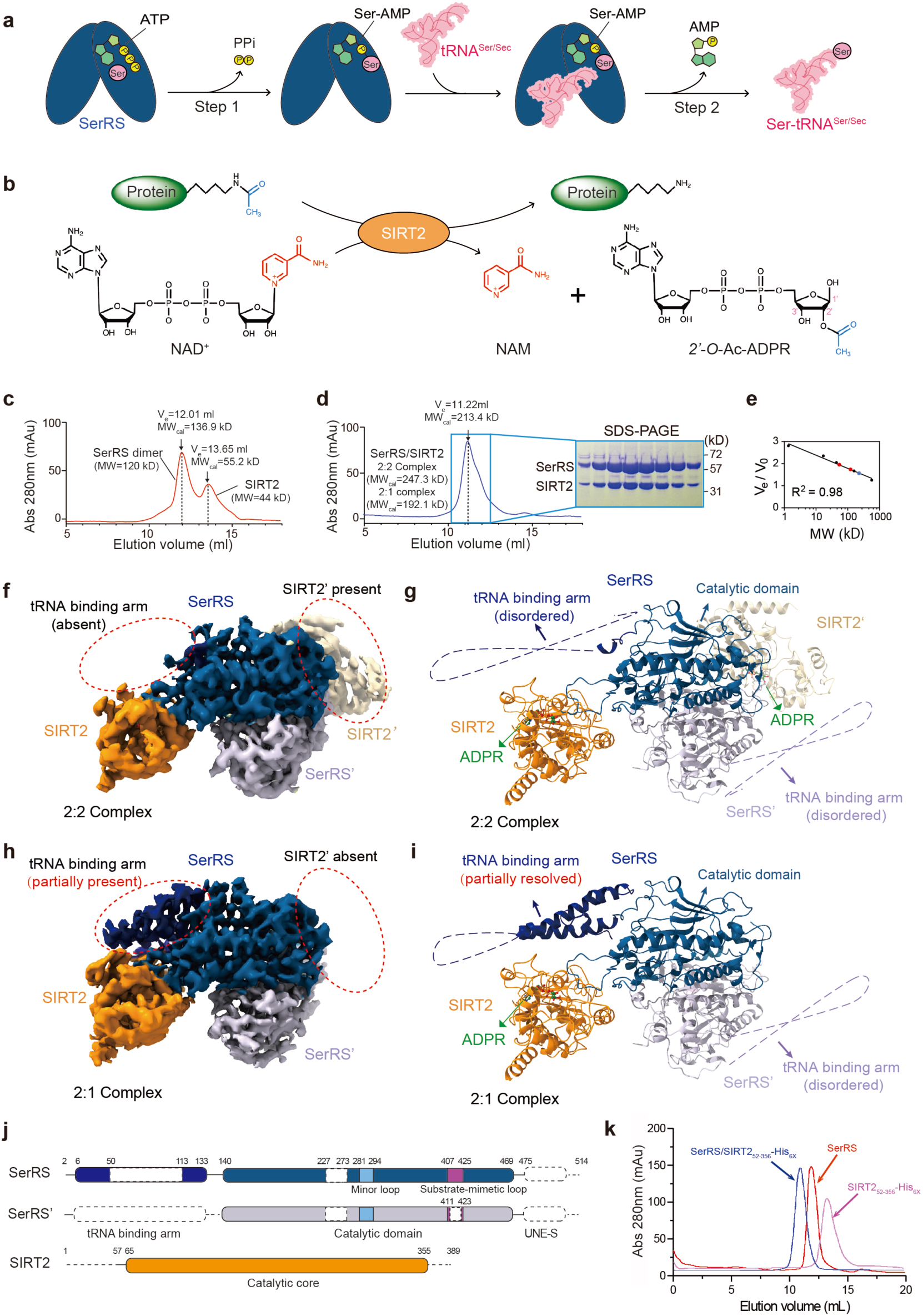
Cryo-EM structure of SerRS/SIRT2 complex. (a) Two-step tRNA serinylation reaction catalyzed by SerRS. In step 1, ATP and serine bind to the catalytic pocket of SerRS, forming seryl-adenylate intermediate with the release of pyrophosphate (PPi); in step 2, the activated serine is transferred to SerRS-bound tRNA^Ser/Sec^, resulting in the formation followed by releasing of Ser-tRNA^Ser^ or Ser-tRNA^Sec^and AMP. (b) SIRT2-mediated NAD^+^-dependent protein deacetylation reaction. The acetyl group from a modified lysine residue of the substrate protein is transferred to the ADP-ribose moiety of NAD^+^, generating 2’-O-Ac-ADP-ribose (OAADPR) and a deacetylated protein. Nicotinamide (NAM) is released as a byproduct. (c) Size exclusion chromatography (SEC) profile of pre-mixed SerRS and SIRT2 proteins shows no evidence of interaction. Recombinant SerRS (tag-free) and SIRT2_FL_-His_6X_ (Full length SIRT2 with a C-terminal His_6X_ tag) was purified separately. The estimated molecular weight (MW_cal_) was calculated from the elution volume (V_e_) using a standard curve generated with molecular weight marker proteins. The theoretical molecular weight (MW) of SerRS and SIRT2 are also indicated. (d) SEC profile of the SerRS/SIRT2 complex formed through co-expression in *E. coli* (left panel) and SDS-PAGE analysis of the peak fractions (right panel). Both SerRS and SIRT2 are present in the single asymmetric peak. The peak of the complex had an estimated MW_cal_ of 213.4 kD, which is between the MW_cal_ of 2:2 complex (136.9+55.2+55.2=247.3(kD)) and the 2:1 complex (136.9+55.2=192.1(kD)). (e) Standard calibration curve depicting elution volume and molecular size marker proteins: vitamin B12 (1.35kDa), horse myoglobin (17kDa), chicken ovalbumin (44kDa), bovine γ-globulin (158kDa), thyroglobulin (670 kDa). Red dots represent SerRS and SIRT2 peaks from (c), while the blue dot represent the SerRS/SIRT2 complex from (d). (f) Cryo-EM map of the SerRS/SIRT2 2:2 complex at an overall resolution of 3.94 Å. The two SIRT2 molecules are labeled in orange (SIRT2) and wheat (SIRT2’), while the two subunits of SerRS dimer are in blue (SerRS) and light purple (SerRS’). The red dash line circles indicate the differences between the 2:2 complex and 2:1 complex in (h). (g) Overall cryo-EM density-fitted model of the SerRS/SIRT2 2:2 complex corresponding to (f). The disordered tRNA binding arms in both SerRS and SerRS’ are represented with dotted lines. An ADPR molecule (colored in green) was found in the co-factor binding pocket of each SIRT2. (h) Cryo-EM map of the SerRS/SIRT2 2:1 complex at 3.87 Å overall resolution, with a 3.2 Å resolution at the SerRS/SIRT2 interface. The red dash line circles indicate the differences between the 2:2 complex and 2:1 complex. (i) Overall cryo-EM density-fitted model of the SerRS/SIRT2 2:1 complex corresponding to (h). The partially disordered tRNA binding arm of SerRS and the fully disordered tRNA binding arm of SerRS’ are illustrated with dotted lines. An ADPR molecule (colored in green) is observed in the co-factor binding pocket of SIRT2. (j) Schematic illustrations of the domain structure of SerRS and SIRT2, with disordered regions in the SerRS/SIRT2 2:1 complex indicated by uncolored dashed lines. (k) SEC profiles showing that SerRS forms stable complex with the catalytical core (Res. G52-S356) of SIRT2 (SIRT2_52-356_**-**His_6X_).

Sirtuins are lysine deacetylases that utilize nicotinamide adenine dinucleotide (NAD^+^) as the co-substrate to remove posttranslational *N*ε-lysine acetylation from both histone and non-histone proteins (Fig. 1b), thereby regulating protein functions^4^. The level of NAD^+^ in the cell represents its energy potential. As the main oxidizing and reducing agents, NAD^+^ and its reduced form, NADH, facilitate oxidation-reduction (redox) reactions during metabolic processes, extracting ATP and biological building blocks (e.g., nucleotides, phospholipids, and amino acids) from nutrients^5, 6, 7, 8^. By using NAD^+^ as the co-substrate, the functions of sirtuin family are intrinsically linked to cellular metabolism and energy homeostasis^9, 10^.

A NAD^+^ molecule is composed of two moieties: ADP-ribose (ADPR) and nicotinamide (NAM) (Fig. 1b). When a *N*ε-lysine acetylated substrate protein is engaged by a NAD^+^-bound sirtuin, the acetyl group on lysine forms an intermediate with NAD^+^ and triggers the cleavage of the glycosidic bond between NAM and ADPR. Following the release of NAM from the enzyme, the acetyl-group is transferred from the Nε-lysine to the remaining ADPR moiety, producing *2′-*O-acetyl-ADP-ribose (OAADPR)^11^. Upon release from the sirtuin, OAADPR can be further converted to ADPR in the cell through an esterase or an acetyltransferase^12, 13^.

Our previous work identified a direct interaction between human cytosolic SerRS and sirtuin 2 (SIRT2)^14^. Of the seven mammalian family members (SIRT1-7), SIRT2 is the only sirtuin primarily localized in the cytoplasm^10^. Among other targets, SIRT2 deacetylates α-tubulin lysine 40 in response to cellular NAD^+^ levels^15, 16^. A small portion of SIRT2 also shuttles into the nucleus in a cell cycle dependent manner^17^, targeting histone H4 lysine 16^17^ and transcription factors^18, 19^. Interestingly, a fraction of SerRS is also found in the nucleus due to the presence of a nuclear localization sequence (NLS) in its C-terminal UNE-S domain, which is specific to higher eukaryotes^20^. In the nucleus, SerRS and SIRT2 act as partners to transcriptionally repress VEGFA expression, thereby playing an essential role in vascular development and angiogenesis regulation^14, 20^. However, the details of how SerRS and SIRT2 interact with each other and how this affects their cytosolic functions remain unknown.

Here, we determined the cryo-EM structures of the SerRS/SIRT2 complex. Remarkably, a ADPR was revealed in the active site of SIRT2, mediating a key hydrogen bond with the K414 residue of SerRS. Interestingly, SerRS binding resembles that of a SIRT2 substrate, however, acetylation of SerRS is not required. Instead, SerRS binding modulates the deacetylase activity of SIRT2, offering insights into how SerRS regulates cellular processes beyond translation. Conversely, SIRT2 binding disrupts the aminoacylation activity of SerRS by directly competing with and allosterically regulating its interaction with tRNA^Ser^, leading to reduced protein synthesis. The ADPR-mediated mutual inhibition between SerRS and SIRT2 suggests that this interaction may function as a cellular stress sensor, regulating translation in response to environmental or metabolic changes. Furthermore, the co-evolution of key residues in both SerRS and SIRT2 across vertebrates highlights the important role of the SerRS/SIRT2 complex in maintaining cellular homeostasis in higher organisms. To our knowledge, this study provides the first structure of a tRNA synthetase complexed with a protein outside the translational machinery and the first evidence of a substrate-like negative regulator of sirtuins.

## Results

### SerRS and SIRT2 interact through their catalytic domain

To enable structural analysis of the SerRS/SIRT2 interaction, we purified recombinant human SerRS and SIRT2 through separate overexpression in *E. coli*. Analytical size-exclusion chromatography (SEC) of pre-mixed SerRS and SIRT2 proteins revealed no detectable interaction, as only two distinct peaks corresponding to dimeric SerRS and monomeric SIRT2 were observed (Fig. 1c). The estimated molecular weight (MW_cal_) of SerRS dimer (136.9kD) and SIRT2 (55.2 kD), based on the elution volume (V_e_), were both greater than their theoretical MW values (i.e., 120 kD and 44 kD, respectively), presumably due to the flexible conformations of certain domains and regions within the proteins (see below). Considering the complex formation typically requires co-factors and/or co-folding processes in the cell, we co-expressed SerRS (tag-free) with SIRT2 (His_6X_-tagged) in *E. coli*. His_6X_-tag affinity purification followed by SEC yielded a stable SerRS/SIRT2 complex that eluted as a single peak (Fig. 1d). The presence of both SerRS and SIRT2 in the complex was confirmed by denaturing SDS-PAGE (Fig. 1d). Based on elution volume (V_e_), the peak of SerRS/SIRT2 complex had an estimated molecular weight of 213.4 kD (Fig. 1d, e), suggesting a mixture of 2:1 complex (SerRS dimer with 1 SIRT2) and 2:2 complex (SerRS dimer with 2 SIRT2).

The peak fraction was analyzed using single-particle cryo-EM to determine the structure of the SerRS/SIRT2 complex (Supplementary Table 1). Two-dimensional classifications of particle images also revealed a mixture of 2:2 and 2:1 complexes (Extended Data Fig. 1a, b), consistent with the SEC analysis (Fig. 1d). A total of 1,113,696 particles displaying high-resolution secondary features were selected for processing (Extended Data Fig. 1b). Both 2:2 and 2:1 complexes were selected for *ab initio* Non-Uniform (NU)-refinement with C2 symmetry, respectively, resulting in a 3D reconstruction of the SerRS/SIRT2 2:2 complex at 3.94 Å overall resolution (Fig. 1f, g; Extended Data Fig. 1) and the 2:1 complex at 4.01 Å overall resolution (Extended Data Fig. 1). Further local refinement achieved by masking different regions of the 2:1 complex map, yielded a 3.87 Å reconstruction map for the 2:1 complex and a 3.2 Å map focused on the SerRS/SIRT2 interface (Fig. 1h, i; Extended Data Fig. 1). The 2:2 complex features a dimeric SerRS at the center, with two SIRT2 molecules each binding to one of the SerRS subunits (Fig. 1g). Essentially, the similar interaction is observed in the 2:1 complex, where only one SIRT2 molecule is bound to the SerRS dimer (Fig. 1i).

In both the 2:2 and 2:1 SerRS/SIRT2 complexes, the N- and C- terminal regions of SIRT2 (M1-E56 and S356-Q389, respectively) are unresolved (Fig. 1j), suggesting they are not involved in the SerRS interaction. Indeed, deletion of both N- and C-terminals regions of SIRT2 does not affect the SerRS/SIRT2 interaction (Fig. 1k). In the 2:1 complex, the N- terminal tRNA binding arm (V2-R133) was partially resolved in one SerRS subunit but remained almost completely disordered in the other (SerRS’) (Fig 1i, j). Conversely, in the 2:2 complex, the tRNA binding arms of both SerRS subunits remained unresolved (Fig 1g, j). Furthermore, the C-terminal UNE-S domains of SerRS (P475-D514) are unresolved in both complexes (Fig. 1j). Consistently, the tRNA binding arm and UNE-S domain of SerRS have been shown to be dispensable for SIRT2 interaction^14^. Therefore, the SerRS/SIRT2 interaction is mediated merely through the catalytic domains of SerRS (H140-K469) and SIRT2 (E65-Q355) (Fig. 1j).

### ADP-ribose triggers SerRS/SIRT2 interaction

To better visualize the context of the SerRS/SIRT2 interaction, we modeled the disordered part of the tRNA binding arm for the SIRT2-bound SerRS subunit in the 2:1 complex (Fig. 2a). We found an extra density in the active site of SIRT2 corresponding to ADPR (Fig. 2a, b) and no density for NAM. To confirm that this bound ligand is ADPR rather than NAD^+^, we extracted the ligand from the co-purified SerRS/SIRT2 protein and subjected it to ESI triple quad mass spectrometry analysis. Only ADPR was detected, with no traces of NAD^+^ or OAADPR (Extended Data Fig. 2). In contrast, neither ADPR, OAADPR nor NAD^+^ was detected from the separately purified SIRT2 and SerRS proteins, confirming that ADPR is the co-factor that triggers the formation of a stable complex between SerRS and SIRT2. Consistently, the conformation of SIRT2 in complex with SerRS more closely resembles that of SIRT2 in complex with ADPR (PDB 5D7O) than any other SIRT2 states (i.e., apo, in complex with NAD^+^ and substrate, or in complex with reaction intermediates mimicry)^21, 22, 23, 24^ (Extended Data Fig. 3).

**Figure 2.**
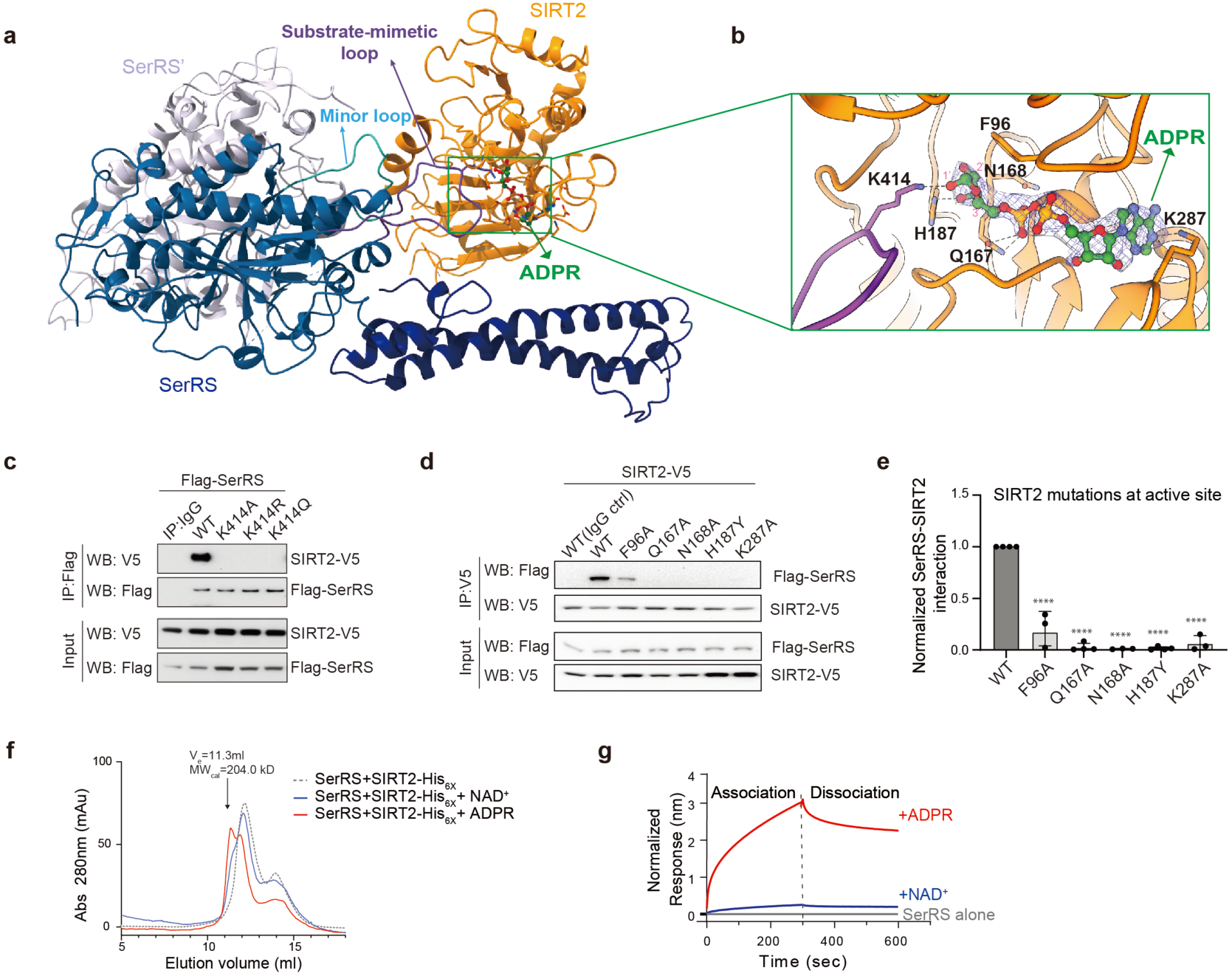
SerRS/SIRT2 interaction is induced by ADP-ribose. (a) Ribbon representation of the overall SerRS/SIRT2 2:1 complex structure. The disordered region of the tRNA binding arm in SerRS subunit bound with SIRT2 is modeled in based on the crystal structure of SerRS in complex with a seryl-adenylate analog (PDB 4L87). The core region of the structure is modeled from the 3.2 Å SerRS/SIRT2 1:1 map. Two loops in SerRS involved in SIRT2 interaction are highlighted: the minor loop (cyan) and the substrate-mimetic loop (purple). The ADPR molecule captured in the active site of SIRT2 is presented as stick-and-ball representation. (b) Zoom-in view of the SIRT2 active site, showing the interaction between ADPR and K414 from SerRS substrate-mimetic loop. The ε-amino group of K414 forms a hydrogen bond with the O1’ atom in the terminal ribose moiety of ADPR. (c) Co-immunoprecipitation (Co-IP) assay showing the critical role of K414 in the SerRS/SIRT2 interaction. Mutations in K414 abolish the interaction. (d) Representative co-IP result showing that SIRT2 active site residues involved in ADPR binding are critical for the SerRS/SIRT2 interaction. (e) Quantification of Co-IP result in (d). Error bars represent ±SD. \*\*\*\**p* < 0.0001, Student’s t-test, n=4. (f) SEC analysis of separately purified recombinant SerRS and SIRT2_FL_-His_6X_, with and without NAD^+^ or ADPR. The V_e_ and MW_cal_ of the ADPR-induced SerRS/SIRT2 complex peak are indicated. (g) Biolayer interferometry analysis showing ADPR induces a stronger SerRS/SIRT2 interaction than NAD^+^ at same concentration (0.5 mM). SIRT2_FL_-His_6X_ was immobilized on Anti-Penta-HIS sensor tips, and background binding signal of SIRT2 with SerRS olny was subtracted.

Within the SIRT2 active site, the ADPR molecule directly interacts with SerRS through a hydrogen bond between O1’ of the ribose and the ε-amino group of K414 in SerRS (Fig. 2b). The K414 residue is positioned at the tip of a long loop (substrate-mimetic loop; residues R407-M425; see below), which protrudes from the catalytic domain of SerRS and inserts into the substrate binding groove of SIRT2. Substitution of K414 with either alanine (SerRS^K414A^), arginine (SerRS^K414R^), or glutamine (SerRS^K414Q^) completely abolished the SerRS/SIRT2 interaction (Fig. 2c), confirming the essential role of K414 in mediating this interaction. Different from the clear density observed for the substrate-mimetic loop in the SIRT2-bound SerRS subunit, the majority of this loop (G411-V423) is disordered in the SerRS’ subunit (Fig 1j), indicating that the SIRT2 interaction stabilizes the conformation of the substrate-mimetic loop in SerRS.

To further demonstrate that the SerRS/SIRT2 interaction is dependent on the presence of ADPR, we introduced single point substitutions of multiple SIRT2 active site residues that are not directly involved in SerRS binding but important in ADPR binding^25^ (Fig. 2b). All substitutions, including SIRT2^F96A^, SIRT2^Q167A^, SIRT2^N168A^, SIRT2^H187Y^, and SIRT2^K287A^, severely impacted or completely abolished the SerRS/SIRT2 interaction (Fig. 2d, e), confirming the interaction is ADPR-dependent. Indeed, adding ADPR into a mixture of the separately purified SerRS and SIRT2 proteins resulted in the formation of a new peak, as detected by SEC (Fig. 2f), corresponding to the SerRS/SIRT2 complex. Interestingly, NAD^+^ can also induce the SerRS/SIRT2 interaction *in vitro*, albeit with a weaker effect than ADPR (Fig. 2f). Bio-layer interferometry (BLI) further confirms a significantly stronger SerRS/SIRT2 interaction in the presence of ADPR compared to NAD^+^ at the same concentration (Fig. 2g).

### SerRS suppresses SIRT2 deacetylation activity

The insertion of SerRS K414 into the active site of SIRT2 raised the possibility that human SerRS K414 could be acetylated, acting as a substrate of SIRT2. Therefore, our structure may have captured a deacetylated K414 along with the ADPR product prior to their release from the enzyme. However, the loss of interaction between acetylation mimetic mutant SerRS^K414Q^ and SIRT2 (Fig. 2c) does not support this possibility. To further evaluate this, we subjected both the recombinant SerRS protein alone and the co-purified SerRS/SIRT2 complex for trypsin digestion followed by mass spectrometry analysis. In both cases, a peptide containing the K414 site (^410^YGQTK^414^) was successfully detected (Extended Data Fig. 4a-c); however, no acetylation of this peptide was detected (Extended Data Fig. 4c). In contrast, more than a dozen lysine acetylation sites were identified in SerRS alone and/or in the SerRS/SIRT2 complex (Extended Data Fig. 4d), including the previously identified K323 site from mammalian cells^26^ (Extended Data Fig. 4e). Given that SerRS is acetylated at various sites, we also conducted a deacetylation reaction to assess if SerRS can act as a substrate of SIRT2. We used an Ac-K40-containing α-tubulin peptide as a positive control. In the presence of NAD^+^, SIRT2 deacetylated the α-tubulin peptide, as indicated by the detection of the reaction products OAADPR and ADPR by ESI triple quad mass spectrometry. In contrast, no reaction products were detected when using SerRS as the substrate (Extended Data Fig. 5). Therefore, although the SerRS/SIRT2 interaction resembles that between SIRT2 and its substrates, our results indicate that the SerRS is not a substrate of SIRT2 and the interaction is independent of SIRT2 catalysis.

Given that the substrate-mimetic loop of SerRS obstructs the substrate binding groove of SIRT2 in an ADPR-induced manner (Fig. 2a, 3a), we speculate that SerRS binding could inhibit SIRT2 deacetylation activity. To test the hypothesis, we conducted an *in vitro* deacetylation assay using the α-tubulin Ac-K40-containing peptide as the substrate. Indeed, SerRS inhibits the deacetylase activity of SIRT2 in a concentration dependent manner (Fig. 3b). Notably, SerRS mutants (i.e., K414Q and K414R), that are unable to bind to SIRT2, exhibited a significant reduction in inhibitory effects (Fig. 3b), confirming that the inhibition is specifically mediated by the SerRS/SIRT2 interaction. Interestingly, in the absence of SerRS, the addition of ADPR at 5-10 times of the NAD^+^ concentration had no impact on SIRT2 activity (Fig. 3c), consistent with previous reports that ADPR is a poor inhibitor of sirtuins^27^. However, in the presence of SerRS, the addition of ADPR enhanced the inhibitory effect of SerRS on SIRT2 activity in a concentration-dependent manner (Fig. 3c), presumably due to the enhanced interaction between SerRS and SIRT2 induced by ADPR.

**Figure 3.**
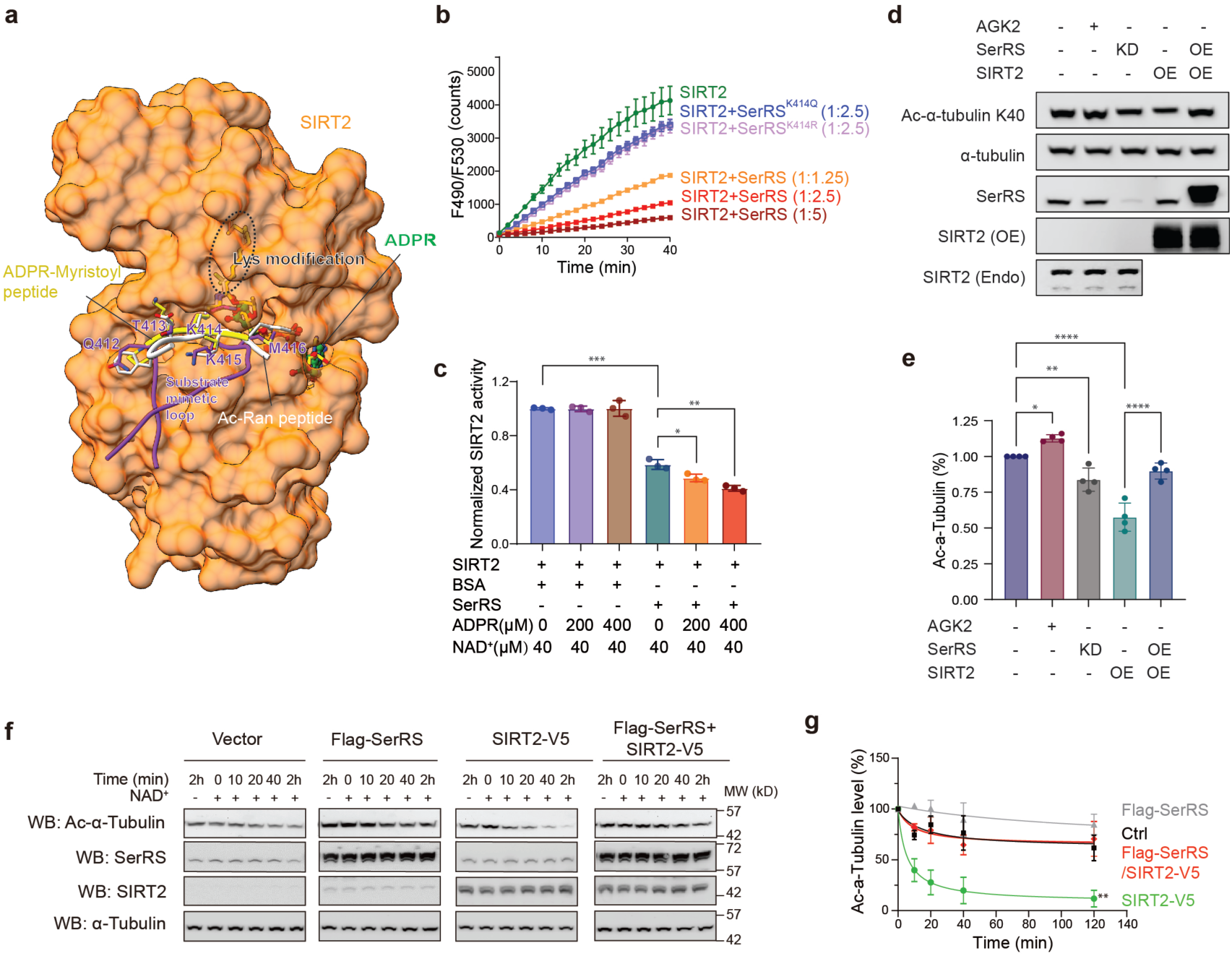
SerRS suppresses SIRT2 deacetylation activity. (a) SerRS interacts with SIRT2 through substrate mimicry. The SIRT2 substrate Ac-Ran peptide (PDB ID: 5FYQ; white), the reaction intermediate mimicry peptide (ADPR-Myristoyl peptide, PDB ID: 4X3O; yellow), and SerRS substrate-mimetic loop (purple) are modeled into the SIRT2 substrate binding groove via superposition of SIRT2 from each complex. Residues in the SerRS Substrate-mimetic loop that are in proximity to SIRT2, along with corresponding residues from the other two peptides, are shown in sticks. (b) *In vitro* deacetylation assay demonstrating that SerRS, but not SerRS K414 mutants, inhibits SIRT2**-**His_6X_ activity in a dose-dependent manner. SIRT2 activity was measured via fluorescence intensity (excitation: 490 nm, emission: 530 nm) from the cleaved substrate peptide. (c) The inhibitory effect of SerRS on SIRT2 deacetylation is enhanced by ADPR in a concentration-dependent manner. BSA was used as a non-specific control. (d) Representative western blot showing α-tubulin K40 acetylation levels in 293AD cells, evaluating the impact of SerRS knockdown (KD) and overexpression (OE) on SIRT2 activity. A SIRT2 specific inhibitor AGK2 (5μM) was used as a control. (e) Quantification of the acetylation levels of α-tubulin K40 from (d). \**p* < 0.05, ** *p* < 0.01, **** *p* < 0.0001, two-way ANOVA test, n=4. (f) Representative deacetylation assays using HEK293 cell lysates overexpressing vector control, SerRS, SIRT2 or both SerRS and SIRT2. (g) Quantification of the Ac-α-tubulin level from (f). Ac-α-Tubulin levels at each time point were normalized to the 0 min time point, which was defined as 100% for each reaction. Error bars represent ±SME for duplicate experiments. ** *p* < 0.01, two-way ANOVA test, n=2.

To further investigate whether SerRS can inhibit SIRT2 activity in cells, we monitored the acetylation level of α-tubulin K40 in HEK293AD cells following modulations of SerRS and SIRT2 expression. Knockdown of SerRS resulted in increased SIRT2 deacetylation activity, leading to lower acetylation level on α-tubulin K40, contrasting with the opposite effect observed using a SIRT2-specific inhibitor AGK2 (Fig. 3d, e). Additionally, co-overexpression of SerRS significantly diminished the enhanced SIRT2 deacetylation of α-tubulin K40 caused by SIRT2 overexpression (Fig. 3d, e). Similar results of SerRS inhibiting SIRT2 activity were observed in HEK293 cell lysates treated with NAD^+^, where we monitored the acetylation level of α-tubulin K40 over time (Fig. 3f, g). These findings indicate that SerRS can repress SIRT2 deacetylation activity both *in vitro* and in cellular contexts.

### SIRT2 inhibits SerRS tRNA aminoacylation activity and global translation

Comparing the structure of SerRS/SIRT2 and SerRS/tRNA^Sec^ (PDB ID: 4RQF) complexes reveals an overlap between the SIRT2 and tRNA binding sites on SerRS (Fig. 4a), suggesting a mutual exclusion of SIRT2 and tRNA binding. Additionally, 3D variability analysis (3DVA) suggests a dynamic correlation between SIRT2 binding and disordering of the tRNA binding arm in SerRS across the dimer (Extended Data Fig. 6, Supplementary Video). In the 2:1 complex, the tRNA binding arm is partially resolved in the SIRT2-bound SerRS subunit but entirely disordered in the SIRT2-free subunit (Fig. 1h, i); while in the 2:2 complex with two bound SIRT2 molecules, both SerRS subunits exhibited completely disordered tRNA binding arms (Fig. 1f, g). Hydrogen deuterium exchange mass spectrometry analysis (HDX-MS) confirms the increased flexibility of tRNA binding arm upon SIRT2 binding. Compared to separately purified proteins, deuterium incorporation in the SerRS/SIRT2 complex decreased in the catalytic domain of both SIRT2 and SerRS (Extended Data Fig. 7), consistent with binding-induced conformational stabilization. However, the tRNA binding arm overall exhibits increased deuterium incorporation (Extended Data Fig. 7a, c), consistent with SIRT2 binding-enhanced flexibility observed from the structure. Therefore, SIRT2 binding directly blocks tRNA binding to one SerRS subunit and allosterically disturb tRNA binding to the other subunit (SerRS’) (Fig. 4a, b). This is the case even in the absence of a second SIRT2, as seen in the 2:1 complex (Fig 1h, i).

**Figure 4.**
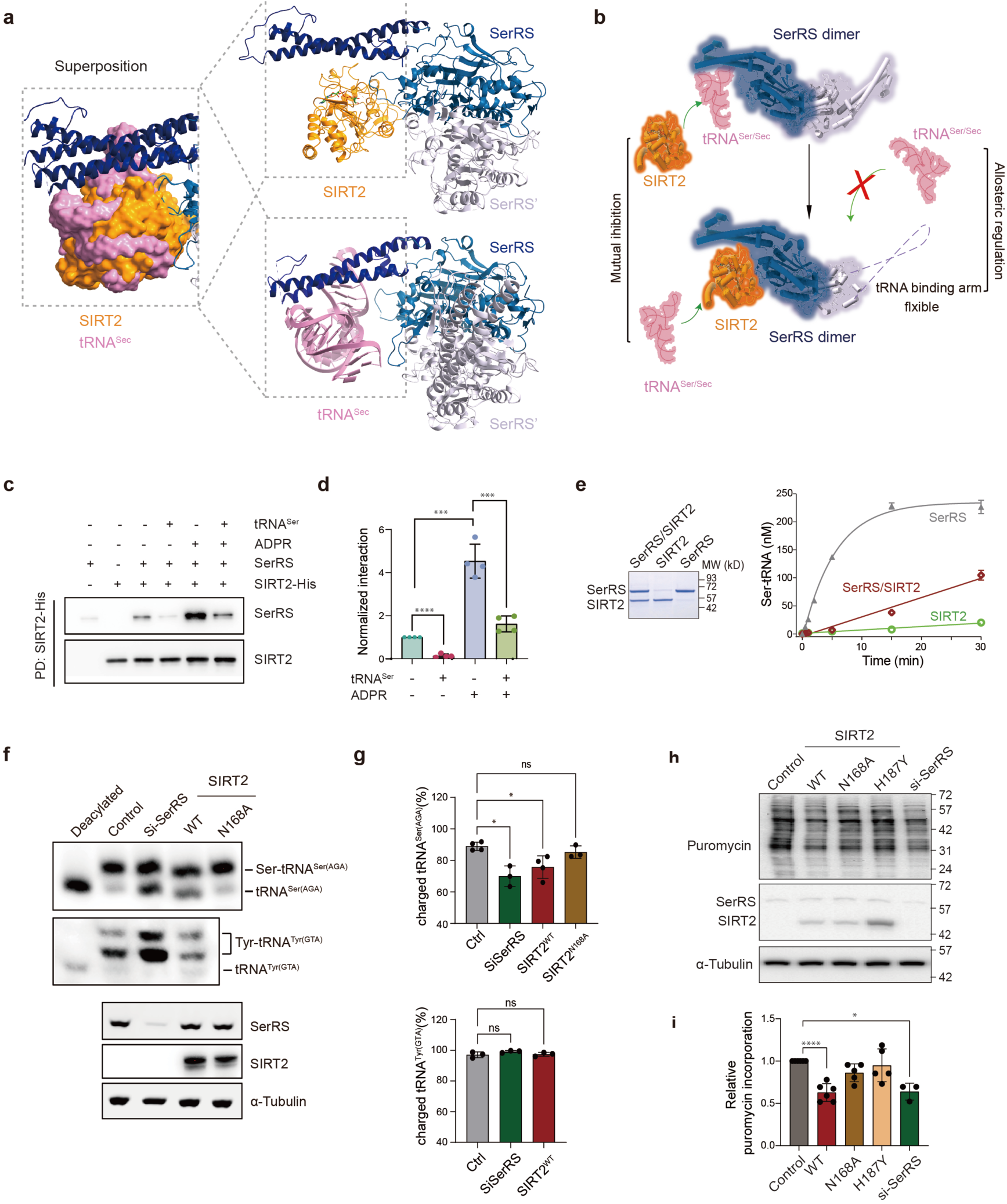
SIRT2 inhibits SerRS tRNA aminoacylation activity and global translation. (a) Superposition of the SerRS/SIRT2 2:1 complex cryo-EM structure with the SerRS/tRNA^Sec^ crystal structure (PDB ID: 4RQF), revealing overlap between SIRT2 and tRNA binding sites on SerRS. (b) Schematic illustration of SIRT2’s inhibition to the tRNA^Ser/Sec^/SerRS interaction via mutual inhibition and allosteric regulation. (c) Ni-NTA pull-down assay showing that the SerRS/SIRT2 interaction is strongly inhibited by tRNA^Ser^ (1:1 mixture of tRNA^Ser(GCT)^ and tRNA^Ser(CGA)^). (d) Quantification of the pull-down assay from (c). Error bars represent ±SD. \*\*\**p* < 0.001, **** *p* < 0.0001, Student’s t-test, n=4. (e) *In vitro* minoacylation assays showing that SIRT2 inhibits SerRS aminoacylation activity. Recombinant SerRS, SIRT2 and SerRS/SIRT2 complex used in the assay were analyzed by SDS-PAGE to ensure equal amounts of each component were included (right panel). Error bars represent ±SD from duplicate experiments. (f) Representative acidic gel northern blot analysis demonstrates that SIRT2 specifically inhibits SerRS charging activity in HEK293AD cells. The knockdown of SerRS and the overexpression of SIRT2 were confirmed by western blot. (g) Quantification of charged tRNA^Ser(AGA)^ and charged tRNA^Tyr(GTA)^ levels in (f). Error bars represent ±SD. * *p* <0.05, two-way ANOVA test, n=3-4. (h) Puromycin incorporation assay shows that either knockdown of SerRS or overexpression of SIRT2 reduces global protein synthesis in HEK293AD cells, while overexpression of SIRT2 mutants unable to interact with SerRS does not. (i) Quantification of puromycin incorporation from (h). Error bars represent ±SD. \**p* < 0.05, **** *p* < 0.0001, Student’s t-test, n=3-6.

To investigate SIRT2’s effect on tRNA binding to SerRS, a pull-down assay with His_6X_-tagged SIRT2 was conducted. A weak interaction between SerRS and SIRT2 was detected, even in the absence of ADPR, while ADPR significantly enhanced the interactions as expected (Fig. 4c, d). Importantly, tRNA^Ser^ inhibited the SerRS and SIRT2 interaction in both the presence and absence of ADPR (Fig. 4c, d). Moreover, decreased tRNA aminoacylation activity of the co-purified SerRS/SIRT2 complex, compared with SerRS alone, was demonstrated by an *in vitro* aminoacylation assay (Fig. 4e). To further explore the inhibitory effect of SIRT2 on tRNA^Ser^ aminoacylation in cells, an acidic gel northern blot was performed to assess the tRNA charging level from HEK293AD cells. Overexpression of SIRT2 reduced the charging level of tRNA^Ser^ from 94% to 71%, similar to the effect of SerRS knockdown (68%) (Fig 4f, g). In contrast, overexpression of SIRT2^N168A^ mutant, which cannot interact with SerRS, exhibited no significant impact (Fig. 4f, g). Notably, the inhibitory effect of SIRT2 on tRNA aminoacylation was specific to tRNA^Ser^, as the charging level of tRNA^Tyr^ was not affected upon overexpression of SIRT2 (Fig 4f, g). Two forms of charged tRNA^Tyr^ were detected, presumably corresponding to differences in the modification status of the tRNA^28^. Furthermore, the SUnSET assay^29^, which measures global translation activity via puromycin incorporation, indicated that overexpression of WT SIRT2, but not the N168A and H187Y mutants, inhibited global translation, consistent with the effect observed with SerRS knockdown (Fig, 4h, i). Collectively, these results suggest that SerRS/SIRT2 interaction suppresses the aminoacylation of tRNA^Ser^ and inhibits protein synthesis.

### SerRS/SIRT2 interacting sites are likely co-evolved

In addition to the ADPR-mediated interaction, SerRS engages with SIRT2 through two critical loops within its catalytic domain (Fig. 2a). The substrate-mimetic loop (residues R407-M425), where K414 reside located, directly interacts with the substrate binding groove of SIRT2 (Fig.3a, 5a). Single mutations at Q412 (SerRS^Q412S^) and M416 (SerRS^M416A^), but not K415 (SerRS^K415A^) and K419 (SerRS^K419A^), abolished the SIRT2 interaction, highlighting the critical role of the substrate-mimetic loop in the SerRS/SIRT2 interaction (Fig. 5d; Extended Data Fig. 8a, b). A second loop (minor loop; residues R281-Y294) also contributes to interaction with SIRT2 (Fig. 5b), partially through stabilizing the substrate-mimetic loop (Fig. 5c). Mutations in the minor loop (e.g., D282A and E283A) do impact the SerRS/SIRT2 interaction, but the effects are generally weaker than those in the substrate-mimetic loop (Fig. 5e; Extended Data Fig. 8c, d).

**Figure 5.**
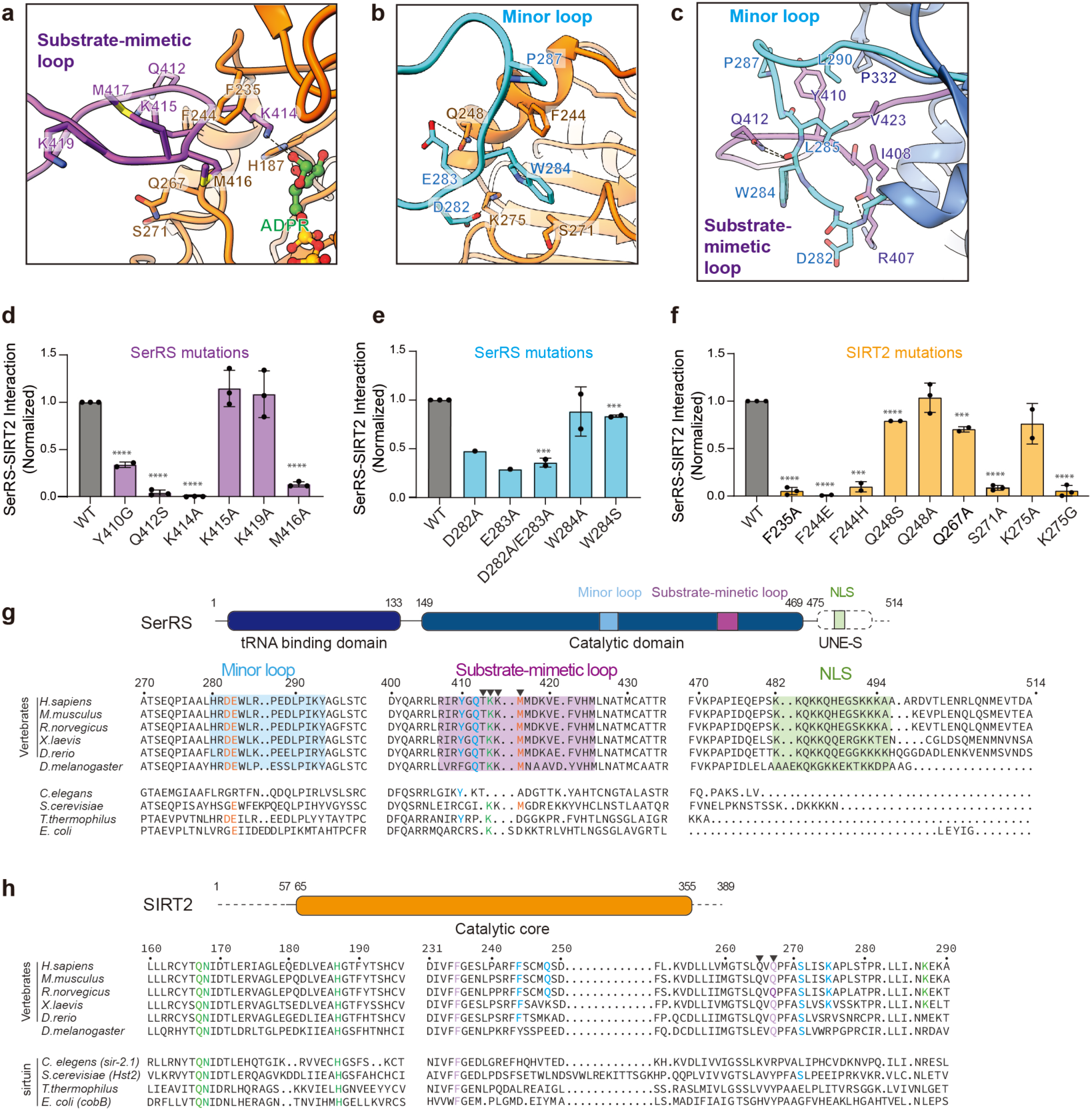
SerRS/SIRT2 interaction and mutagenesis studies. (a-b) SerRS/SIRT2 interaction at the substrate-mimetic loop (a) and minor loop (b) of SerRS. Residues near the interface are shown in sticks. Hydrogen bonds are indicated in black dashed lines. (c) Interactions between the substrate-mimetic loop and minor loop of SerRS. (d, e and f) Quantification of mutagenesis and Co-IP assays in Extended Data Fig. 8. The mutations Y410G, Q412S, K414A and M416A in substrate-mimetic loop (d) and D282A and E283A in the minor loop (e) of SerRS either abolish or weaken SIRT2 binding; while mutations F235A, F244E, F244H, S271A, and K275G in SIRT2 (f) either abolish or weaken the SerRS binding. Relative levels of the Co-IPed protein versus IPed protein were quantified by ImageJ. Error bars represent ±SD for duplicate or triplicate experiments. ** *p* < 0.01, **** *p* < 0.0001. **** *p* < 0.0001. Student’s t-test, n=2-3. (g) Sequence alignment of SerRS orthologs highlighting conserved key residues involved in the SerRS/SIRT2 interaction across vertebrates. minor loop is highlighted in cyan, the substrate-mimetic loop in purple, and the previously reported NLS in green. Key residues in SerRS that interact directly with SIRT2 are colored in orange, while those involved in the interaction between the minor loop and substrate-mimetic loop are in cyan. K414 from the substrate-mimetic loop is indicated in green. (h) Sequence alignment of SIRT2 orthologs showing that the residues important for SerRS binding are conserved in vertebrates. Residues forming hydrogen bonds with ADPR are colored in green, those interacting with the minor loop of SerRS in cyan and those interact with Substrate-mimetic loop of SerRS in purple. In both (g) and (h), residues exhibiting mainchain interactions with their binding partner are indicated with a black triangle above.

On the SIRT2 side, multiple residues, including F235, F244, and S271, are crucial for the SerRS interaction (Fig. 5a, b, f; Extended Data Fig. 8e-g). F235 of SIRT2 makes a cation-π interaction with K414 of SerRS (Fig. 5a) and SIRT2^F235A^ mutation disrupted this interaction (Fig. 5f, Extended Data Fig. 8f). Mutations of F244 (i.e., SIRT2^F244H^ and SIRT2^F244E^) also abolished the interaction with SerRS (Fig. 5f; Extended Data Fig. 8e-g), providing an explanation for the selectivity of SerRS for SIRT2 over SIRT1^14^, which contains a histidine at this position (Extended Data Fig. 9).The non-conservation of F244 among sirtuins suggests a specific interaction between SerRS and SIRT2 (Extended Data Fig. 9), within sirtuin family. As expected, SIRT2^F244H^ remains at least partial deacetylase activity (Extended Data Fig. 8h, i). while SIRT2^S271A^ can no longer interact with SerRS but maintains full deacetylase activity (Fig. 5f; Extended Data Fig. 8g-i). In contrast, the K275G mutation abolished both SerRS/SIRT2 interaction and SIRT2 activity (Fig. 5f, Extended Data Fig. 8g-i). The F244H and S271A mutations may serve as valuable tools for exploring the biological implications of the SerRS/SIRT2 interaction in future studies.

The residues in SerRS that are critical for SIRT2 interaction (e.g., Q412, K414, and M416 in the substrate-mimetic loop; D282 and E283 in the minor loop) are conserved from *Drosophila* to humans (Fig. 5g). A similar conservation pattern is found in SIRT2 residues that are directly involved in the SerRS interaction (e.g., F235 and S271) (Fig. 5h). In contrast, the active site residues of SIRT2 (e.g., Q167, N168, and H187), which interact with SerRS via the ADPR co-factor, are strictly conserved throughout evolution, apparently for functions beyond the SerRS/SIRT2 interaction (Fig. 5h). The evolutionary co-emergence of these direct interacting residues on both sides supports the idea that these enzymes have co-evolved for mutual regulation, rather than functioning within a traditional enzyme-substate relationship.

## Discussion

Depletion of cytoplasmic NAD^+^ by 50% or more can block metabolic processes and eventually lead to loss of cell viability^30^. Although the interconversion between NAD^+^ and NADH are required to drive the metabolic reactions, these reactions do not significantly alter the overall cytoplasmic NAD^+^ levels, which are significantly higher than those of NADH. Cytoplasmic NAD^+^/NADH ratios range between 60-700 in a typical eukaryotic cell^31, 32^. However, when cells are under stress, major NAD^+^ consuming enzymes, such as sirtuins and PARPs, are activated, resulting in the production of NAM and OAADPR/ADPR during or immediately following the reaction, or through further enzyme-catalyzed hydrolysis^33^. Unlike NAM, which can be recycled into new NAD^+^ through the salvage pathway (the predominant NAD^+^ synthesis pathway)^5^, OAADPR/ADPR does not contribute to NAD^+^ synthesis. Instead, both ADPR and OAADPR have been implicated as signaling molecules. For example, in mammalian cells, accumulation of free ADPR from oxidative and nitrosative stress induces activation of TRPM2 cation channels, causing Ca^2+^ influx^34, 35, 36^. OAADPR has a similar activity as ADPR in gating TRPM2^37^. In yeast cells, a buildup of OAADPR/ADPR has been linked to a metabolic shift that lowers endogenous reactive oxygen species and diverts glucose from glycolysis towards pentose phosphate pathways to generate nicotinamide adenine dinucleotide phosphate (NADPH)^38^.

In this study, we have shown NAD^+^ metabolites, particularly ADPR, can potently induce the formation of SerRS/SIRT2 complex *in vitro*. We propose that the accumulation of OAADPR/ADPR under cellular stress would induce the SerRS/SIRT2 interaction in the cell. As we have shown here, on one hand, the interaction inhibits SIRT2 activity and therefore would help preserve NAD^+^. On the other hand, and perhaps more importantly, formation of SerRS/SIRT2 complex inhibits serylation of tRNA and protein synthesis, and therefore would preserve ATP and amino acids, especially serine. In addition to being a proteingenic amino acid, serine serves as a central node in the metabolic network^39^. It is required for the biosynthesis of many biological building blocks, including nucleotides, other amino acids (e.g., glycine, cysteine, and methionine), and lipids (e.g., phospholipids and sphingolipids). Serine is also a major carbon donor for the biosynthesis of antioxidant glutathione (GSH) and reducing agent nicotinamide adenine dinucleotide phosphate (NADPH), therefore supporting cellular redox balance. Serine deficiencies are linked to many diseases including diabetes, macular diseases, and peripheral neuropathy^40, 41, 42, 43^. Despite these key roles in metabolism, the cellular level of serine is the lowest among all protein genic amino acids^44^. Therefore, ADPR induced SerRS/SIRT2 interaction and mutual regulation could be an important resource-preservation mechanism that protects cells under stress. Conversely, when the recourses are abundant, as represented in the abundance of tRNA, ATP, and serine, SerRS would be engaged in tRNA serylation and protein synthesis, becoming less available for interacting with SIRT2. In this way, translation and post-translational modification-mediated functional regulations can be coordinated in response to cellular metabolism and energy status. Interestingly, GlyRS has also been reported to interact with SIRT2, but not SIRT1, and the interaction also inhibits the deacetylation activity of SIRT2^45^. Although the impact of SIRT2 on GlyRS’s catalytic activity and whether the GlyRS and SIRT2 interaction is triggered by ADPR or other stress-related signaling molecules are not known, the fact that serine and glycine are biosynthetically linked and mutually convertible supports the relevance of the SerRS/SIRT2 interaction to amino acid metabolism and suggests that both SerRS and GlyRS could be involved in the regulation.

In addition to the role of ADPR, the SerRS and SIRT2 interaction is also mediated through the substrate-mimetic loop of SerRS and the substrate binding groove of SIRT2, resembling the interaction between SIRT2 and its substrates (Fig. 3a), despite acetylation of SerRS is not required for the interaction. Notably, the interaction between SerRS and SIRT2 can also be induced by NAD^+^ (Fig. 2f, g), albeit to a weaker extant than ADPR. Considering the relatively high concentration of free cytosolic NAD^+^ in mammalian cells^5^ (∼40 to 70 μM), we propose that, even in the absence of ADPR, SerRS can act as a substrate-like competitive inhibitor to modulate SIRT2 activity and specificity, allowing only those with high abundance and/or high affinity for SIRT2 to be deacetylated. Therefore, under normal conditions, when NAD^+^/ADPR ratios are high, SerRS may safeguard SIRT2’s specificity within the acetylome. However, under stress conditions, when ADPR levels rise rapidly, SerRS would form a stable complex with SIRT2 to inhibit each other as a stress response to protect the cell. This response could be further enhanced with a decrease in tRNA levels. Indeed, under a variety of stress conditions, tRNAs are cleaved by angiogenin^46^. Depletion of tRNA can also result from tRNA retrograde transport, which is also increased upon oxidative stress^47^. Future studies to identify biological conditions that strengthen the SerRS/SIRT2 interaction are likely to provide new insights into cellular stress responses.

Although SIRT1 and SIRT2 share a high degree of sequence similarity in the catalytic domain (68%, Extended Data Fig. 9), SerRS can interact with SIRT2 but not SIRT1^14^. Our structure of the SerRS/SIRT2 complex and the subsequent mutagenesis and functional studies provide the explanations for the selectivity. A key residue for the SerRS/SIRT2 interaction, F244, is only found in SIRT2, but not in other members of the sirtuin family (Fig. 5f; Extended Data Fig. 9). Furthermore, previous studies have shown that sirtuins recognize their substrates in a context-dependent manner, lacking a clear consensus sequence^48^. Analyzing the sequence context around the K414 site in the substrate-mimetic loop of SerRS, we found that Q412 at the −2 position and M416 at the +2 positions are present exclusively in SIRT2 substrates but not in those of other sirtuins. Indeed, mutations in Q412 and M416 both dramatically attenuated the SerRS/SIRT2 interaction (Fig. 5d), further supporting the specificity of SerRS for SIRT2 over other sirtuins.

## Supporting information

Report of 9DPI

Report of 9DPD

## Author Contributions

Q.Z. and X.-L.Y. designed the studies. Q.Z. and H.Z. designed and performed the biochemical and cell-based experiments. M.H. performed the initial cryo-EM studies. J.Y. collected and processed the cryo-EM data and determined the structures. S.L. conducted the HDX analysis. Q.Z., H.Z., X.-L.Y., J.Y., and G.C.L. analyzed and interpreted the data. Q.Z., H.Z., and X.-L.Y. drafted and revised the manuscript with contributions from all authors.

## Acknowledgments

This work was funded by National Institutes of Health R35 GM139627 grant to X-L Y. Specifically, we also want to thank Elizabeth Billings and Bill Webb (Center for Metabolomics and Mass Spectrometry at The Scripps Research Institute) for their contribution in ESI triple quad mass spectrometry analysis.

---

**Extended Data Fig. 1.**
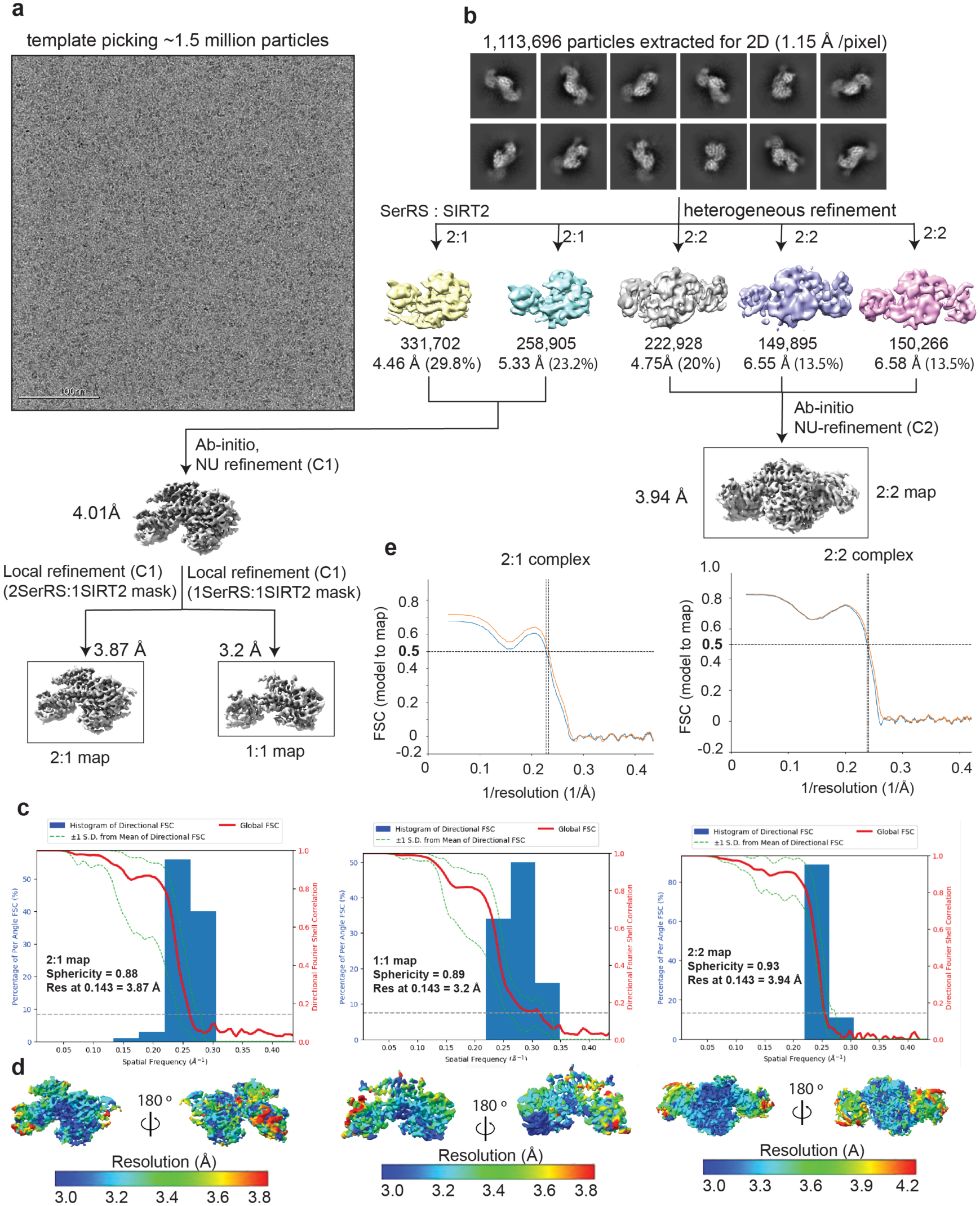
Cryo-EM structure determination of SerRS/SIRT2 complex. (a) Representative micrograph of cryo-EM data collection. (b) Workflow of cryo-EM data processing using CryoSparc software^49^. The final 3D reconstruction 2:1 and 2:2 maps were used for model building and refinement. (c) Three-Dimensional Fourier Shell Correlation (3DFSC)^50^ of the 2:1, 1:1, and 2:2 SerRS/SIRT2 reconstructions reporting a global resolution at 3.7 Å, 3.2 Å, and 3.94 Å respectively. (d) Final reconstructions in (c) are filtered and colored by local resolutions estimated from CryoSparc. (e) Model-to-map FSC plots for the 2:1 and 2:2 maps from Phenix validation.

**Extended Data Fig. 2.**
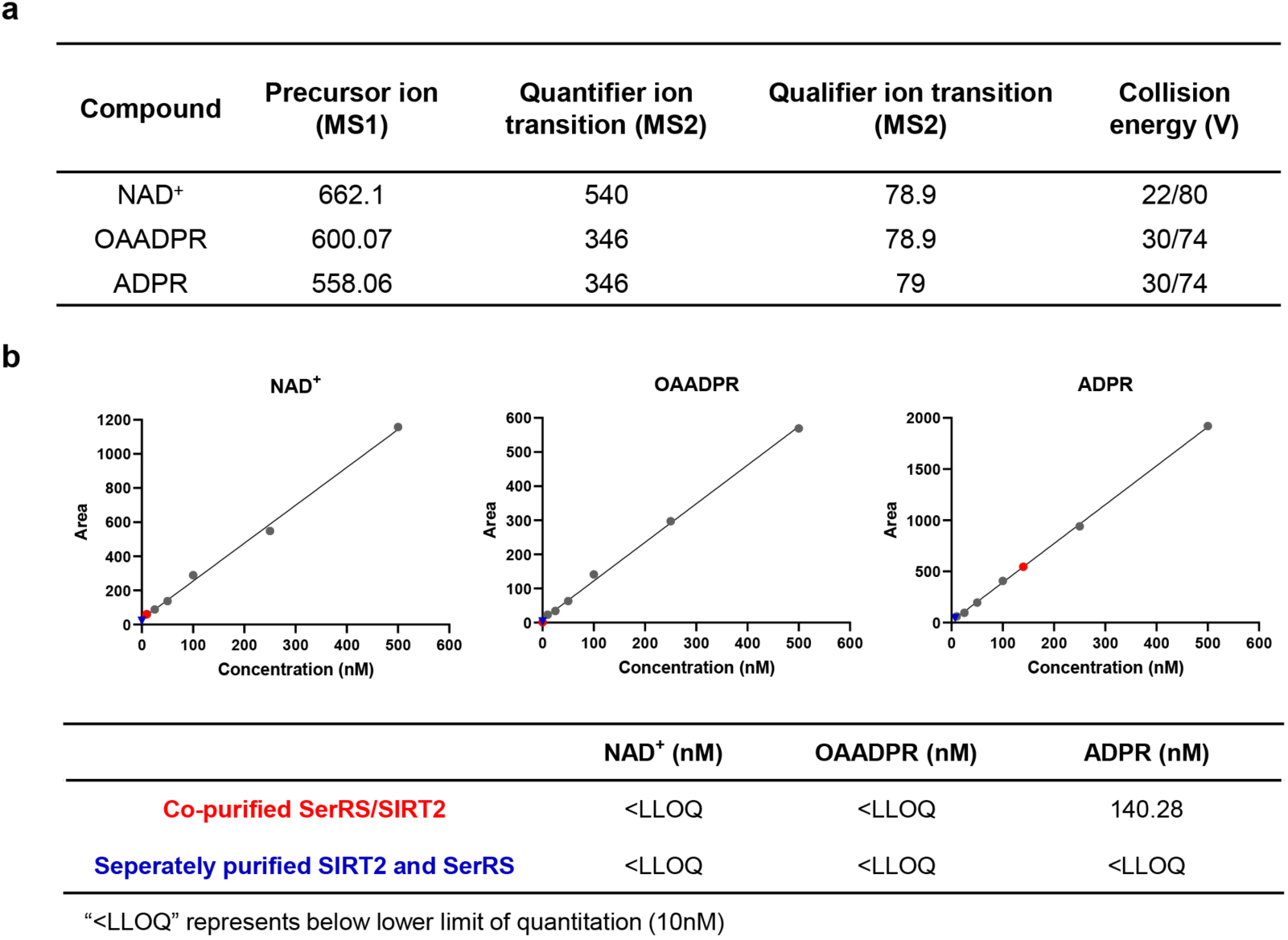
ADPR but not NAD^+^ or OAADPR was detected in the co-purified SerRS/SIRT2 complex. (a) ESI Triple Quad Mass Spectrometry analysis parameters for NAD^+^, OAADPR, and ADPR. (b) The standard curve of NAD^+^, OAADPR, and ADPR (6 points ranging from 10-500nM are shown here) established through ESI Triple Quad Mass Spectrometry for quantifying the concentrations of ligands extracted from co-purified SerRS/SIRT2 complex (red dots) and individually purified SIRT2 and SerRS proteins (blue triangles).

**Extended Data Fig. 3.**
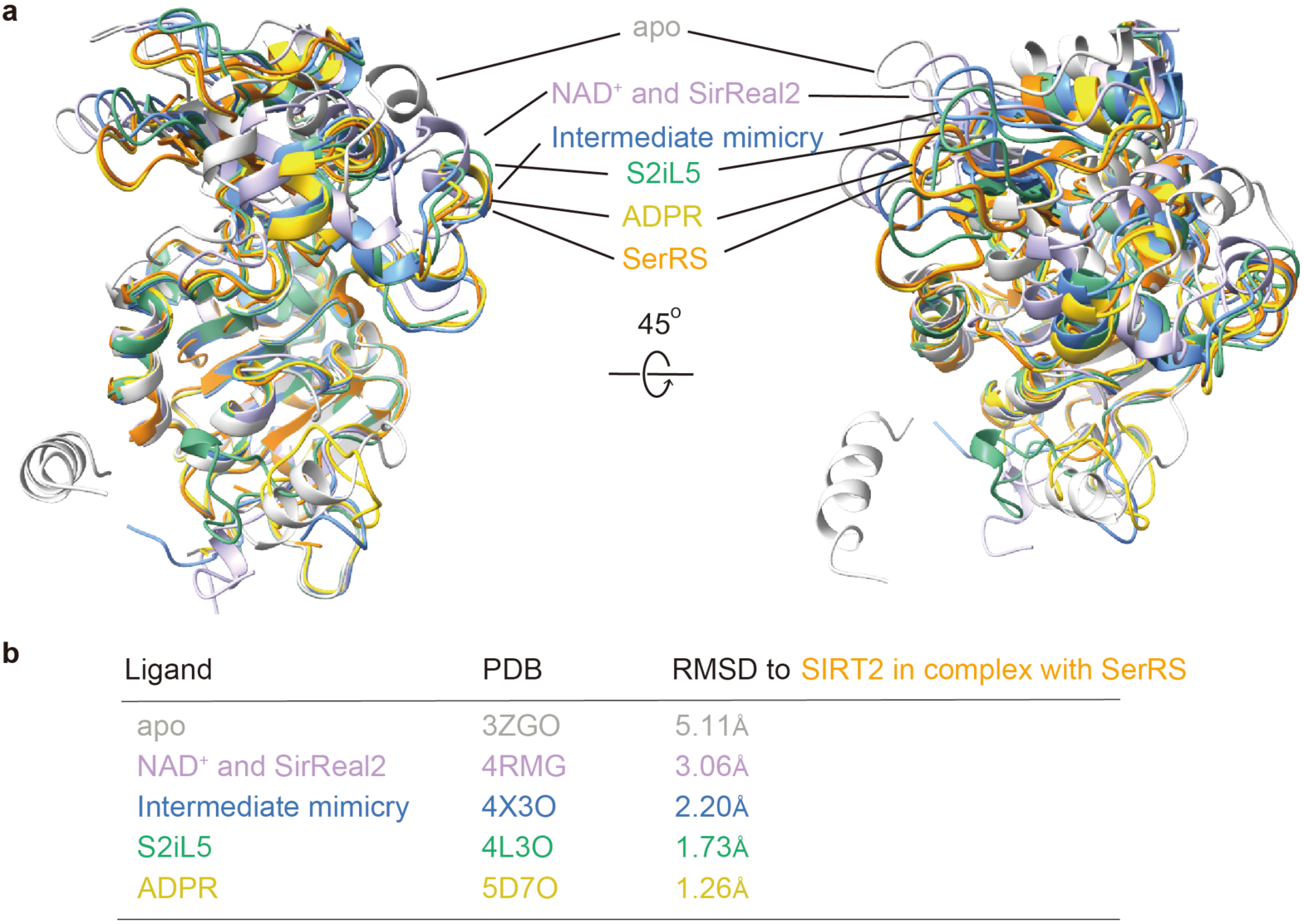
Structural comparison of the SerRS/SIRT2 complex and other SIRT2 complexes. (a) Superposition of SIRT2 structure from the SerRS/SIRT2 complex with apo SIRT2 and different ligands bound SIRT2. (b) RMSD values between SIRT2 in SerRS/SIRT2 complex and other ligand bound SIRT2 structures.

**Extended Data Fig. 4.**
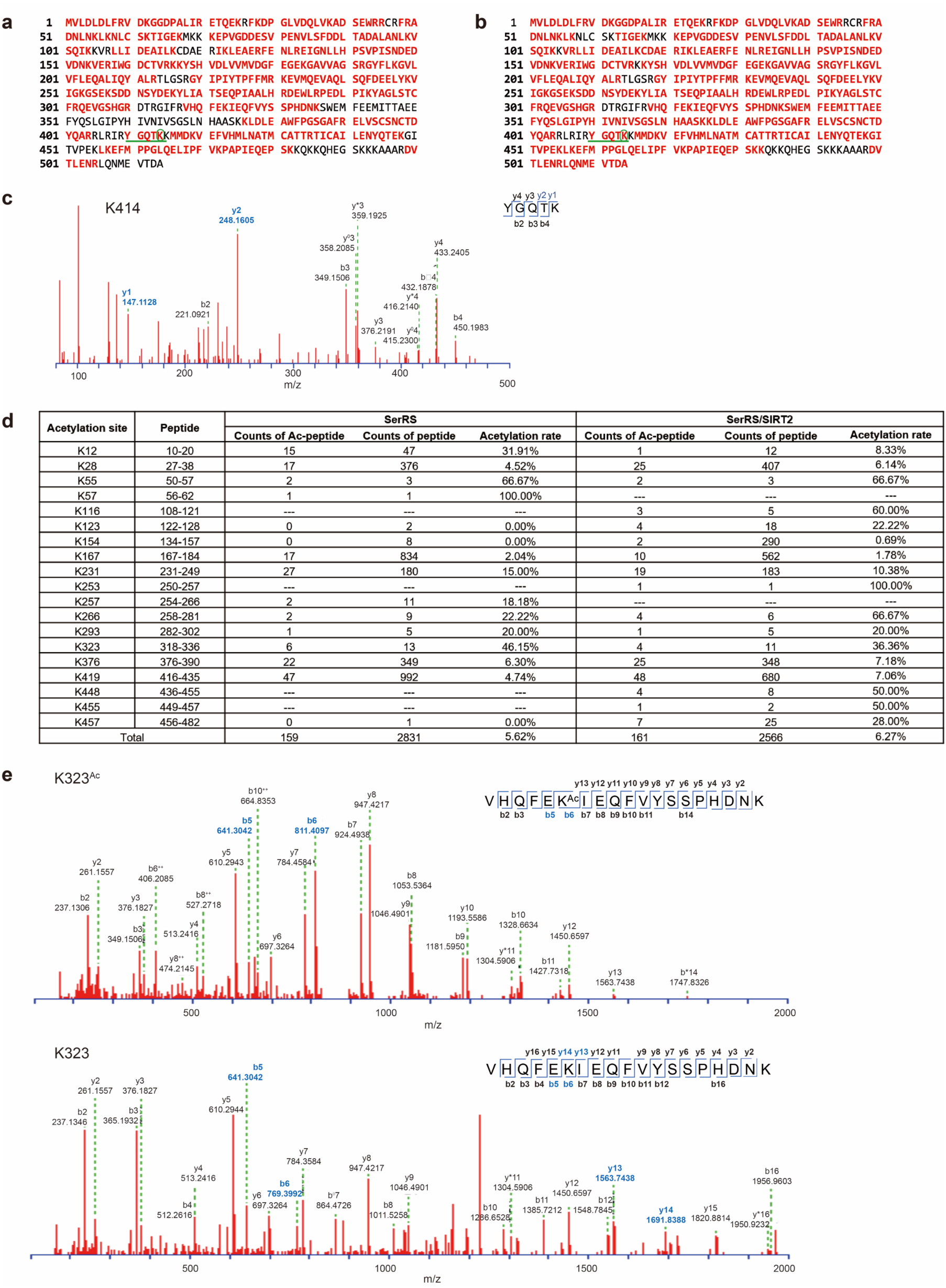
Trypsin digestion and mass spectrometry analysis indicating SerRS K414 is not acetylated. (a and b) Coverages of peptides (red) for the free recombinant SerRS (a) and SerRS in the co-purified SIRT2/SerRS complex (b). The K414-containing peptide (^410^YGQTK^4^^14^) is underlined, and the K414 residue is highlighted in a circle. (c) Mass spectrometry analysis of the K414-containing peptide (^410^YGQTK^4^^14^) shows the absence of K414 acetylation. (d) List of lysine acetylation sites detected by the mass spectrometry analysis. (e) An example of lysine acetylation (K323) detected by the mass spectrometry analysis.

**Extended Data Fig. 5.**
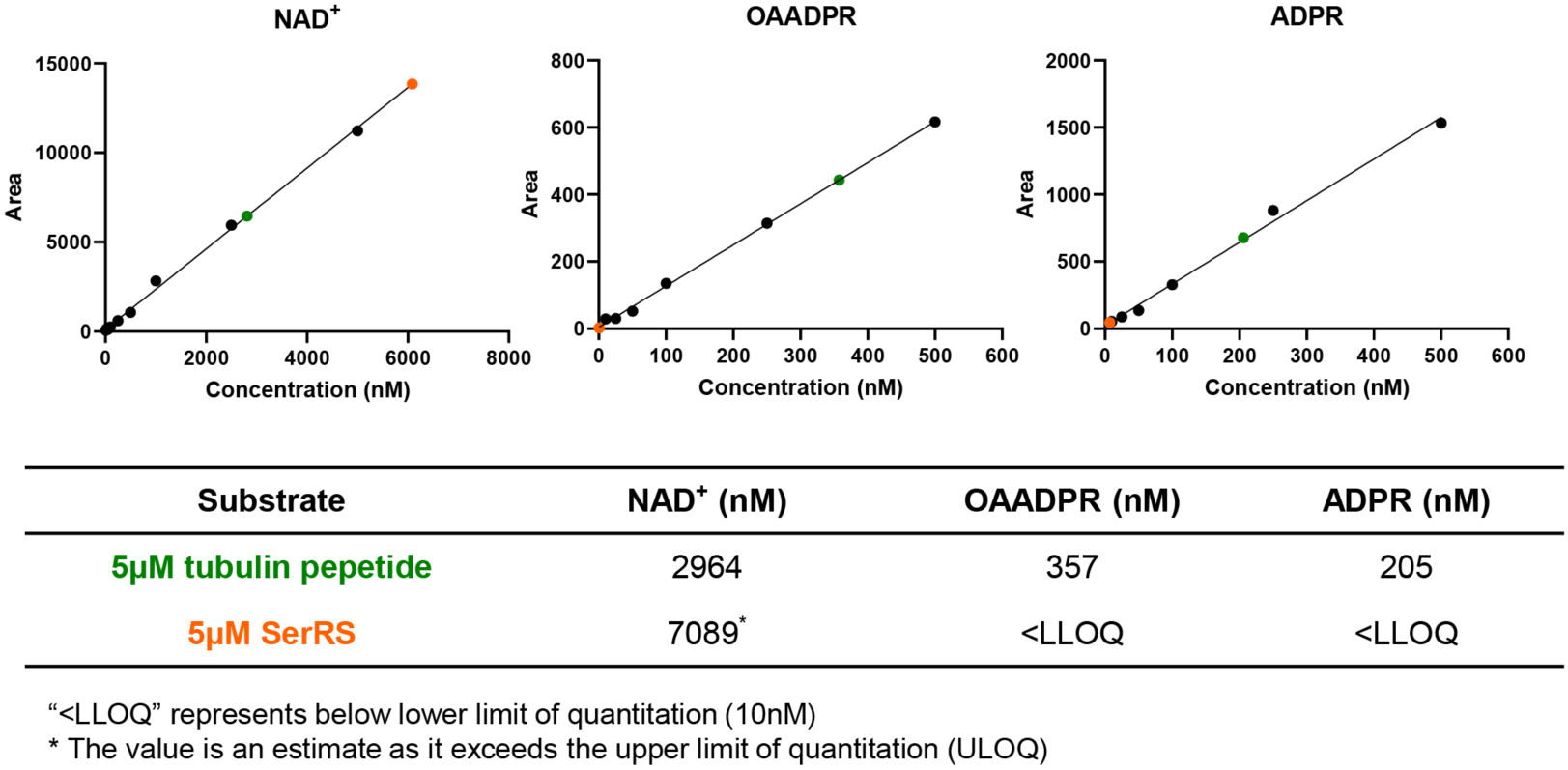
Absence of SIRT2 deacetylation activity with SerRS. The concentration of ligands extracted from SIRT2 deacetylation reactions using acetyl-α-tubulin peptide (green dots) or SerRS (orange dots) as the substrate is quantified and indicated on the standard curve of NAD^+^, OAADPR, and ADPR.

**Extended Data Fig. 6.**
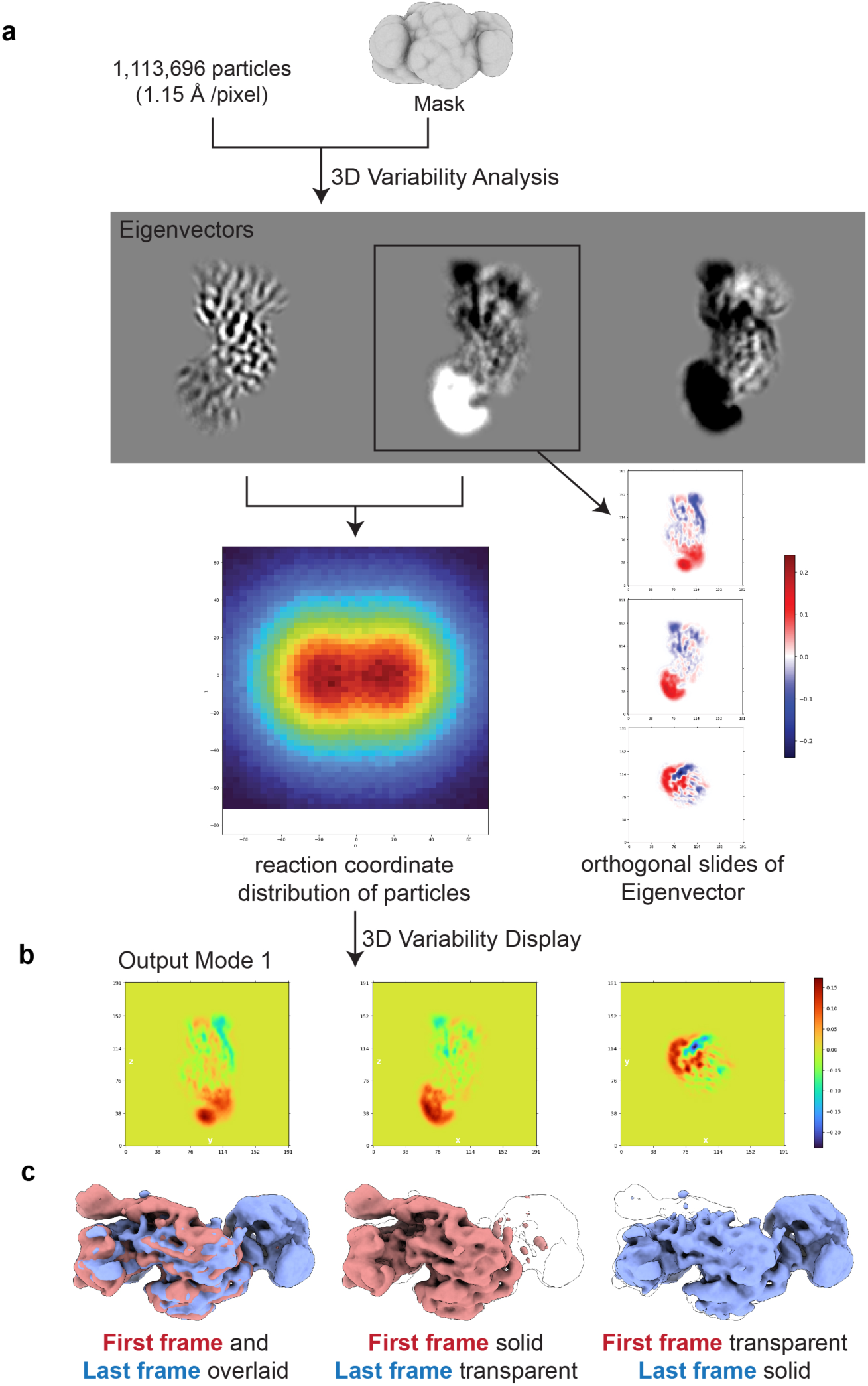
3D Variability Analysis (3DVA) of SerRS/SIRT2 complexes. (a) Schematic workflow of 3D Variability Analysis (3DVA) performed using CryoSparc. (b) 3D variability display of a representative volume series, showing dynamic conformational changes observed within the SerRS and SIRT2 complexes. (c) Overlay of the first and last frames from the representative volume series displayed in (b).

**Extended Data Fig. 7.**
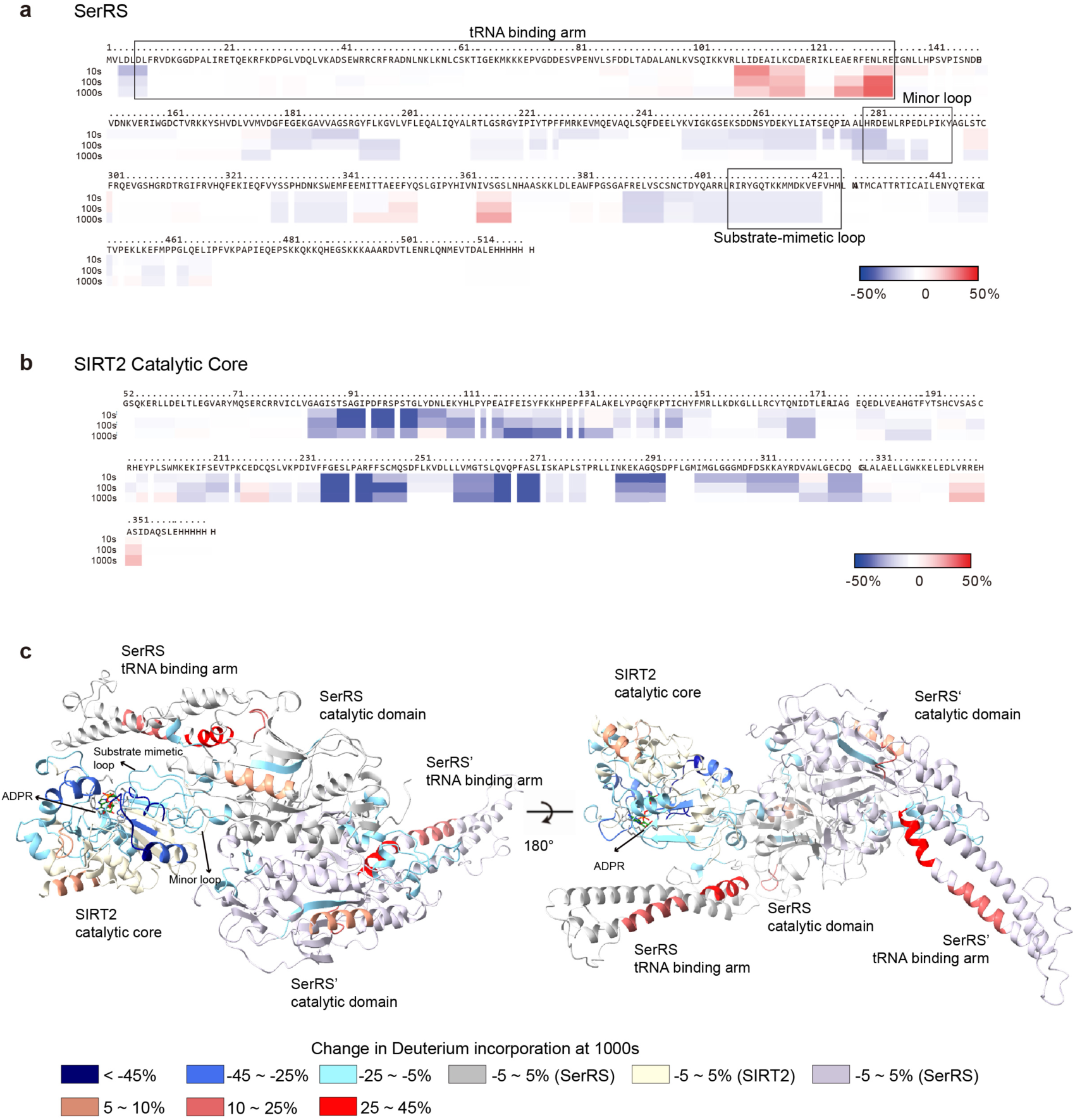
Hydrogen-Deuterium Exchange (HDX)-Mass Spectrometry analysis of SerRS/SIRT2 interaction. (a) Differential deuterium incorporation of SerRS upon binding with SIRT2_52-356_-His_6X_ over the time course of 10-1000 seconds, mapped onto SerRS sequence. Blue regions indicate areas with increased protection upon SIRT2 binding, while red regions indicate areas with increased exposure upon SIRT2_52-356_-His_6X_ binding. (b) Differential deuterium incorporation of SIRT2_52-356_-His_6X_ caused by SerRS binding over 10-1000 seconds, mapped onto the SIRT2_52-356_-His_6X_ sequence. (c) Differential deuterium incorporations at 1000 seconds from (a) and (b), mapped onto the cryo-EM structure of SerRS/SIRT2 2:1 complex.

**Extended Data Fig. 8.**
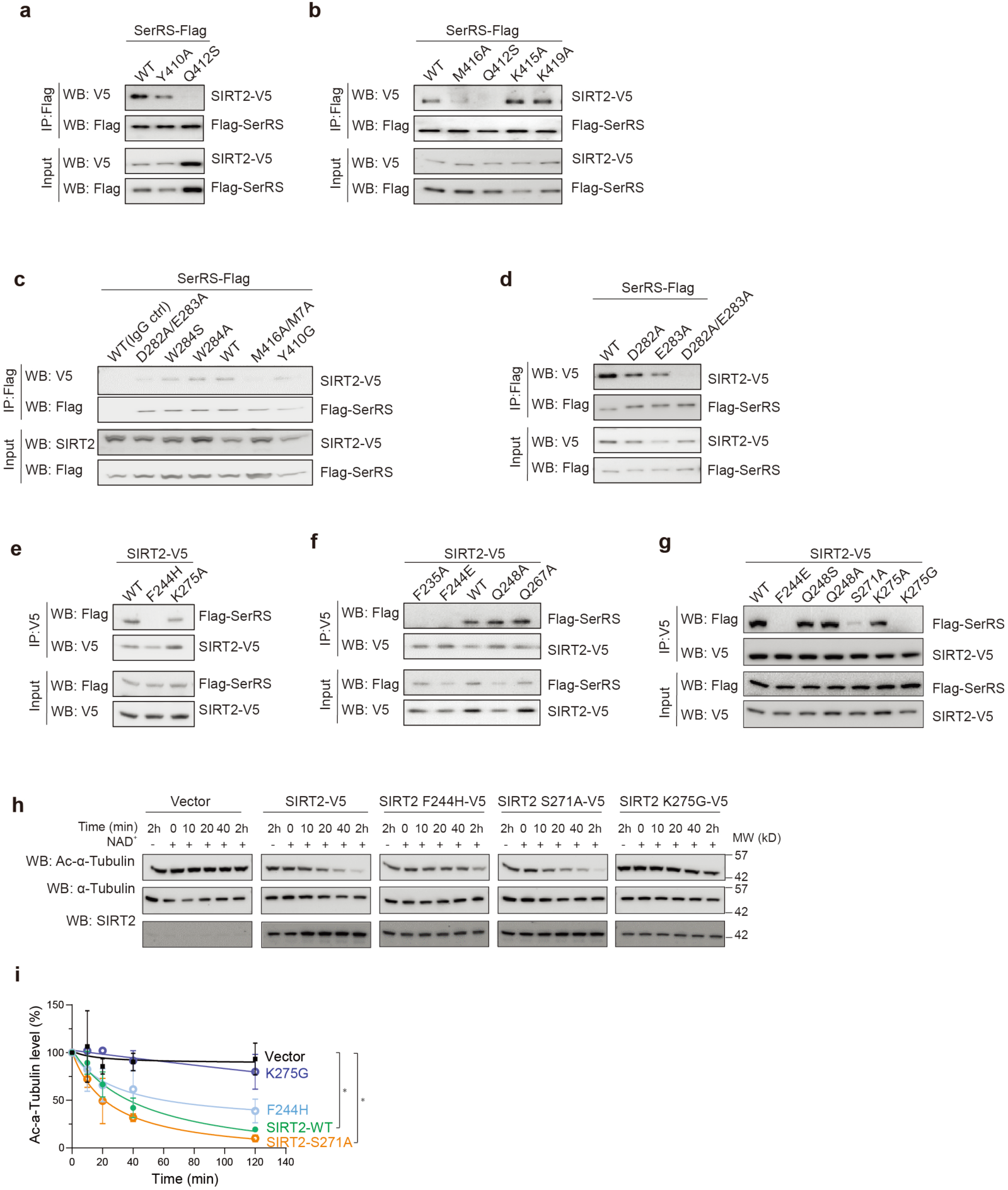
Identifying key residues for SerRS/SIRT2 interaction. (a-g) Mutagenesis and Co-immunoprecipitation assays identifying key residues from both SerRS (a-d) and SIRT2 (e-g) involved in the SerRS/SIRT2 interaction (related to Fig. 4d, e, f). (h) time course cell-based deacetylation assays using HEK293 cell lysates with overexpression of SIRT2 or SIRT2 mutants. (i) Quantification analysis of Ac-α-Tubulin levels from (h). Ac-α-Tubulin levels at each time point were normalized to that at 0 min time point, which was defined as 100% for each reaction. Error bars represent ±SME for duplicate experiments. ** *p* < 0.01, two-way ANOVA test, n=2.

**Extended Data Fig. 9.**
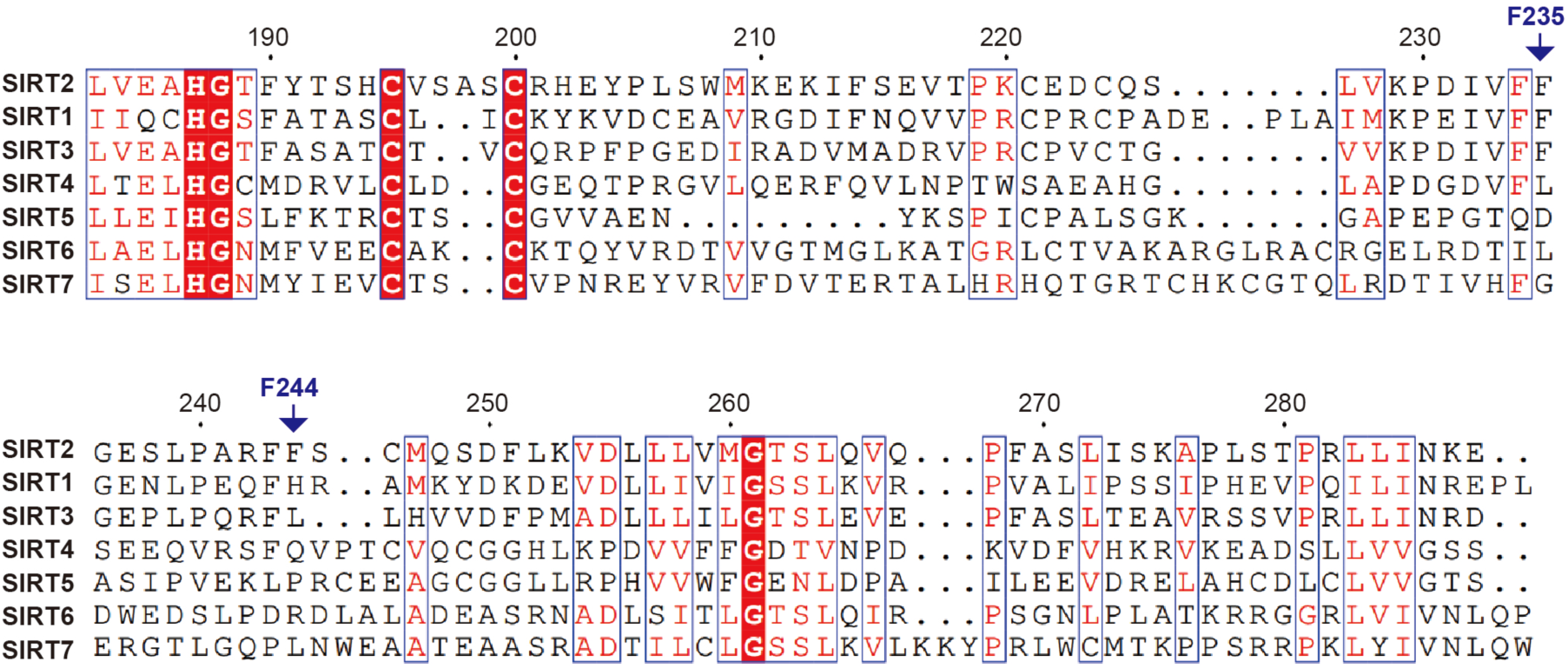
Multiple sequence alignment of Sirtuin family proteins. Multiple sequence alignment of Sirtuin family proteins is presented. F235 and F244 in SIRT2 are labeled.

**Supplementary Table 1**

**Cryo-EM data collection, refinement, and validation**

**Supplementary Video**

**3D Variability Analysis (3DVA) of SerRS and SIRT2 complexes.**

This video illustrates the dynamic conformational changes in the SerRS and SIRT2 complexes, as revealed by 3D Variability Analysis (3DVA) using CryoSparc. The video contains 20 frames from a volume series, capturing conformational variability within the complex. Notably, the tRNA binding arm (colored in magenta) of SerRS exhibits an anti-correlated movement with the neighboring SIRT2 binding (colored in blue).

## Material and Method

### Antibodies

Custom-made mouse monoclonal and rabbit polyclonal anti-human SerRS antibodies were raised against human recombinant SerRS and affinity purified. The following commercial antibodies were used: Anti-Flag rabbit and mouse (#14793 and #8146), Anti-V5 rabbit (#13202), anti-α-tubulin (#3873), and anti-acetylated-α-tubulin (K40Ac) (#5335), all from Cell Signaling Technology (Danvers, MA, USA). Anti-SIRT2 antibody (#12650) for western blot was purchased from Cell Signaling.

### Protein expression and purification

For protein expression, N-terminal His-SUMO-tagged or tag-free human SerRS construct was generated by subcloning human SerRS gene into a modified pET-28a vector with a SUMO gene inserted at the 5’ of the MSC site. C-terminal His-tagged constructs of human SerRS, SIRT2 and SIRT2_52-356_ (catalytical core region of G52-S356 aa) were subcloned into pET-20b (+) vector (Novagen, Darmstadt, Germany). Recombinant proteins were overexpressed or co-expressed (for the SerRS/SIRT2-His_6X_ complex) in *E. coli* BL21(DE3) cells. For protein purification, the cells were lysed by M-110L Microfuidizer Processor (Microfluidics) in 25mM HEPES-Na (pH7.5), 500mM NaCl, and 20mM imidazole (binding buffer). Cell lysates were centrifuged at 28,000 rpm for 40 min, and the soluble fractions were applied to a Ni-NTA column (QIAGEN). After washing the Ni-NTA resin with the binding buffer containing 10-40mM imidazole, bound proteins were eluted with the same buffer containing 250mM imidazole, The eluted proteins from Ni-NTA were further purified by either a HiTrap Q HP anion exchange column (GE Healthcare, Pittsburgh, PA, USA) for SIRT2-His_6X_ and SIRT2_52-356_-His_6X_, or a HiTrap Heparin HP affinity column (GE Healthcare) for SerRS and SerRS/SIRT2-His_6X_ complex, using a NaCl gradient from 50 to 500 mM. Final protein purification was achieved by size exclusion chromatography using a HiLoad 16/600 Superdex 200 pg column (GE Healthcare) in 25mM HEPES-Na (pH7.5) and 150mM NaCl. Tag-free SerRS was generated by overnight incubation of the Ni-NTA resin-bound His_6X_-SUMO-SerRS protein with homemade Ulp1 enzyme to cleave the His_6X_-SUMO tag. The resulting tag-free SerRS was collected the following day in the flow through and further purified by heparin affinity and size exclusion chromatography.

### Sample preparation of cryo-EM

Purified SerRS/SIRT2 complex (0.5mg/mL) was supplemented with 0.05% Lauryl Maltose Neopentyl Glycol (LMNG) prior to cryo-freezing. The sample (4 µL at 0.5mg/mL) was applied onto 300 mesh R1.2/1.3 UltraAuFoil Holey Gold grids (Quantifoil)that has been plasma cleaned for 15 seconds using a Pelco glow discharge cleaning system (TED PELLA, INC.) with atmospheric gases at 15 mA. The sample (4 uL at 0.1mg/mL) was also applied onto 300 mesh R1.2/1.3 UltraAuFoil Holey Gold grids (Quantifoil) that were coated with a single layer of graphene^51^. The graphene grids were treated with UV ozone for 4 mins before usage. Both samples were plunge-frozen using a Thermo Fisher Vitrobot Mark IV at 4 °C, 5 s, 100% humidity, and a blot force of 0 using Whatman #1 blotting paper.

### Cryo-EM data acquisition, processing and reconstruction

Cryo-EM data were collected on a Thermo-Fisher Talos Arctica transmission electron microscope operating at 200 keV using parallel illumination conditions^52, 53^. Micrographs were acquired with a Gatan K2 Summit direct electron detector with a total electron exposure of 50e^−^/Å^2^ as a 98-frame dose-fractionated movie during a 9800-ms exposure time. The Leginon data collection software was used to collect 2126 micrographs on unmodified Holey Gold grids and 1394 micrographs on graphene coated Gold grids at 36,000 nominal magnification (1.15 Å/pixel) with a nominal defocus ranging from −.7 µm to −1.2 µm. Stage movement was used to target the center of four 1.2-µm holes for focusing, and image shift was used to acquire high magnification images. The Appion image processing wrapper^54^ was used to run MotionCor2 for micrograph frame alignment and dose-weighting in real-time during data collection^55^. All subsequent image processing was done in Cryosparc for the combined 3520 micrographs^56^. The CTF parameters were estimated using Patch CTF estimation (multi). Representative views of 2D classes were used as templates for template-based particles picking in Cryosparc and 5,869,493 picked particle coordinates were extracted at 1.15 Å/pixel from the motion-corrected and dose-weighted micrographs with a box size of 192 pixels. The extracted particles were classified into 400 classes of 2D averages using default parameters. 2D classes (1,113,696 particles) displaying high-resolution secondary features were selected and used to firstly generate a reference-free 3D Ab-initio model, and then subjected to 5 classes of 3D classification using the heterogeneous refinement in Cryosparc with default parameters. In one processing strategy, class 1 (331,702 particles) and class 2 (222,928 particles), which displayed high-resolution structural details, were selected for 3D NU-Refinement, which resulted in a reconstruction of ∼4.01 Å. The map was further refined using Local Refinement by masking the region of either the SerRS dimer and SIRT2, or the SerRS subunit and SIRT2, which resulted in final reconstructions of reported resolutions at 3.87 Å and 3.2 Å respectively. In an alternative processing strategy, class 2 (222,928 particles), class 4 (149,895 particles), and class 5 (150,266 particles) containing SerRS dimer bound with 2 SIRT2 molecules were selected for 3D NU-Refinement with C2 symmetry, which resulted in a reconstruction of 3.94 Å. All the resolutions were estimated using the 3DFSC server (https://3dfsc.salk.edu)^50^ using a Fourier Shell Correlation (FSC) cutoff at 0.143 (Extended Data Fig. 1). The crystal structure of SerRS dimer (PDB 4L87) and the crystal structure of SIRT2 (PDB 4RMI) were initially docked into the reconstructed density maps using UCSF Chimera. Real-space structural refinement and manual model building were performed with Phenix^57^ and Coot^58^, respectively. Phenix real-space refinement includes global minimization, rigid body, local grid search, atomic displacement parameters, and morphing for the first cycle, and was run for 100 iterations, 5 macro cycles, with a target bonds RMSD of 0.01 and a target angles RMSD of 1.0. The refinement settings also include the secondary structure restraints, Ramachandran restraints. Figures of the structures were prepared using UCSF Chimera, and UCSF ChimeraX ^59, 60^. Supplementary Table 1 includes data collection, refinement, and validation statistics ^61^.

### 3D Variability Analysis

A total of 1,113,696 high-quality particles were selected for 3D Variability Analysis (3DVA) using CryoSparc. The particles were first subjected to non-uniform refinement to generate a consensus 3D reconstruction, which served as the reference model for the 3DVA. The particle stack and output mask from the non-uniform refinement job were used as inputs for the 3D Variability job. For 3DVA, the filter resolution was set to 4.5 Å, while all other parameters were kept at their default settings. The variability was analyzed using principal component analysis (PCA), and three volume series were generated, with each series containing 20 volume frames. The 3D variability was visualized through the 3D Var Display job in CryoSparc using default visualization parameters.

### Cell culture and constructs

For cell-based assays, human full-length SIRT2 was cloned into pCDNA6-V5/His-C vector (Life Technologies), while human SerRS was cloned into pFlag-CMV-2 vector (Sigma-Aldrich). These constructs were used to generate the SerRS and SIRT2 mutants tested in this study. All cells were cultured in a humidified incubator at 37°C with 5% CO_2_. Human HEK293 cell lines were purchased from the American Type Culture Collection (ATCC, Manassas, VA, USA) and the human HEK293AD cell line was purchased from Agilent (Agilent, Santa Clara, CA, USA). Cells were maintained in Dulbecco’s Modified Eagle Medium (ThermoFisher Scientific, Grand Island, NY, USA) supplemented with 10% fetal bovine serum (Omega Scientific, Tarzana, CA, USA) to a final concentration of 10% and 1% Penicillin-Streptomycin (ThermoFisher Scientific). Transient transfections were performed by using Lipofectamine 3000 (ThermoFisher Scientific).

### Coimmunoprecipitation assays and Western blot analysis

HEK293AD cells were washed with PBS pH7.4 and lysed on ice for 5min using cell lysis buffer (20mM Tris-HCl pH7.5, 150mM NaCl, 1mM EDTA, 1mM EGTA, 1% NP40, 0.5% sodium deoxycholate, and freshly added protease inhibitor cocktail). The indicated antibodies were incubated with 20µL protein-G-conjugated agarose beads (Invitrogen) at 4°C for 2 hours. The supernatants of the cell lysates were collected and incubated with the antibodies coated beads overnight at 4°C. The beads were then washed three times with ice-cold wash buffer (20mM Tris-HCl pH7.5, 150mM NaCl, 1mM EDTA, 1mM EGTA, 0.1% Triton X-100, and freshly added protease inhibitor cocktail). After the final wash, the beads were incubated in SDS loading buffer for 5min, followed by SDS-PAGE and Western blotting analysis with the indicated antibodies.

### Biolayer interferometry

SerRS/SIRT2 interactions were analyzed using Octet RED96 system (Sartorius) at 25 °C with 1× kinetic buffer purchased from Sartorius. SIRT2-His_6X_ was immobilized on HIS1K biosensor tips for 5 minutes, equilibrated in buffer containing 0.5mM NAD^+^,0.5mM ADPR or no ligand to establish a baseline. The SIRT2 coated tips were then dipped into 180 nM SerRS for 5 minutes association, followed by 5 minutes dissociation in ligand-only buffer. Data were processed by subtracting the reference (SerRS alone without ligand) and aligning the association and disassociation curves.

### ESI triple quad mass spectrometry analysis

Calibration standards were prepared by diluting stocks of ADPR, Acetyl-ADPR and NAD to the appropriate calibration levels. To probe whether SerRS acts as a substrate for SIRT2, the SIRT2 deacetylation assay was performed at room temperature for 3h in 100μL of 1μM SIRT2, 10μM NAD^+^, and 5μM acetyl-α-tubulin peptide or SerRS. To facilitate the release of enzyme-bound ligands, 100μL of 10μM co-purified SerRS/SIRT2 complex or the individually purified proteins were denatured by boiling at 95℃ for 2 mins. All samples were followed by the addition of 400μL of ice-cold MeOH and incubation at −20℃ for 1h. After centrifugation at 13,000rpm for 15min at 4℃, the supernatants were evaporated overnight in a SpeedVac at 10℃. Each sample was added with 100μL of 50:50 MeOH:H_2_O, sonicated for 10min and centrifuged at 12,700rpm for 10min, then transferred to autosampler vials for analysis. The samples were run on an Agilent 6495 triple quadrupole mass spectrometer coupled to an Agilent 1290 UPLC stack. The column used was a Waters BEH-amide 2.1×150mm. The mobile phase composition was as follows: mobile phase A: 10mM NH_4_OAc, 5μM deactivator, pH=9, and mobile phase B: 90:10 ACN:H_2_O, 10mM NH_4_OAc, 5μM deactivator, pH=9. The flowrate was set to 250µL/min and the sample injection volume was 5 µL. The gradient was: T0: 0/100, T1: 0/100, T10: 65/35 T13: off, with a 5min re-equilibration step. Operating in negative ion mode, the source conditions were as follows: Capillary voltage: 2500V, Gas temp: 120℃, Gas flow: 11L/min, Nebulizer pressure: 35psi. Sheath gas temp: 350℃, sheath gas flow: 12L/min. Nozzle voltage was set to 1500V. Data was processed using Agilent Quantitative Analysis software. Calibration curves were prepared in concentrations ranging from 10nM to 5000nM, with the lower limit of quantitation (LLOQ) set at 10nM and the upper limit of quantitation (ULOQ) set at 5000nM.

### Trypsin digestion and mass spectrometry analysis

SDS-PAGE gel bands of SerRS (from recombinant SerRS alone or SerRS/SIRT2 complex) were cut out, destained, reduced (10 mM DTT), alkylated (55mM iodoacetamide), and digested with 10ul of trypsin overnight before nano-LC-MS/MS analysis. The analysis was performed using Thermo Scientific Easy nano LC 1200 coupled with Thermo Scientific Q Exactive Plus using Nanospray Flex ion source. 8μL of digested peptides were separated by reverse-phase chromatography on a 14cm length x 75mm inner diameter nano electrospray capillary column, packed in-house with Phenomenex Aqua 3 µm C18 125 Å. Mobile phase A was water including 0.1% formic acid, and mobile phase B was 80% acetonitrile plus 0.1% formic acid. The elution gradient started at 2% B for 5min, ramped up to 25% B for 100mins, 40%B for 20 mins, 95%B for 1 min and held at 95% B for 14 mins. Data acquisition was performed using Xcalibur (version 4.3). One MS scan of m/z 400-2000 was followed by 10 MS/MS scans on the most abundant ions with application of the dynamic exclusion of 10sec. MS/MS Raw data was searched against the Custom sequence database which contains the sequence of SerRS and the NCBI Homo sapiens (human) database using Mascot (Version 2.8.0, Matrix Science). Mascot searches were conducted using a peptide mass tolerance of 10ppm, a fragment ion mass tolerance of 0.01 Da, fixed modifications of carbamidomethyl (C), variable modifications of Acetylation (K), Myristoylation (K) and Palmitoylation (K), an enzyme of trypsin and a maximum missed cleavage of one. Identification was done with a false discovery rate (FDR) of <2% as determined by using a decoy database search.

### Fluorometric SIRT2 deacetylase activity assay

SIRT2 deacetylase activity was measured using Cyclex SIRT2 Deacetylase Fluorometric Assay Kit (CycLex, Nagano, Japan). The reaction buffer contains 50mM Tris-HCl (pH 8.8), 0.5mM DTT, 0.25mAu/mL Lysylendopeptidase, 1 µM Trichostatin A, 0.8mM NAD^+^, 20uM Fluoro-Substrate peptide, 1µM recombinant SIRT2 and recombinant SerRS at different ratio indicated in Fig. 3b. For assessing the impact of ADPR (Fig. 3c), 0.25µM recombinant SIRT2, 1µM SerRS or BSA, 4µM Fluoro-Substrate peptide, 40µM NAD^+^, and 0, 200, or 400 µM of ADPR were used.

### Cell-based SIRT2 deacetylase activity assay

HEK293T or HEK293AD cells were seeded in 6 well plate and allowed to adhere overnight. The following treatments were then applied: 5 µM SIRT2 inhibitor AGK2, SerRS knockdown by siRNA (Santa Cruz), SIRT2 overexpression, or combined overexpression of both SIRT2 and SerRS. Tree days post-treatment, 5 µM Tubastin A was applied for 6 hours to inhibit HDAC6-mediated deacetylation. cells were then washed with cold PBS (pH 7.4) and lysed on ice with lysis buffer containing freshly added protease inhibitor cocktail (Thermo scientific) for 5 minutes. All samples were applied to SDS-PAGE followed by Western blot analysis using the indicated antibodies. For the time course investigation, HEK293AD cells were harvested 48 hours post-transfection of SerRS, SIRT2, or both. cells were washed and lysed within 150ul/well cell lysis buffer (freshly added 8µM Tubastin A and protease inhibitor cocktail) on ice for 5minutes. Supernatants were collected and incubated at room temperature for 40 minutes to stabilize Ac-α-Tubulin levels. For each reaction, 100µL supernatant was used in a 400 µL reaction system containing 50mM Tris-HCl (pH 9.0), 4mM MgCl_2_, 0.2mM DTT, 8µM Tubastin A, and 2mM NAD^+^. At each time points, 40uL reaction mixtures were sampled for Western blotting analysis with indicated antibodies.

### Hydrogen deuterium exchange mass spectrometry

To initiated hydrogen deuterium exchange (HDX) reactions, 3 μl of each protein stock solution (SerRS, SIRT2 or co-purified SerRS/SIRT2 protein) was mixed with 9 μl of D_2_O buffer (8.3 mM Tris, 150 mM NaCl in D_2_O, pH 7.6), respectively, and incubated for 10, 100, 1000 sec at 0°C. At the designated times, HDX reactions were quenched by adding 18 μl of ice old quench buffer (0.08 M GuHCl, 0.8% formic acid, 16.6% glycerol, pH 2.4) and samples were immediately frozen at –80°C. Control samples, both non-deuterated and fully deuterated, were also prepared following the previously established method^62^. The 30 μl quenched samples were thawed at 4°C and immediately passed over the immobilized pepsin column (1 × 20 mm, 40 mg/ml porcine pepsin (Sigma). The digested peptides were collected on a trap column (Michrom MAGIC C18AQ, 0.2 × 2 mm) for desalting and separated with a Magic C18 column (Michrom, 0.2 × 50 mm, 3 μm, 200 Å) by a 30 min linear acetonitrile gradient of 6.4–38.4%. The accurate peptide mass measurement was performed with an OrbiTrap Elite mass spectrometer (Thermo Fisher Scientific), which was setup for optimal HDX performance with minimum back-exchange rate^63^. Data acquisition was completed in either data-dependent MS/MS or MS1 profile mode and peptide identification was done using Proteom Discoverer software (Thermo Fisher Scientific). A total of 275 peptides with 98.40 % of SIRT2 sequence coverage and 273 peptides with 96.16% of SerRS sequence coverage were employed to track deuterium exchange. Deuterium incorporation levels of each peptide at each time-points were determined by HDXaminer (Sierra Analytics Inc) with back-exchange correction use above control samples.

### In vitro transcription of tRNA

Template DNAs for human tRNA^Ser(CGA)^ and tRNA^Ser(GCT)^ were amplified through PCR. Subsequent *in vitro* transcription was performed using T7 RNA polymerase (HiScribe T7 High yield RNA Synthesis kit, NEB). The reaction was performed at 37°C for overnight. The transcription product was then subjected to 10% Urea-PAGE (Life Tech) to assess purity and separate it from proteins and NTPs. The specific tRNA transcripts bands were excised, dissolved in 0.3M NaOAc (pH 5.0) and 1mM EDTA, followed by ethanol precipitation of the supernatant. Prior to use in pull-down assay, tRNA^Ser(CGA)^ and tRNA^Ser(GCT)^ were mixed at a 1:1 ratio, incubated at 95°C for 5 min, and subsequently refolded at 60°C in the presence of 5 mM MgCl_2_ and gradually cooling down to room temperature.

### Ni-NTA Pull down assay

The *in vitro* Ni-NTA pull down assay was performed in the buffer with 20 mM Hepes (pH 7.5), 150mM NaCl, 50 mM imidazole (pH 8.0), 10% glycerol and 0.5% NP-40. 3 μg Recombinant SIRT2-His_6X_ was incubated with nickel beads at 4 °C for 1h. Then, 3.6 μg SerRS protein with/without 1mM ADPR (2x) and/or 10 μM tRNA ser (2x) was added to co-incubated at 4 °C for 2h. After 3 times washing, proteins pulled down by nickel beads are dissolved with sample loading buffer and detected by Western blot.

### Aminoacylation assay

Aminoacylation assays were performed with 200nM SerRS or SIRT2 or SerRS/SIRT2 complex in 50mM HEPES pH7.5, 20mM KCl, 5mM MgCl_2_, 4mM ATP, 2 µM [^3^H]-L-serine, 20 µM L-serine, 2 mM DTT, 4ug/ml pyrophosphatase (Roche), and 20 µM yeast total tRNA. The reaction was initiated by addition of tested protein samples. At varying timer intervals, 5 µL aliquots were removed and applied to a MultiScreen 96-well filter plate (0.45 µm pore size, hydrophobic, low-protein-binding membrane; Merck Millipore) containing 100 µL quench solution (0.3M NaOAc pH3.0, 0.5 mg/mL salmon sperm DNA, 0.1 M EDTA). After all time points were collected, 100 µL of 20% (w/v) trichloroacetic acid (TCA) was added to precipitate nucleic acids. The pellets in the filter plate were then washed 4 times with 200 µL wash solution (100 mM cold serine with 5% TCA) followed by one wash of 200 µL 95% ethanol. 70 µL of freshly prepared 100 mM NaOH was added to each well after the plate was dried out. After 10 min, the solution in the plate was centrifuge into a 96-well flexible PET microplate (PerkinElmer) with 150 µL of Supermix scintillation mixture (PerkinElmer). After mixing, the radioactivity in each well of the plate was measured in a 1450 MicroBeta Micoplate Scintillation and Luminescence Counter (PerkinElmer).

### RNA Extraction and Acidic gel northern blot

HEK293AD cell lines over-expressing SIRT2^WT^ or SIRT2^N168A^ were selected by 1μg/mL puromycin for 7days. SerRS knockdown was achieved by siRNA (Santa Cruz) for 5days. Cells were harvest for both detection of protein expression level and tRNA charging level. Total RNA was extracted by Trizol (Invitrogen) and all deacetylated samples are dissolved in 10mM NaOAc (pH 4.5). For deacetylated samples, total RNA was incubated at 37 °C for 1.5h in 0.2M Tris-HCl (pH 9.5), followed by ethanol precipitation and resuspension in 10mM NaOAc (pH 4.5).

5μg of total RNA from each sample was loaded to an acidic 6.5% polyacrylamide/8 M urea gel and electrophoresed in 0.1 M NaOAc (pH 5.0) at constant 500V for 18h at 4°C^64^. After electrophoresis, RNA was transferred onto an Amersham^TM^ Hybond-N^+^ membrane. The membrane was pre-hybridized for 30min at 65°C for tRNA^Ser(AGA)^ and 60°C for tRNA^Tyr(GTA)^, in a buffer containing 20mM Sodium phosphate (pH 7.2), 0.3M NaCl and 1% SDS. Hybridization was performed overnight at the corresponding temperature using the same buffer with biotin labeled-DNA probes (tRNA^Ser(AGA)^ probe: 5’-TGGCGTAGTCG GCAGGATTCGAACCTGCGCGGGGARACCCCAATGGATTTCTAGTCCATCGCC TTAACCACTCGGCCACGACTAC-3’, tRNA^Tyr(GTA)^ probe : 5’-TCCTTCGAGCCGG ASTCGAACCAGCGACCTAAGGATCTACAGTCCTCCGCTCTACCARCTGAGCTA TCGAAGG-3’)^28^. The membrane was washed with buffer containing 20mM Sodium phosphate (pH 7.2), 0.3M NaCl, 0.1% SDS and 2mM EDTA, and subsequently incubate with HRP-streptavidin (Thermo Scientific) at 37°C for 30min. The aminoacylation fraction was calculated as the ratio of charged tRNA to the total amount of charged and uncharged tRNAs.

### Puromycin incorporation assay

HEK293AD cells transiently overexpressing WT, N168A or H187Y SIRT2 or with SerRS knockdown by siRNA (Cell signaling) were seeded in 6-well plates. Fourteen hours post-transfection, 10µg/ml Puromycin was add to each well and incubated for 30min in the humidified incubator. Cells were then washed once with fresh medium and incubated in fresh medium for an additional 1-1.5 hours. After washing with ice cold PBS, cells were lysed in 500ul/well lysis buffer. Then equal amounts of proteins from each sample were then subjected to SDS-PAGE and analyzed by Western blotting with indicated antibodies.

### Quantification and statistical analysis

The quantification of all the Western blot images was done by using ImageJ. GraphPad Prism 9 was used for data analysis and statistical significance was calculated using unpaired Student’s t-test or two-way ANOVA. Statistically significant differences are indicated in figure legends with the accompanying p values. Error bars in figures indicate the standard deviation except for Fig. 3g, which indicates the standard error of the mean (SEM) for the number of replicates (n≥2). p < 0.05 (*), p < 0.01 (**), p < 0.005 (***) and p < 0.0001 (****) were considered significant.

