## Supplementary Table 1 for "Human SerRS/SIRT2 complex structure reveals cross regulation between translation and NAD^+^ metabolism"

**Supplementary Table 1. Cryo-EM data collection, refinement, and validation**

|  | | **SerRS: SIRT2 (2:2)**  **EMDB: EMD-47105**  **PDB: 9DPI** | | **SerRS: SIRT2 (2:1)**  **EMDB: EMD-47103**  **PDB: 9DPD** | | **SerRS: SIRT2 (1:1)**  **EMDB: XXX**  **PDB: XXX** |
| --- | --- | --- | --- | --- | --- | --- |
| **Data collection** |  | |  | |  |  |
| Microscope | | Talos Arctica | | Talos Arctica | | Talos Arctica |
| Voltage (keV) | | 200 | | 200 | | 200 |
| Detector | | K2 Summit (Counting) | | K2 Summit (Counting) | | K2 Summit (Counting) |
| Magnification (nominal/calibrated) | | 36,000X / 43,478X | | 36,000X / 43,478X | | 36,000X / 43,478X |
| Exposure navigation | | Image shift to 16 holes | | Image shift to 16 holes | | Image shift to 16 holes |
| Data acquisition software | | Leginon | | Leginon | | Leginon |
| Total electron exposure (e^-^/Å^2^) | | 50 | | 50 | | 50 |
| Exposure rate (e^-^/pixel/sec) | | 6.7 | | 6.7 | | 6.7 |
| Frame length (ms)  Number of frames per micrograph | | 100  98 | | 100  98 | | 100  98 |
| Pixel size (Å) | | 1.15 | | 1.15 | | 1.15 |
| Defocus range (µm) | | -0.7 to -1.3 | | -0.7 to -1.3 | | -0.7 to -1.3 |
| Micrographs collected (no.) | | 3520 | | 3520 | | 3520 |
| **Reconstruction** |  | |  | |  |  |
| Image processing package | | CryoSPARC | | CryoSPARC | | CryoSPARC |
| Total extracted particles (no.) | | 5,038,656 | | 5,038,656 | | 5,038,656 |
| Refined particles (no.) | | 247,287 | | 500,718 | | 500,718 |
| Final particles (no.) | | 247,287 | | 500,718 | | 500,718 |
| Symmetry imposed | | C2 | | C1 | | C1 |
| Resolution (Å) |  | |  | |  |  |
| FSC 0.5 (unmasked / masked) | | / | | / | |  |
| FSC 0.143 (unmasked / masked) | | / 3.73 | | / 3.38 | | / 3.42 |
| Resolution range (local) | | 3.3 – 4.5 | | 3.0 – 4.5 | |  |
| 3DFSC Sphericity | |  | |  | |  |
| Sharpening B-factor (Å^2^) | | -269.1 | | -251.8 | |  |
| **Model Composition** |  | |  | |  |  |
| Protein residues | | 1806 | | 1417 | |  |
| Ligands | | 2 | | 1 | |  |
| **Model Refinement** |  | |  | |  |  |
| Refinement package | | Phenix | | Phenix | |  |
| CC (volume / mask) | |  | | 0.63 / 0.63 | |  |
| R.m.s. deviations |  | |  | |  |  |
| Bond lengths | | 0.002 | | 0.006 | |  |
| Bond angles (°) | | 0.504 | | 1.109 | |  |
| **Validation** |  | |  | |  |  |
| Map-to-model FSC 0.5 (unmasked) | | 4.27 | | 4.22 | |  |
| Ramachandran (%) | |  | |  | |  |
| Outliers | | 0.22 | | 0.12 | |  |
| Allowed | | 9.06 | | 7.08 | |  |
| Favored | | 90.72 | | 92.80 | |  |
| MolProbity score | | 2.98 | | 2.63 | |  |
| Poor rotamers (%) | | 3.90 | | 3.07 | |  |
| Clashscore (all atoms) | | 33.16 | | 20.21 | |  |
| C-beta deviations | | 0.44 | | 0.51 | |  |
| CaBLAM Outliers (%) | | 3.13 | | 2.57 | |  |
| EMRinger | | 0.55 | | 0.56 | |  |
