## Supplementary material for "Human SerRS/SIRT2 complex structure reveals cross regulation between translation and NAD^+^ metabolism": Report of 9DPD

### Full wwPDB EM Validation Report (i)

Sep 26, 2024 – 06:58 AM EDT

| | | |
|---|---|---|
| PDB ID | : | 9DPD |
| EMDB ID | : | EMD-47103 |
| Title | : | Cryo-EM structure of SerRS dimer in complex with one SIRT2 |
| Deposited on | : | 2024-09-21 |
| Resolution | : | 3.87 Å(reported) |

**This wwPDB validation report is for manuscript review**

For Manuscript Review

This is a Full wwPDB EM Validation Report.

This report is produced by the wwPDB biocuration pipeline after annotation of the structure.

We welcome your comments at **
A user guide is available at
https://www.wwpdb.org/validation/2017/EMValidationReportHelp
with specific help available everywhere you see the (i) symbol.

The types of validation reports are described at
http://www.wwpdb.org/validation/2017/FAQs#types.

The following versions of software and data (see references (i)) were used in the production of this report:

| | | |
|---|---|---|
| EMDB validation analysis | : | 0.0.1.dev112 |
| Mogul | : | 1.8.5 (274361), CSD as541be (2020) |
| MolProbity | : | 4.02b-467 |
| buster-report | : | 1.1.7 (2018) |
| Percentile statistics | : | 20231227.v01 (using entries in the PDB archive December 27th 2023) |
| MapQ | : | 1.9.13 |
| Ideal geometry (proteins) | : | Engh & Huber (2001) |
| Ideal geometry (DNA, RNA) | : | Parkinson et al. (1996) |
| Validation Pipeline (wwPDB-VP) | : | 2.38.2 |

### 1 Overall quality at a glance (i)

The following experimental techniques were used to determine the structure:
*ELECTRON MICROSCOPY*

The reported resolution of this entry is 3.87 Å.

Percentile scores (ranging between 0-100) for global validation metrics of the entry are shown in the following graphic. The table shows the number of entries on which the scores are based.

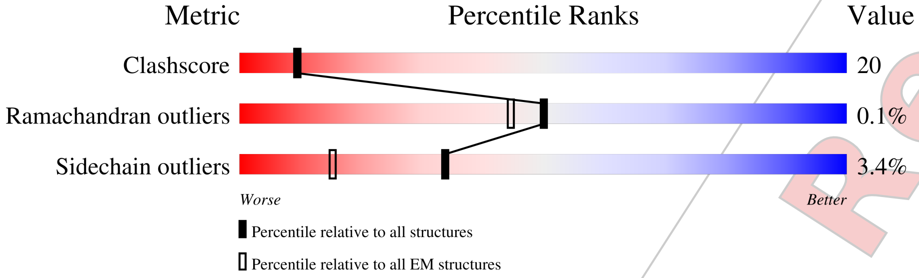


| Metric | Whole archive (#Entries) | EM structures (#Entries) |
|---|---|---|
| Clashscore | 210492 | 15764 |
| Ramachandran outliers | 207382 | 16835 |
| Sidechain outliers | 206894 | 16415 |

The table below summarises the geometric issues observed across the polymeric chains and their fit to the map. The red, orange, yellow and green segments of the bar indicate the fraction of residues that contain outliers for >=3, 2, 1 and 0 types of geometric quality criteria respectively. A grey segment represents the fraction of residues that are not modelled. The numeric value for each fraction is indicated below the corresponding segment, with a dot representing fractions <=5% The upper red bar (where present) indicates the fraction of residues that have poor fit to the EM map (all-atom inclusion < 40%). The numeric value is given above the bar.

| Mol | Chain | Length | Quality of chain |
|---|---|---|---|
| 1 | B | 514 | 6% / 28% 13% • 58% |
| 1 | D | 514 | 8% / 45% 18% • 35% |
| 2 | C | 389 | 11% / 39% 31% • 28% |

For Manuscript Review

### 2 Entry composition (i)

There are 3 unique types of molecules in this entry. The entry contains 6664 atoms, of which 0 are hydrogens and 0 are deuteriums.

In the tables below, the AltConf column contains the number of residues with at least one atom in alternate conformation and the Trace column contains the number of residues modelled with at most 2 atoms.

- Molecule 1 is a protein called Serine–tRNA ligase, cytoplasmic.

| Mol | Chain | Residues | Atoms | | | | | AltConf | Trace |
|---|---|---|---|---|---|---|---|---|---|
| 1 | B | 216 | Total | C | N | O | S | 0 | 0 |
| | | | 1742 | 1128 | 288 | 314 | 12 | | |
| 1 | D | 335 | Total | C | N | O | S | 0 | 0 |
| | | | 2693 | 1724 | 456 | 497 | 16 | | |

- Molecule 2 is a protein called NAD-dependent protein deacetylase sirtuin-2.

| Mol | Chain | Residues | Atoms | | | | | AltConf | Trace |
|---|---|---|---|---|---|---|---|---|---|
| 2 | C | 280 | Total | C | N | O | S | 2 | 0 |
| | | | 2193 | 1401 | 367 | 410 | 15 | | |

- Molecule 3 is [(2R,3S,4R,5R)-5-(6-AMINOPURIN-9-YL)-3,4-DIHYDROXY-OXOLAN-2-YL]METHYL [HYDROXY-[[(2R,3S,4R,5S)-3,4,5-TRIHYDROXYOXOLAN-2-YL]METHOXY]PHOSPHORYL] HYDROGEN PHOSPHATE (three-letter code: AR6) (formula: $C_{15}H_{23}N_5O_{14}P_2$) (labeled as "Ligand of Interest" by depositor).

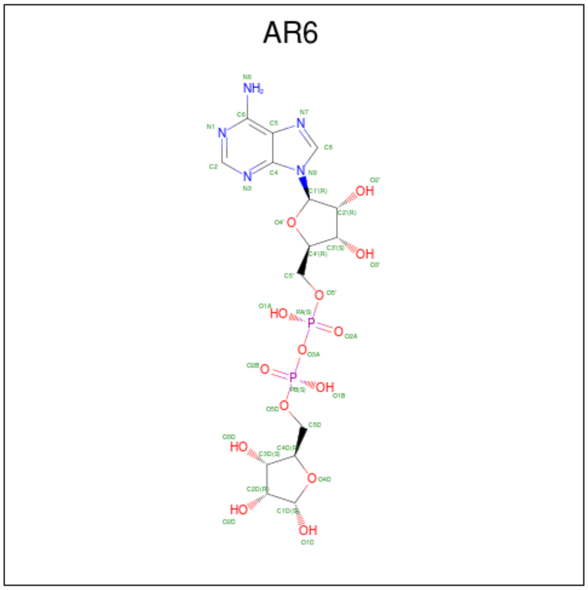

| Mol | Chain | Residues | Atoms | | | | | AltConf |
|---|---|---|---|---|---|---|---|---|
| | | | Total | C | N | O | P | |
| 3 | C | 1 | 36 | 15 | 5 | 14 | 2 | 0 |

For Manuscript Review

### 3 Residue-property plots (i)

These plots are drawn for all protein, RNA, DNA and oligosaccharide chains in the entry. The first graphic for a chain summarises the proportions of the various outlier classes displayed in the second graphic. The second graphic shows the sequence view annotated by issues in geometry and atom inclusion in map density. Residues are color-coded according to the number of geometric quality criteria for which they contain at least one outlier: green = 0, yellow = 1, orange = 2 and red = 3 or more. A red diamond above a residue indicates a poor fit to the EM map for this residue (all-atom inclusion < 40%). Stretches of 2 or more consecutive residues without any outlier are shown as a green connector. Residues present in the sample, but not in the model, are shown in grey.

- Molecule 1: Serine–tRNA ligase, cytoplasmic

Chain B:

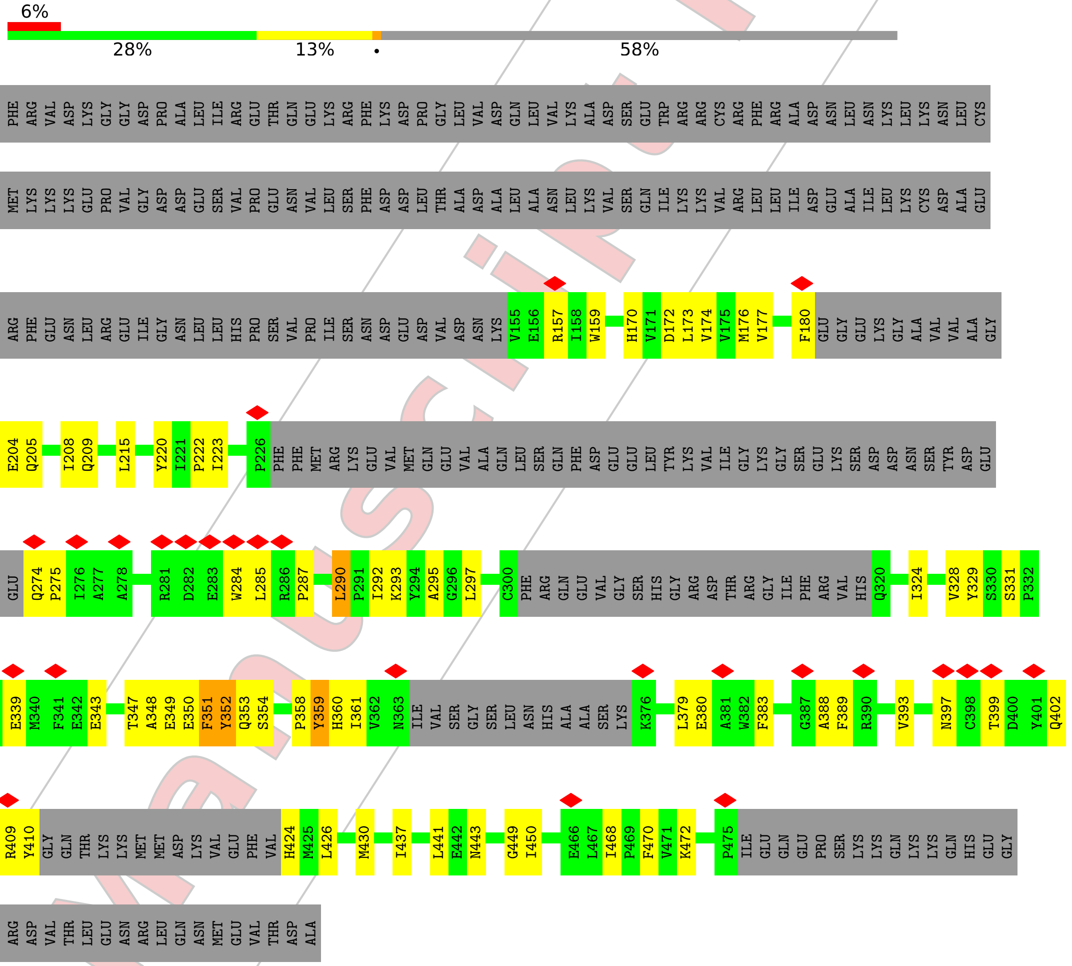


- Molecule 1: Serine–tRNA ligase, cytoplasmic

Chain D:

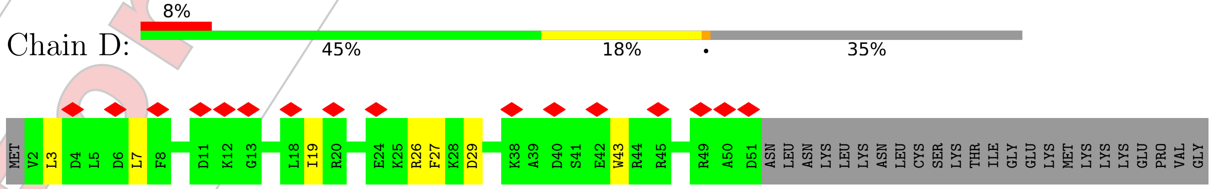

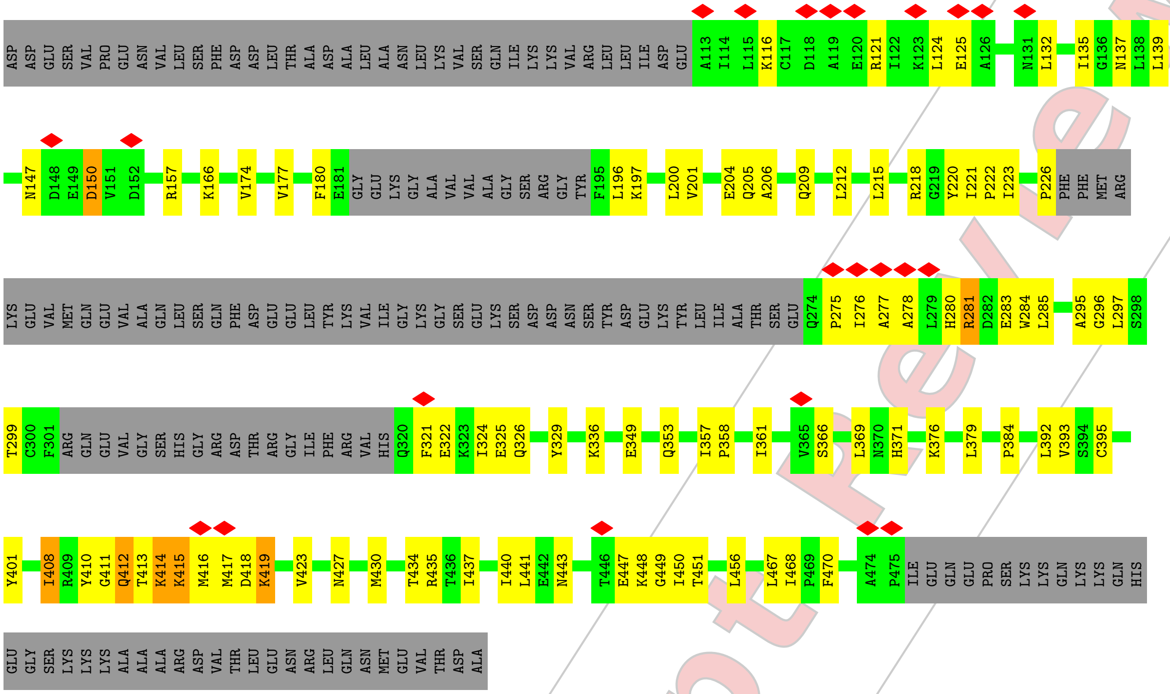


• Molecule 2: NAD-dependent protein deacetylase sirtuin-2

Chain C:

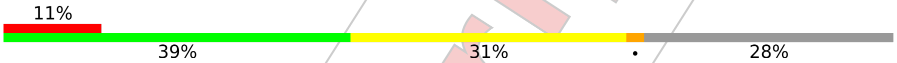


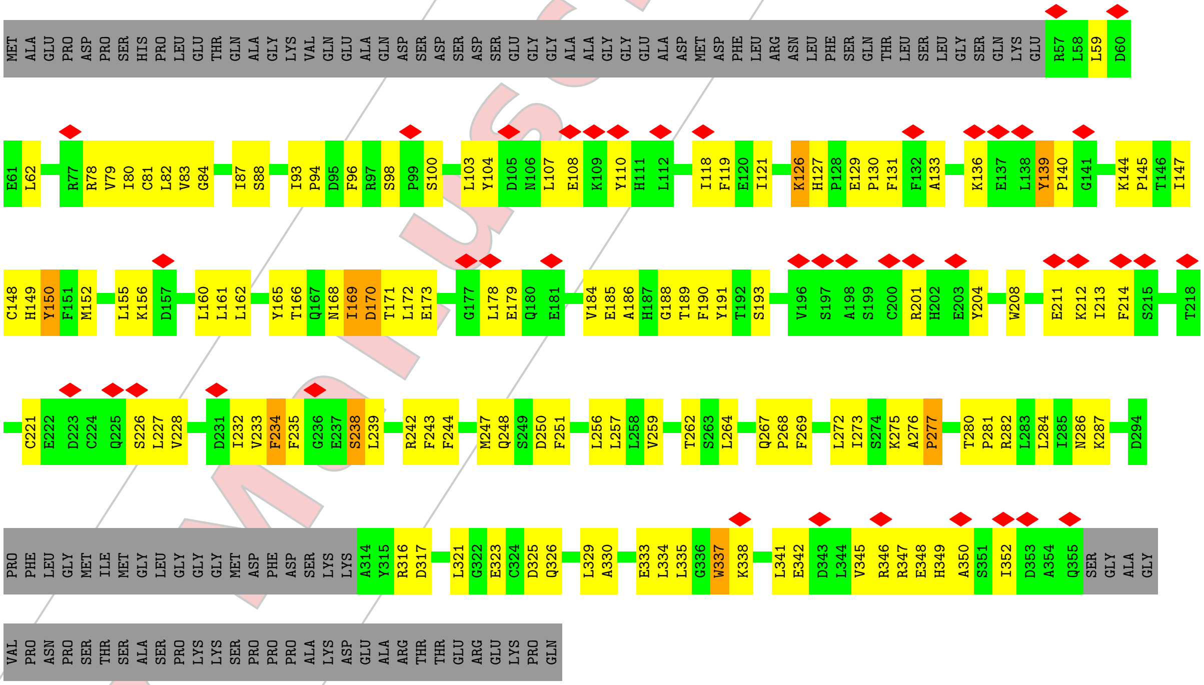


For Manuscript Review

### 4 Experimental information ⓘ

| Property | Value | Source |
|---|---|---|
| EM reconstruction method | SINGLE PARTICLE | Depositor |
| Imposed symmetry | POINT, Not provided | |
| Number of particles used | 590607 | Depositor |
| Resolution determination method | FSC 0.143 CUT-OFF | Depositor |
| CTF correction method | PHASE FLIPPING ONLY | Depositor |
| Microscope | FEI TALOS ARCTICA | Depositor |
| Voltage (kV) | 200 | Depositor |
| Electron dose $(e^-/\text{Å}^2)$ | 50 | Depositor |
| Minimum defocus (nm) | 700 | Depositor |
| Maximum defocus (nm) | 1200 | Depositor |
| Magnification | Not provided | |
| Image detector | GATAN K2 SUMMIT (4k x 4k) | Depositor |
| Maximum map value | 6.615 | Depositor |
| Minimum map value | -5.263 | Depositor |
| Average map value | 0.001 | Depositor |
| Map value standard deviation | 0.126 | Depositor |
| Recommended contour level | 0.651 | Depositor |
| Map size (Å) | 220.79999, 220.79999, 220.79999 | wwPDB |
| Map dimensions | 192, 192, 192 | wwPDB |
| Map angles (°) | 90.0, 90.0, 90.0 | wwPDB |
| Pixel spacing (Å) | 1.15, 1.15, 1.15 | Depositor |

For Manuscript Review

### 5 Model quality (i)

#### 5.1 Standard geometry (i)

Bond lengths and bond angles in the following residue types are not validated in this section: AR6

The Z score for a bond length (or angle) is the number of standard deviations the observed value is removed from the expected value. A bond length (or angle) with $|Z| > 5$ is considered an outlier worth inspection. RMSZ is the root-mean-square of all Z scores of the bond lengths (or angles).

| Mol | Chain | Bond lengths | | Bond angles | |
|---|---|---|---|---|---|
| | | RMSZ | #\|Z\| >5 | RMSZ | #\|Z\| >5 |
| 1 | B | 0.34 | 0/1784 | 0.64 | 3/2416 (0.1%) |
| 1 | D | 0.34 | 0/2750 | 0.58 | 1/3715 (0.0%) |
| 2 | C | 0.34 | 0/2248 | 0.71 | 4/3042 (0.1%) |
| All | All | 0.34 | 0/6782 | 0.64 | 8/9173 (0.1%) |

Chiral center outliers are detected by calculating the chiral volume of a chiral center and verifying if the center is modelled as a planar moiety or with the opposite hand.A planarity outlier is detected by checking planarity of atoms in a peptide group, atoms in a mainchain group or atoms of a sidechain that are expected to be planar.

| Mol | Chain | #Chirality outliers | #Planarity outliers |
|---|---|---|---|
| 1 | B | 0 | 1 |

There are no bond length outliers.

All (8) bond angle outliers are listed below:

| Mol | Chain | Res | Type | Atoms | Z | Observed(°) | Ideal(°) |
|---|---|---|---|---|---|---|---|
| 2 | C | 277 | PRO | N-CA-C | -7.68 | 92.13 | 112.10 |
| 1 | B | 351 | PHE | N-CA-CB | 7.17 | 123.50 | 110.60 |
| 1 | B | 352 | TYR | C-N-CA | -6.47 | 105.52 | 121.70 |
| 2 | C | 234 | PHE | CB-CA-C | 6.36 | 123.11 | 110.40 |
| 1 | B | 351 | PHE | CB-CA-C | -5.66 | 99.08 | 110.40 |
| 2 | C | 170 | ASP | CB-CA-C | 5.42 | 121.25 | 110.40 |
| 2 | C | 234 | PHE | N-CA-CB | -5.07 | 101.48 | 110.60 |
| 1 | D | 408 | ILE | CG1-CB-CG2 | -5.03 | 100.33 | 111.40 |

There are no chirality outliers.

All (1) planarity outliers are listed below:

For Manuscript Review

| Mol | Chain | Res | Type | Group |
|---|---|---|---|---|
| 1 | B | 290 | LEU | Peptide |

#### 5.2 Too-close contacts (i)

In the following table, the Non-H and H(model) columns list the number of non-hydrogen atoms and hydrogen atoms in the chain respectively. The H(added) column lists the number of hydrogen atoms added and optimized by MolProbity. The Clashes column lists the number of clashes within the asymmetric unit, whereas Symm-Clashes lists symmetry-related clashes.

| Mol | Chain | Non-H | H(model) | H(added) | Clashes | Symm-Clashes |
|---|---|---|---|---|---|---|
| 1 | B | 1742 | 0 | 1720 | 60 | 0 |
| 1 | D | 2693 | 0 | 2671 | 78 | 0 |
| 2 | C | 2193 | 0 | 2134 | 135 | 0 |
| 3 | C | 36 | 0 | 21 | 1 | 0 |
| All | All | 6664 | 0 | 6546 | 260 | 0 |

The all-atom clashscore is defined as the number of clashes found per 1000 atoms (including hydrogen atoms). The all-atom clashscore for this structure is 20.

All (260) close contacts within the same asymmetric unit are listed below, sorted by their clash magnitude.

| Atom-1 | Atom-2 | Interatomic distance (Å) | Clash overlap (Å) |
|---|---|---|---|
| 2:C:152:MET:O | 2:C:161:LEU:HD11 | 1.37 | 1.19 |
| 2:C:170:ASP:O | 2:C:171:THR:HG22 | 1.63 | 0.98 |
| 2:C:152:MET:O | 2:C:161:LEU:CD1 | 2.13 | 0.96 |
| 1:D:410:TYR:HB2 | 1:D:423:VAL:HG22 | 1.51 | 0.90 |
| 2:C:188:GLY:HA3 | 2:C:232:ILE:HA | 1.58 | 0.85 |
| 2:C:140:PRO:HG3 | 2:C:170:ASP:O | 1.78 | 0.84 |
| 2:C:277:PRO:O | 2:C:280:THR:HB | 1.78 | 0.82 |
| 2:C:333:GLU:HB2 | 2:C:338:LYS:HD2 | 1.63 | 0.80 |
| 2:C:150:TYR:HE2 | 2:C:345:VAL:HA | 1.46 | 0.79 |
| 2:C:80:ILE:HD13 | 2:C:251:PHE:CE1 | 2.18 | 0.79 |
| 2:C:165:TYR:CZ | 2:C:251:PHE:CE2 | 2.72 | 0.78 |
| 2:C:189:THR:HG22 | 2:C:191:TYR:H | 1.49 | 0.78 |
| 2:C:165:TYR:CE1 | 2:C:251:PHE:CZ | 2.73 | 0.77 |
| 1:D:26:ARG:HE | 1:D:139:LEU:HD11 | 1.50 | 0.76 |
| 1:D:137:ASN:HD22 | 1:D:401:TYR:HB2 | 1.51 | 0.74 |
| 2:C:155:LEU:CB | 2:C:161:LEU:HG | 2.17 | 0.74 |
| 2:C:118:ILE:HG13 | 2:C:119:PHE:HD1 | 1.54 | 0.72 |
| 2:C:323:GLU:HB2 | 2:C:326:GLN:HE22 | 1.54 | 0.72 |
| 2:C:156:LYS:N | 2:C:161:LEU:HD12 | 2.05 | 0.72 |

Continued on next page...

For Manuscript Review

*Continued from previous page...*

| Atom-1 | Atom-2 | Interatomic distance (Å) | Clash overlap (Å) |
|---|---|---|---|
| 2:C:345:VAL:HG23 | 2:C:349:HIS:HD2 | 1.55 | 0.71 |
| 2:C:79:VAL:HG22 | 2:C:256:LEU:HB3 | 1.73 | 0.71 |
| 1:D:174:VAL:HG12 | 1:D:180:PHE:HB2 | 1.73 | 0.70 |
| 2:C:121:ILE:HG12 | 2:C:234:PHE:HB2 | 1.71 | 0.70 |
| 2:C:165:TYR:CE1 | 2:C:251:PHE:HZ | 2.10 | 0.70 |
| 2:C:87:ILE:HG22 | 2:C:148:CYS:SG | 2.31 | 0.69 |
| 2:C:276:ALA:O | 2:C:277:PRO:C | 2.30 | 0.68 |
| 2:C:118:ILE:HG13 | 2:C:119:PHE:CD1 | 2.29 | 0.68 |
| 2:C:165:TYR:CZ | 2:C:251:PHE:HE2 | 2.12 | 0.68 |
| 2:C:233:VAL:HG21 | 2:C:238:SER:O | 1.93 | 0.67 |
| 2:C:129:GLU:HG2 | 2:C:130:PRO:HD3 | 1.77 | 0.67 |
| 2:C:87:ILE:HD11 | 2:C:168:ASN:HD21 | 1.59 | 0.66 |
| 1:B:359:TYR:HE2 | 1:B:361:ILE:HD13 | 1.59 | 0.66 |
| 2:C:165:TYR:OH | 2:C:251:PHE:HE2 | 1.79 | 0.66 |
| 1:B:393:VAL:HG13 | 1:B:430:MET:HA | 1.77 | 0.65 |
| 2:C:165:TYR:OH | 2:C:251:PHE:CE2 | 2.48 | 0.65 |
| 2:C:88:SER:HB2 | 2:C:93:ILE:HD13 | 1.79 | 0.65 |
| 2:C:272:LEU:HD13 | 2:C:275:LYS:HD2 | 1.79 | 0.65 |
| 1:D:451:THR:OG1 | 1:D:467:LEU:CD2 | 2.44 | 0.64 |
| 2:C:329:LEU:HB3 | 2:C:338:LYS:HE3 | 1.79 | 0.64 |
| 1:D:325:GLU:HA | 1:D:430:MET:H | 1.63 | 0.64 |
| 1:B:157:ARG:HH21 | 1:B:159:TRP:HE1 | 1.46 | 0.64 |
| 2:C:221:CYS:HB3 | 2:C:226:SER:H | 1.64 | 0.63 |
| 1:D:222:PRO:HA | 1:D:295:ALA:HB3 | 1.81 | 0.63 |
| 2:C:165:TYR:CZ | 2:C:251:PHE:CZ | 2.87 | 0.63 |
| 1:B:222:PRO:HA | 1:B:295:ALA:HB3 | 1.81 | 0.62 |
| 1:D:212:LEU:HD11 | 1:D:297:LEU:HD22 | 1.81 | 0.62 |
| 1:D:349:GLU:O | 1:D:353:GLN:HG2 | 1.99 | 0.62 |
| 2:C:140:PRO:CG | 2:C:170:ASP:O | 2.48 | 0.62 |
| 2:C:59:LEU:HD21 | 2:C:281:PRO:HB3 | 1.81 | 0.62 |
| 2:C:170:ASP:O | 2:C:171:THR:CG2 | 2.46 | 0.62 |
| 2:C:287:LYS:O | 2:C:321:LEU:HD12 | 1.99 | 0.62 |
| 2:C:345:VAL:HG23 | 2:C:349:HIS:CD2 | 2.34 | 0.62 |
| 1:B:203:LEU:HG | 1:B:437:ILE:HG12 | 1.81 | 0.61 |
| 2:C:156:LYS:HZ2 | 2:C:179:GLU:H | 1.48 | 0.61 |
| 2:C:82:LEU:HG | 2:C:257:LEU:HD11 | 1.82 | 0.61 |
| 2:C:262:THR:HG23 | 2:C:264:LEU:HD22 | 1.83 | 0.61 |
| 1:D:357:ILE:HD13 | 1:D:443:ASN:HD22 | 1.66 | 0.60 |
| 2:C:149:HIS:CE1 | 2:C:172:LEU:HD23 | 2.36 | 0.60 |
| 2:C:208:TRP:NE1 | 2:C:212:LYS:HD3 | 2.16 | 0.60 |
| 1:B:287:PRO:HA | 1:B:290:LEU:HD12 | 1.84 | 0.60 |

*Continued on next page...*

For Manuscript Review

*Continued from previous page...*

| Atom-1 | Atom-2 | Interatomic distance (Å) | Clash overlap (Å) |
|---|---|---|---|
| 2:C:345:VAL:O | 2:C:349:HIS:HB2 | 2.02 | 0.59 |
| 1:D:132:LEU:HD12 | 1:D:135:ILE:HD12 | 1.83 | 0.59 |
| 2:C:149:HIS:HE1 | 2:C:172:LEU:HD23 | 1.67 | 0.59 |
| 1:D:324:ILE:HG22 | 1:D:430:MET:HB2 | 1.85 | 0.59 |
| 1:D:441:LEU:HD22 | 1:D:450:ILE:HG12 | 1.84 | 0.59 |
| 1:B:352:TYR:O | 1:B:353:GLN:C | 2.38 | 0.59 |
| 2:C:259:VAL:HG11 | 2:C:264:LEU:HD21 | 1.85 | 0.59 |
| 2:C:330:ALA:O | 2:C:334:LEU:HD12 | 2.02 | 0.59 |
| 2:C:145:PRO:HD2 | 2:C:352:ILE:HD12 | 1.84 | 0.59 |
| 1:D:177:VAL:HG11 | 1:D:450:ILE:HD11 | 1.83 | 0.59 |
| 1:D:299:THR:HG22 | 1:D:322:GLU:HG3 | 1.85 | 0.58 |
| 1:D:278:ALA:HA | 1:D:281:ARG:HG2 | 1.84 | 0.58 |
| 1:B:359:TYR:CD2 | 1:B:379:LEU:HG | 2.38 | 0.58 |
| 2:C:155:LEU:CB | 2:C:161:LEU:CG | 2.81 | 0.58 |
| 1:D:408:ILE:O | 1:D:423:VAL:HG23 | 2.04 | 0.58 |
| 1:D:414:LYS:HB3 | 2:C:235:PHE:HA | 1.86 | 0.58 |
| 2:C:156:LYS:N | 2:C:161:LEU:CD1 | 2.67 | 0.58 |
| 1:B:290:LEU:HD13 | 1:B:331:SER:HB2 | 1.85 | 0.57 |
| 2:C:96:PHE:HB3 | 2:C:104:TYR:CD2 | 2.39 | 0.57 |
| 2:C:169:ILE:HB | 2:C:190:PHE:CE2 | 2.38 | 0.57 |
| 1:D:285:LEU:HD12 | 1:D:423:VAL:HG21 | 1.87 | 0.57 |
| 1:D:408:ILE:O | 1:D:408:ILE:HG22 | 2.04 | 0.57 |
| 1:B:393:VAL:HG22 | 1:B:430:MET:O | 2.04 | 0.57 |
| 1:D:157:ARG:HG2 | 1:D:361:ILE:HB | 1.86 | 0.57 |
| 1:D:379:LEU:HD22 | 1:D:393:VAL:HG23 | 1.87 | 0.56 |
| 1:B:223:ILE:HG22 | 1:D:197:LYS:HG2 | 1.88 | 0.56 |
| 2:C:87:ILE:HD11 | 2:C:168:ASN:ND2 | 2.19 | 0.56 |
| 2:C:201:ARG:O | 2:C:201:ARG:NH1 | 2.35 | 0.56 |
| 1:D:206:ALA:HB1 | 1:D:456:LEU:HD11 | 1.87 | 0.56 |
| 2:C:184:VAL:HG12 | 2:C:242:ARG:HD3 | 1.87 | 0.56 |
| 1:B:380:GLU:HB3 | 1:B:389:PHE:HB3 | 1.88 | 0.56 |
| 2:C:82:LEU:HB3 | 2:C:269:PHE:CZ | 2.41 | 0.56 |
| 2:C:193:SER:HB3 | 2:C:228:VAL:HG12 | 1.88 | 0.56 |
| 1:B:293:LYS:HA | 1:B:328:VAL:HG12 | 1.88 | 0.55 |
| 2:C:277:PRO:O | 2:C:280:THR:CB | 2.52 | 0.55 |
| 2:C:335:LEU:HG | 2:C:337:TRP:CD1 | 2.41 | 0.55 |
| 2:C:133:ALA:O | 2:C:136:LYS:HG2 | 2.06 | 0.55 |
| 1:B:359:TYR:HE2 | 1:B:361:ILE:CD1 | 2.20 | 0.55 |
| 2:C:80:ILE:HG21 | 2:C:251:PHE:HE1 | 1.70 | 0.55 |
| 1:B:350:GLU:O | 1:B:351:PHE:C | 2.44 | 0.55 |
| 2:C:126:LYS:HE3 | 2:C:127:HIS:ND1 | 2.23 | 0.54 |

*Continued on next page...*

For Manuscript Review

*Continued from previous page...*

| Atom-1 | Atom-2 | Interatomic distance (Å) | Clash overlap (Å) |
|---|---|---|---|
| 2:C:147:ILE:HA | 2:C:150:TYR:HB2 | 1.90 | 0.54 |
| 1:B:173:LEU:O | 1:B:177:VAL:HG22 | 2.07 | 0.54 |
| 1:B:358:PRO:HD2 | 1:B:443:ASN:OD1 | 2.08 | 0.54 |
| 1:B:172:ASP:O | 1:B:176:MET:HG2 | 2.08 | 0.54 |
| 1:B:203:LEU:HD23 | 1:B:437:ILE:HG23 | 1.90 | 0.54 |
| 1:B:331:SER:HB3 | 1:B:336:LYS:HD2 | 1.89 | 0.54 |
| 1:D:379:LEU:HB3 | 1:D:393:VAL:HB | 1.89 | 0.54 |
| 2:C:342:GLU:HA | 2:C:345:VAL:HG12 | 1.90 | 0.54 |
| 2:C:80:ILE:CD1 | 2:C:251:PHE:CE1 | 2.91 | 0.53 |
| 2:C:150:TYR:CE2 | 2:C:345:VAL:HA | 2.35 | 0.53 |
| 2:C:126:LYS:HG2 | 2:C:127:HIS:ND1 | 2.24 | 0.53 |
| 2:C:59:LEU:HD12 | 2:C:59:LEU:H | 1.72 | 0.53 |
| 2:C:208:TRP:CD1 | 2:C:212:LYS:HD3 | 2.43 | 0.53 |
| 1:D:392:LEU:HD22 | 1:D:435:ARG:HB3 | 1.91 | 0.53 |
| 1:D:418:ASP:HA | 2:C:267:GLN:HB2 | 1.91 | 0.53 |
| 2:C:87:ILE:HD12 | 2:C:149:HIS:CD2 | 2.44 | 0.53 |
| 1:D:451:THR:OG1 | 1:D:467:LEU:HD23 | 2.09 | 0.52 |
| 2:C:201:ARG:O | 2:C:201:ARG:CG | 2.57 | 0.52 |
| 1:D:450:ILE:HB | 1:D:468:ILE:HD12 | 1.91 | 0.52 |
| 1:D:321:PHE:HD1 | 1:D:434:THR:HG21 | 1.74 | 0.52 |
| 1:D:223:ILE:HD11 | 1:D:276:ILE:HD11 | 1.91 | 0.52 |
| 1:B:399:THR:HA | 1:B:424:HIS:HB3 | 1.92 | 0.51 |
| 1:D:26:ARG:HH12 | 1:D:371:HIS:HA | 1.74 | 0.51 |
| 2:C:173:GLU:HB3 | 2:C:178:LEU:HD12 | 1.92 | 0.51 |
| 1:B:174:VAL:HG23 | 1:B:180:PHE:HB2 | 1.92 | 0.51 |
| 1:D:358:PRO:HD2 | 1:D:443:ASN:ND2 | 2.25 | 0.51 |
| 1:B:204:GLU:O | 1:B:208:ILE:HG12 | 2.10 | 0.51 |
| 1:B:208:ILE:HG23 | 1:B:324:ILE:HD11 | 1.92 | 0.51 |
| 1:D:166:LYS:HG3 | 1:D:384:PRO:HB2 | 1.92 | 0.51 |
| 2:C:96:PHE:O | 2:C:104:TYR:HB2 | 2.11 | 0.51 |
| 2:C:80:ILE:HG21 | 2:C:251:PHE:CE1 | 2.46 | 0.51 |
| 1:D:196:LEU:HD22 | 1:D:200:LEU:HB3 | 1.93 | 0.50 |
| 1:D:7:LEU:HB3 | 1:D:19:ILE:HD11 | 1.92 | 0.50 |
| 2:C:82:LEU:HB3 | 2:C:269:PHE:HZ | 1.75 | 0.50 |
| 2:C:284:LEU:HD11 | 2:C:286:ASN:HD21 | 1.76 | 0.50 |
| 2:C:346:ARG:HH21 | 2:C:350:ALA:HB2 | 1.77 | 0.50 |
| 2:C:156:LYS:CA | 2:C:161:LEU:HD12 | 2.41 | 0.49 |
| 2:C:325:ASP:O | 2:C:329:LEU:HD22 | 2.12 | 0.49 |
| 2:C:247:MET:HA | 2:C:251:PHE:CD2 | 2.47 | 0.49 |
| 1:D:451:THR:OG1 | 1:D:467:LEU:HD21 | 2.12 | 0.49 |
| 2:C:347:ARG:NH2 | 2:C:348:GLU:HB2 | 2.28 | 0.49 |

*Continued on next page...*

For Manuscript Review

*Continued from previous page...*

| Atom-1 | Atom-2 | Interatomic distance (Å) | Clash overlap (Å) |
|---|---|---|---|
| 1:B:343:GLU:O | 1:B:347:THR:HG23 | 2.12 | 0.49 |
| 2:C:94:PRO:HB2 | 2:C:103:LEU:HD12 | 1.95 | 0.49 |
| 2:C:169:ILE:HG21 | 2:C:232:ILE:HB | 1.94 | 0.49 |
| 1:B:205:GLN:HG3 | 1:D:212:LEU:HD12 | 1.95 | 0.49 |
| 2:C:273:ILE:HB | 2:C:282:ARG:NH1 | 2.27 | 0.48 |
| 2:C:201:ARG:O | 2:C:201:ARG:HG3 | 2.13 | 0.48 |
| 2:C:127:HIS:HB3 | 2:C:129:GLU:OE2 | 2.13 | 0.48 |
| 1:B:285:LEU:H | 1:B:410:TYR:HA | 1.77 | 0.48 |
| 1:B:397:ASN:HA | 1:B:426:LEU:HD13 | 1.94 | 0.48 |
| 1:D:326:GLN:HG3 | 1:D:430:MET:CG | 2.43 | 0.48 |
| 1:D:415:LYS:O | 2:C:267:GLN:O | 2.32 | 0.48 |
| 2:C:341:LEU:O | 2:C:345:VAL:HG12 | 2.13 | 0.47 |
| 1:D:221:ILE:O | 1:D:223:ILE:HG23 | 2.14 | 0.47 |
| 2:C:136:LYS:N | 2:C:136:LYS:HD2 | 2.28 | 0.47 |
| 1:D:376:LYS:HG3 | 1:D:395:CYS:O | 2.14 | 0.47 |
| 1:B:159:TRP:HB3 | 1:B:353:GLN:HE22 | 1.80 | 0.47 |
| 1:D:27:PHE:CD1 | 1:D:147:ASN:HB2 | 2.50 | 0.47 |
| 1:B:336:LYS:HB2 | 1:B:336:LYS:HE3 | 1.68 | 0.47 |
| 2:C:108:GLU:H | 2:C:108:GLU:CD | 2.18 | 0.47 |
| 1:B:441:LEU:HA | 1:B:450:ILE:HD12 | 1.97 | 0.46 |
| 1:B:348:ALA:O | 1:B:349:GLU:C | 2.53 | 0.46 |
| 2:C:247:MET:HA | 2:C:251:PHE:HD2 | 1.81 | 0.46 |
| 2:C:96:PHE:HB3 | 2:C:104:TYR:CE2 | 2.51 | 0.46 |
| 2:C:96:PHE:HB3 | 2:C:104:TYR:HD2 | 1.79 | 0.46 |
| 2:C:204:TYR:HD2 | 2:C:208:TRP:CZ3 | 2.34 | 0.46 |
| 1:D:414:LYS:HA | 2:C:268:PRO:HD3 | 1.97 | 0.45 |
| 1:D:336:LYS:HB2 | 1:D:336:LYS:HE3 | 1.78 | 0.45 |
| 1:B:222:PRO:HB2 | 1:D:201:VAL:HG11 | 1.97 | 0.45 |
| 1:B:336:LYS:HA | 1:B:339:GLU:OE2 | 2.17 | 0.45 |
| 1:B:379:LEU:HD22 | 1:B:393:VAL:HG23 | 1.96 | 0.45 |
| 2:C:244:PHE:O | 2:C:248:GLN:HG2 | 2.16 | 0.45 |
| 1:B:284:TRP:HA | 1:B:409:ARG:O | 2.16 | 0.45 |
| 1:D:419:LYS:HE2 | 1:D:419:LYS:HB2 | 1.73 | 0.45 |
| 1:D:43:TRP:HB2 | 1:D:124:LEU:HD13 | 1.99 | 0.45 |
| 2:C:208:TRP:O | 2:C:212:LYS:HG2 | 2.17 | 0.45 |
| 1:D:137:ASN:ND2 | 1:D:401:TYR:HB2 | 2.26 | 0.45 |
| 2:C:155:LEU:CB | 2:C:161:LEU:HD21 | 2.47 | 0.45 |
| 1:B:349:GLU:O | 1:B:350:GLU:C | 2.55 | 0.45 |
| 1:D:223:ILE:HD11 | 1:D:276:ILE:CD1 | 2.47 | 0.45 |
| 2:C:59:LEU:HD13 | 2:C:316:ARG:HA | 1.98 | 0.45 |
| 1:B:215:LEU:HD13 | 1:B:295:ALA:HB1 | 1.98 | 0.44 |

*Continued on next page...*

For Manuscript Review

Continued from previous page...

| Atom-1 | Atom-2 | Interatomic distance (Å) | Clash overlap (Å) |
|---|---|---|---|
| 1:D:205:GLN:HA | 1:D:205:GLN:OE1 | 2.17 | 0.44 |
| 1:B:406:LEU:HD23 | 1:B:406:LEU:HA | 1.82 | 0.44 |
| 2:C:189:THR:HG22 | 2:C:191:TYR:N | 2.25 | 0.44 |
| 2:C:84:GLY:O | 2:C:87:ILE:HG12 | 2.17 | 0.44 |
| 1:D:447:GLU:HG3 | 1:D:448:LYS:HD2 | 2.00 | 0.44 |
| 2:C:62:LEU:HD12 | 2:C:317:ASP:O | 2.17 | 0.44 |
| 1:B:450:ILE:HB | 1:B:468:ILE:HD12 | 2.00 | 0.44 |
| 2:C:238:SER:O | 2:C:239:LEU:HD12 | 2.17 | 0.44 |
| 1:D:204:GLU:HB2 | 1:D:437:ILE:HD11 | 1.99 | 0.44 |
| 2:C:211:GLU:OE1 | 2:C:212:LYS:HD2 | 2.16 | 0.44 |
| 1:B:351:PHE:O | 1:B:354:SER:N | 2.48 | 0.44 |
| 1:D:358:PRO:HD2 | 1:D:443:ASN:HD21 | 1.84 | 0.43 |
| 1:D:366:SER:HA | 1:D:369:LEU:HD12 | 2.00 | 0.43 |
| 1:B:215:LEU:O | 1:B:220:TYR:HB2 | 2.18 | 0.43 |
| 2:C:155:LEU:CB | 2:C:161:LEU:CD2 | 2.96 | 0.43 |
| 2:C:78:ARG:HB2 | 2:C:162:LEU:HB2 | 2.01 | 0.43 |
| 2:C:82:LEU:O | 2:C:259:VAL:HA | 2.18 | 0.43 |
| 2:C:273:ILE:HB | 2:C:282:ARG:HH11 | 1.83 | 0.43 |
| 1:D:296:GLY:HA3 | 1:D:325:GLU:HG2 | 2.00 | 0.43 |
| 1:D:226:PRO:HG3 | 1:D:275:PRO:HB2 | 2.01 | 0.43 |
| 2:C:144:LYS:HD2 | 2:C:349:HIS:CE1 | 2.53 | 0.43 |
| 2:C:156:LYS:NZ | 2:C:179:GLU:H | 2.14 | 0.43 |
| 1:B:209:GLN:OE1 | 1:D:205:GLN:HB3 | 2.18 | 0.43 |
| 1:B:328:VAL:HG23 | 1:B:426:LEU:HB2 | 1.99 | 0.43 |
| 1:D:408:ILE:HD12 | 1:D:408:ILE:HG23 | 1.80 | 0.43 |
| 2:C:81:CYS:SG | 2:C:161:LEU:HD22 | 2.59 | 0.43 |
| 2:C:342:GLU:HA | 2:C:345:VAL:CG1 | 2.49 | 0.43 |
| 1:B:205:GLN:HB3 | 1:D:209:GLN:OE1 | 2.18 | 0.43 |
| 1:D:357:ILE:HD11 | 1:D:440:ILE:HG12 | 2.00 | 0.43 |
| 1:D:361:ILE:HD13 | 1:D:361:ILE:HA | 1.86 | 0.43 |
| 2:C:213:ILE:HG22 | 2:C:214:PHE:CD1 | 2.54 | 0.43 |
| 2:C:323:GLU:HA | 3:C:401:AR6:N1 | 2.34 | 0.42 |
| 1:B:383:PHE:HB2 | 1:B:388:ALA:O | 2.18 | 0.42 |
| 1:B:220:TYR:CD1 | 1:B:293:LYS:HB3 | 2.55 | 0.42 |
| 2:C:147:ILE:O | 2:C:150:TYR:HB2 | 2.18 | 0.42 |
| 1:B:297:LEU:HD21 | 1:D:205:GLN:NE2 | 2.34 | 0.42 |
| 1:B:404:ARG:HA | 1:B:404:ARG:CZ | 2.49 | 0.42 |
| 2:C:149:HIS:CD2 | 2:C:149:HIS:N | 2.87 | 0.42 |
| 2:C:166:THR:HG21 | 2:C:173:GLU:HG2 | 2.02 | 0.42 |
| 1:B:402:GLN:O | 1:B:406:LEU:HB2 | 2.19 | 0.42 |
| 2:C:326:GLN:H | 2:C:326:GLN:CD | 2.22 | 0.42 |

Continued on next page...

For Manuscript Review

*Continued from previous page...*

| Atom-1 | Atom-2 | Interatomic distance (Å) | Clash overlap (Å) |
|---|---|---|---|
| 1:B:292:ILE:CG2 | 1:B:329:TYR:HB2 | 2.49 | 0.42 |
| 1:D:218:ARG:NH1 | 1:D:218:ARG:HB2 | 2.34 | 0.42 |
| 1:B:195:PHE:CZ | 1:D:226:PRO:HD3 | 2.55 | 0.42 |
| 1:D:3:LEU:HD21 | 1:D:137:ASN:HA | 2.01 | 0.42 |
| 2:C:93:ILE:HD12 | 2:C:93:ILE:H | 1.85 | 0.42 |
| 2:C:145:PRO:HA | 2:C:172:LEU:HD21 | 2.02 | 0.42 |
| 2:C:227:LEU:HD12 | 2:C:227:LEU:H | 1.85 | 0.42 |
| 1:D:215:LEU:O | 1:D:220:TYR:HB2 | 2.20 | 0.41 |
| 1:D:416:MET:HB2 | 2:C:235:PHE:CD1 | 2.55 | 0.41 |
| 2:C:59:LEU:HD11 | 2:C:281:PRO:HG3 | 2.02 | 0.41 |
| 1:D:284:TRP:HB3 | 1:D:412:GLN:HG2 | 2.01 | 0.41 |
| 1:B:170:HIS:O | 1:B:174:VAL:HG13 | 2.20 | 0.41 |
| 1:D:150:ASP:OD1 | 1:D:150:ASP:N | 2.53 | 0.41 |
| 2:C:186:ALA:HB2 | 2:C:243:PHE:CD1 | 2.55 | 0.41 |
| 2:C:189:THR:HG22 | 2:C:191:TYR:HB2 | 2.03 | 0.41 |
| 1:B:449:GLY:HA2 | 1:B:470:PHE:CZ | 2.55 | 0.41 |
| 1:D:326:GLN:O | 1:D:427:ASN:HA | 2.20 | 0.41 |
| 1:D:449:GLY:HA3 | 1:D:468:ILE:O | 2.20 | 0.41 |
| 2:C:87:ILE:HD12 | 2:C:149:HIS:NE2 | 2.35 | 0.41 |
| 1:B:351:PHE:O | 1:B:352:TYR:C | 2.58 | 0.41 |
| 1:D:280:HIS:O | 1:D:283:GLU:HB3 | 2.20 | 0.41 |
| 2:C:107:LEU:HA | 2:C:110:TYR:HD2 | 1.84 | 0.41 |
| 1:B:223:ILE:CG2 | 1:D:197:LYS:HG2 | 2.50 | 0.41 |
| 1:D:121:ARG:O | 1:D:125:GLU:HG2 | 2.21 | 0.41 |
| 2:C:83:VAL:CG2 | 2:C:87:ILE:HD13 | 2.51 | 0.41 |
| 2:C:139:TYR:HD1 | 2:C:140:PRO:HD2 | 1.85 | 0.41 |
| 1:B:274:GLN:HB3 | 1:B:275:PRO:HD3 | 2.03 | 0.41 |
| 1:D:277:ALA:HA | 1:D:329:TYR:OH | 2.21 | 0.41 |
| 1:D:449:GLY:HA2 | 1:D:470:PHE:CZ | 2.56 | 0.41 |
| 2:C:79:VAL:N | 2:C:160:LEU:O | 2.54 | 0.41 |
| 1:B:173:LEU:HD12 | 1:B:173:LEU:HA | 1.88 | 0.40 |
| 1:B:360:HIS:HE1 | 1:B:389:PHE:CG | 2.39 | 0.40 |
| 1:B:404:ARG:HA | 1:B:404:ARG:NH1 | 2.36 | 0.40 |
| 2:C:349:HIS:HA | 2:C:352:ILE:HG12 | 2.03 | 0.40 |
| 2:C:275:LYS:O | 2:C:276:ALA:C | 2.60 | 0.40 |
| 1:D:205:GLN:O | 1:D:209:GLN:HG2 | 2.21 | 0.40 |

There are no symmetry-related clashes.

For Manuscript Review

#### 5.3 Torsion angles (i)

##### 5.3.1 Protein backbone (i)

In the following table, the Percentiles column shows the percent Ramachandran outliers of the chain as a percentile score with respect to all PDB entries followed by that with respect to all EM entries.

The Analysed column shows the number of residues for which the backbone conformation was analysed, and the total number of residues.

| Mol | Chain | Analysed | Favoured | Allowed | Outliers | Percentiles | |
|---|---|---|---|---|---|---|---|
| 1 | B | 204/514 (40%) | 193 (95%) | 11 (5%) | 0 | 100 | 100 |
| 1 | D | 325/514 (63%) | 305 (94%) | 19 (6%) | 1 (0%) | 37 | 70 |
| 2 | C | 278/389 (72%) | 251 (90%) | 27 (10%) | 0 | 100 | 100 |
| All | All | 807/1417 (57%) | 749 (93%) | 57 (7%) | 1 (0%) | 50 | 80 |

All (1) Ramachandran outliers are listed below:

| Mol | Chain | Res | Type |
|---|---|---|---|
| 1 | D | 411 | GLY |

##### 5.3.2 Protein sidechains (i)

In the following table, the Percentiles column shows the percent sidechain outliers of the chain as a percentile score with respect to all PDB entries followed by that with respect to all EM entries.

The Analysed column shows the number of residues for which the sidechain conformation was analysed, and the total number of residues.

| Mol | Chain | Analysed | Rotameric | Outliers | Percentiles | |
|---|---|---|---|---|---|---|
| 1 | B | 189/453 (42%) | 186 (98%) | 3 (2%) | 58 | 73 |
| 1 | D | 293/453 (65%) | 283 (97%) | 10 (3%) | 32 | 55 |
| 2 | C | 235/332 (71%) | 224 (95%) | 11 (5%) | 22 | 47 |
| All | All | 717/1238 (58%) | 693 (97%) | 24 (3%) | 34 | 56 |

All (24) residues with a non-rotameric sidechain are listed below:

| Mol | Chain | Res | Type |
|---|---|---|---|
| 1 | B | 335 | ASN |
| 1 | B | 359 | TYR |

*Continued on next page...*

For Manuscript Review

*Continued from previous page...*

| Mol | Chain | Res | Type |
|---|---|---|---|
| 1 | B | 472 | LYS |
| 1 | D | 29 | ASP |
| 1 | D | 116 | LYS |
| 1 | D | 150 | ASP |
| 1 | D | 281 | ARG |
| 1 | D | 412 | GLN |
| 1 | D | 413 | THR |
| 1 | D | 414 | LYS |
| 1 | D | 415 | LYS |
| 1 | D | 417 | MET |
| 1 | D | 419 | LYS |
| 2 | C | 98 | SER |
| 2 | C | 100 | SER |
| 2 | C | 126 | LYS |
| 2 | C | 131 | PHE |
| 2 | C | 139 | TYR |
| 2 | C | 150 | TYR |
| 2 | C | 169 | ILE |
| 2 | C | 185 | GLU |
| 2 | C | 238 | SER |
| 2 | C | 250 | ASP |
| 2 | C | 337 | TRP |

Sometimes sidechains can be flipped to improve hydrogen bonding and reduce clashes. All (6) such sidechains are listed below:

| Mol | Chain | Res | Type |
|---|---|---|---|
| 1 | B | 360 | HIS |
| 1 | D | 35 | GLN |
| 1 | D | 137 | ASN |
| 1 | D | 443 | ASN |
| 2 | C | 149 | HIS |
| 2 | C | 349 | HIS |

##### 5.3.3 RNA ⓘ

There are no RNA molecules in this entry.

#### 5.4 Non-standard residues in protein, DNA, RNA chains ⓘ

There are no non-standard protein/DNA/RNA residues in this entry.

For Manuscript Review

#### 5.5 Carbohydrates (i)

There are no oligosaccharides in this entry.

#### 5.6 Ligand geometry (i)

1 ligand is modelled in this entry.

In the following table, the Counts columns list the number of bonds (or angles) for which Mogul statistics could be retrieved, the number of bonds (or angles) that are observed in the model and the number of bonds (or angles) that are defined in the Chemical Component Dictionary. The Link column lists molecule types, if any, to which the group is linked. The Z score for a bond length (or angle) is the number of standard deviations the observed value is removed from the expected value. A bond length (or angle) with $|Z| > 2$ is considered an outlier worth inspection. RMSZ is the root-mean-square of all Z scores of the bond lengths (or angles).

| Mol | Type | Chain | Res | Link | Bond lengths | | | Bond angles | | |
|---|---|---|---|---|---|---|---|---|---|---|
| | | | | | Counts | RMSZ | $\#\|Z\| > 2$ | Counts | RMSZ | $\#\|Z\| > 2$ |
| 3 | AR6 | C | 401 | - | 34,39,39 | 0.95 | 1 (2%) | 40,60,60 | 1.04 | 2 (5%) |

In the following table, the Chirals column lists the number of chiral outliers, the number of chiral centers analysed, the number of these observed in the model and the number defined in the Chemical Component Dictionary. Similar counts are reported in the Torsion and Rings columns. '-' means no outliers of that kind were identified.

| Mol | Type | Chain | Res | Link | Chirals | Torsions | Rings |
|---|---|---|---|---|---|---|---|
| 3 | AR6 | C | 401 | - | - | 4/18/54/54 | 0/4/4/4 |

All (1) bond length outliers are listed below:

| Mol | Chain | Res | Type | Atoms | Z | Observed(Å) | Ideal(Å) |
|---|---|---|---|---|---|---|---|
| 3 | C | 401 | AR6 | C8-N7 | -2.29 | 1.30 | 1.34 |

All (2) bond angle outliers are listed below:

| Mol | Chain | Res | Type | Atoms | Z | Observed($^o$) | Ideal($^o$) |
|---|---|---|---|---|---|---|---|
| 3 | C | 401 | AR6 | PB-O3A-PA | -3.04 | 122.41 | 132.83 |
| 3 | C | 401 | AR6 | O4D-C1D-C2D | 2.55 | 107.61 | 104.46 |

There are no chirality outliers.

All (4) torsion outliers are listed below:

For Manuscript Review

| Mol | Chain | Res | Type | Atoms |
|---|---|---|---|---|
| 3 | C | 401 | AR6 | C5D-O5D-PB-O2B |
| 3 | C | 401 | AR6 | O4D-C4D-C5D-O5D |
| 3 | C | 401 | AR6 | C3D-C4D-C5D-O5D |
| 3 | C | 401 | AR6 | C3'-C4'-C5'-O5' |

There are no ring outliers.

1 monomer is involved in 1 short contact:

| Mol | Chain | Res | Type | Clashes | Symm-Clashes |
|---|---|---|---|---|---|
| 3 | C | 401 | AR6 | 1 | 0 |

The following is a two-dimensional graphical depiction of Mogul quality analysis of bond lengths, bond angles, torsion angles, and ring geometry for all instances of the Ligand of Interest. In addition, ligands with molecular weight > 250 and outliers as shown on the validation Tables will also be included. For torsion angles, if less then 5% of the Mogul distribution of torsion angles is within 10 degrees of the torsion angle in question, then that torsion angle is considered an outlier. Any bond that is central to one or more torsion angles identified as an outlier by Mogul will be highlighted in the graph. For rings, the root-mean-square deviation (RMSD) between the ring in question and similar rings identified by Mogul is calculated over all ring torsion angles. If the average RMSD is greater than 60 degrees and the minimal RMSD between the ring in question and any Mogul-identified rings is also greater than 60 degrees, then that ring is considered an outlier. The outliers are highlighted in purple. The color gray indicates Mogul did not find sufficient equivalents in the CSD to analyse the geometry.

For Manuscript Review

**Ligand AR6 C 401**

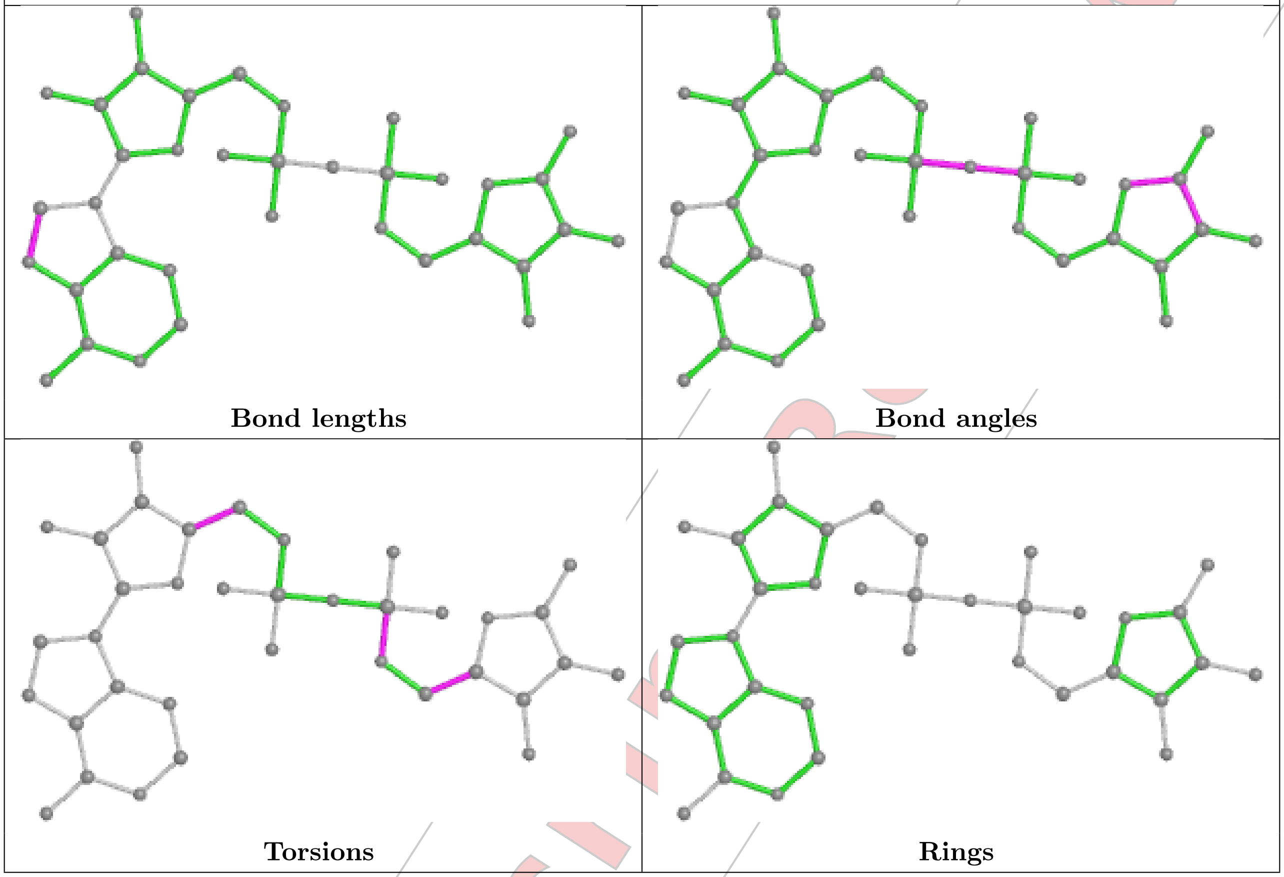


#### 5.7 Other polymers (i)

There are no such residues in this entry.

#### 5.8 Polymer linkage issues (i)

There are no chain breaks in this entry.

For Manuscript Review

### 6 Map visualisation (i)

This section contains visualisations of the EMDB entry EMD-47103. These allow visual inspection of the internal detail of the map and identification of artifacts.

Images derived from a raw map, generated by summing the deposited half-maps, are presented below the corresponding image components of the primary map to allow further visual inspection and comparison with those of the primary map.

#### 6.1 Orthogonal projections (i)

##### 6.1.1 Primary map

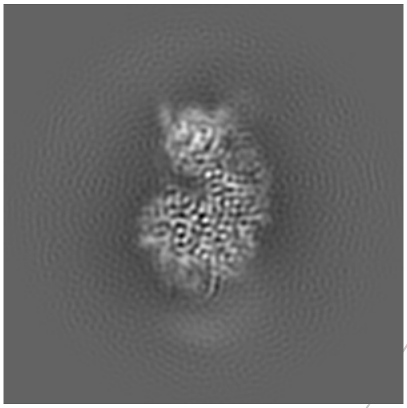
X

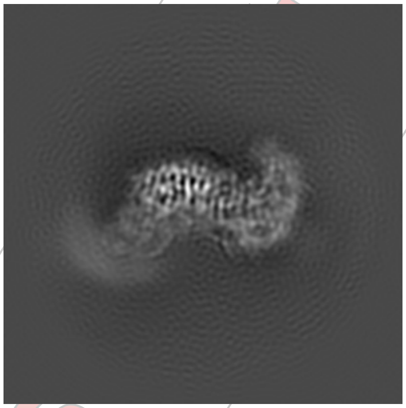
Y

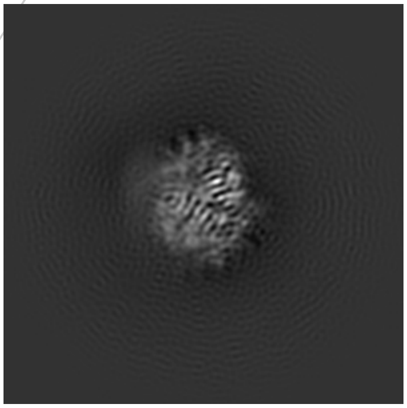
Z

##### 6.1.2 Raw map

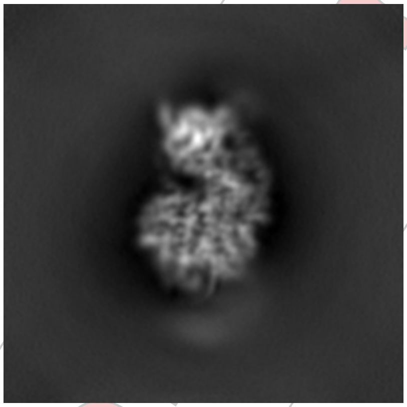
X

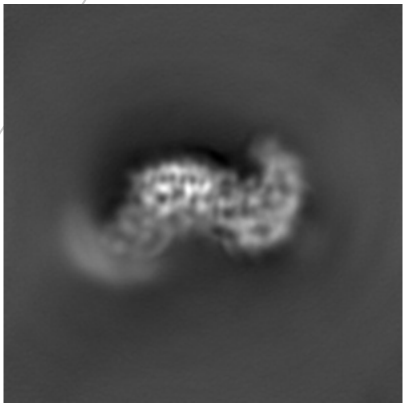
Y

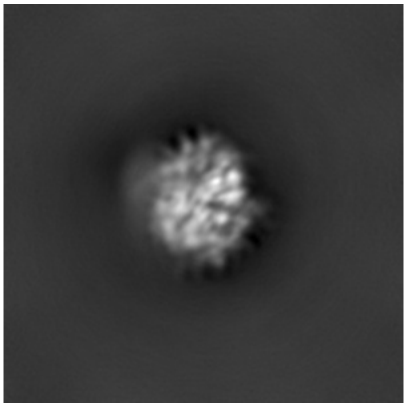
Z

The images above show the map projected in three orthogonal directions.

*For Manuscript Review*

#### 6.2 Central slices (i)

##### 6.2.1 Primary map

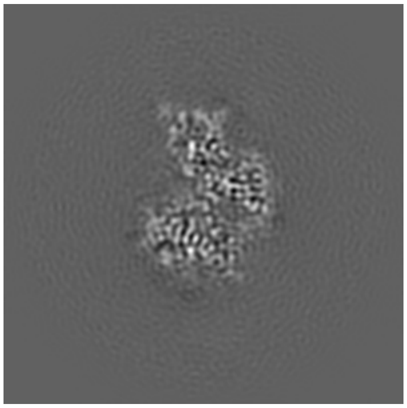

X Index: 96

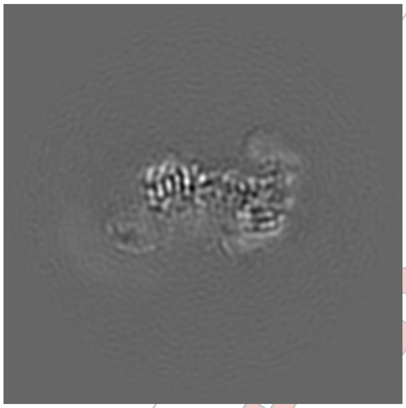

Y Index: 96

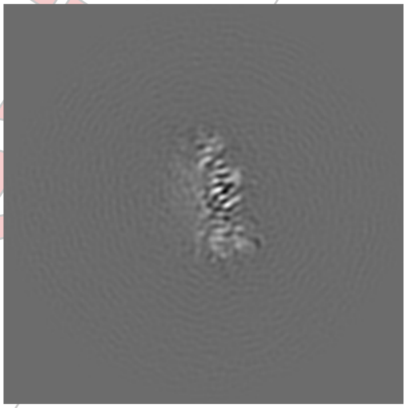

Z Index: 96

##### 6.2.2 Raw map

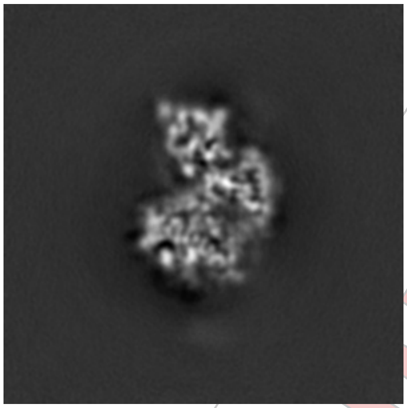

X Index: 96

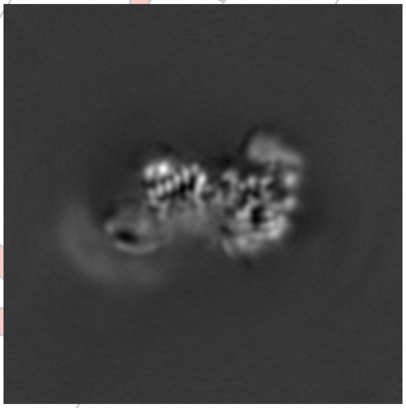

Y Index: 96

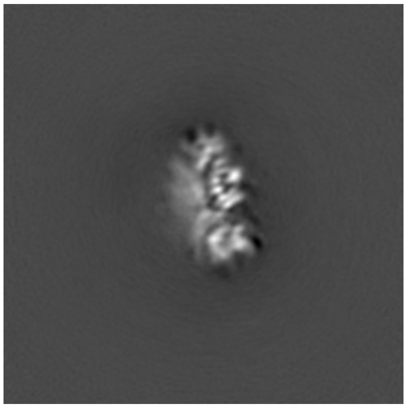

Z Index: 96

The images above show central slices of the map in three orthogonal directions.

For Manuscript Review

#### 6.3 Largest variance slices ⓘ

##### 6.3.1 Primary map

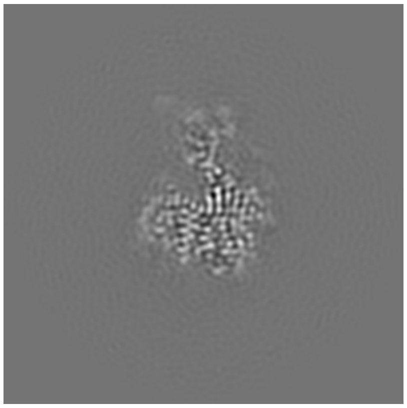
X Index: 104

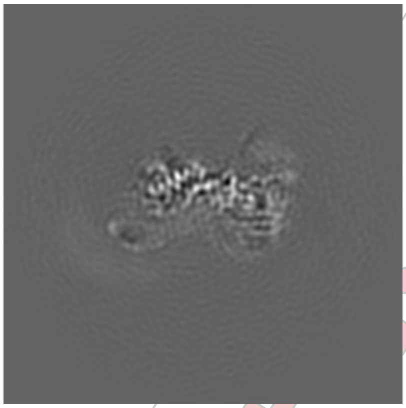
Y Index: 98

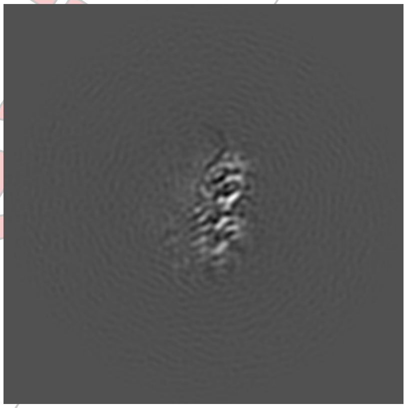
Z Index: 83

##### 6.3.2 Raw map

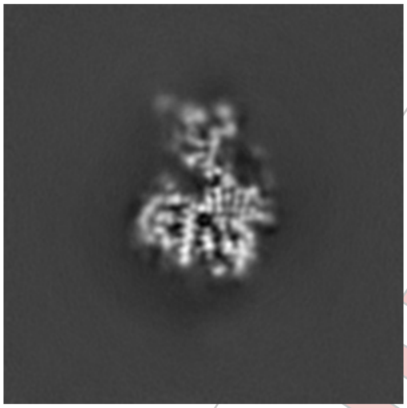
X Index: 104

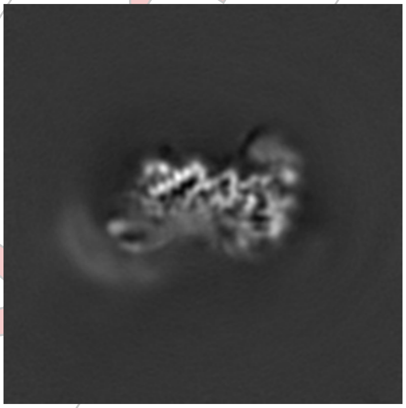
Y Index: 98

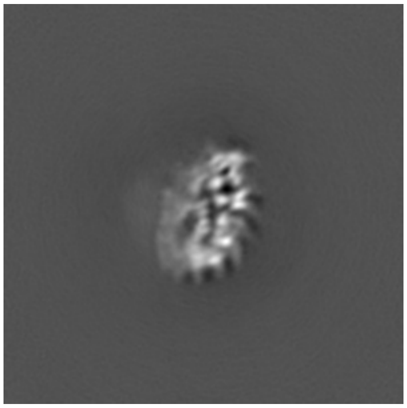
Z Index: 78

The images above show the largest variance slices of the map in three orthogonal directions.

For Manuscript Review

#### 6.4 Orthogonal standard-deviation projections (False-color) (i)

##### 6.4.1 Primary map

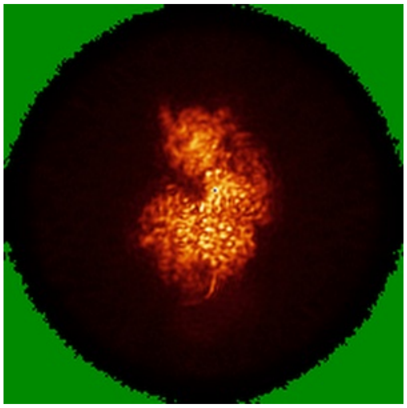
X

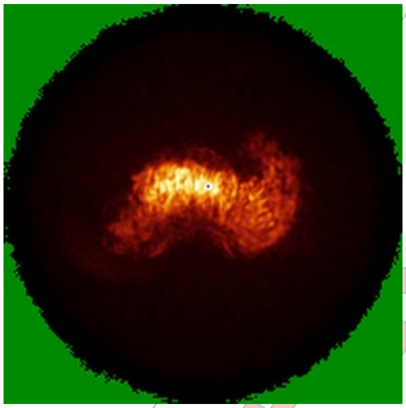
Y

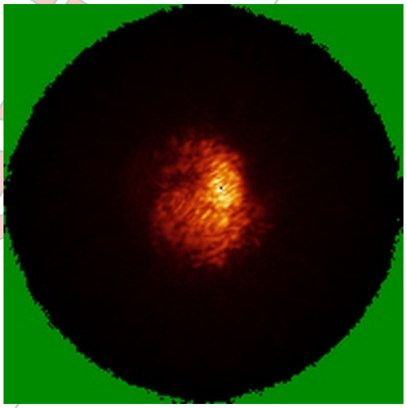
Z

##### 6.4.2 Raw map

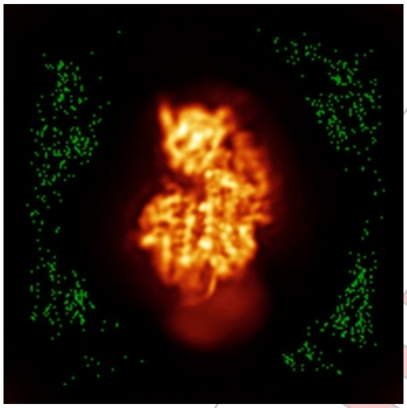
X

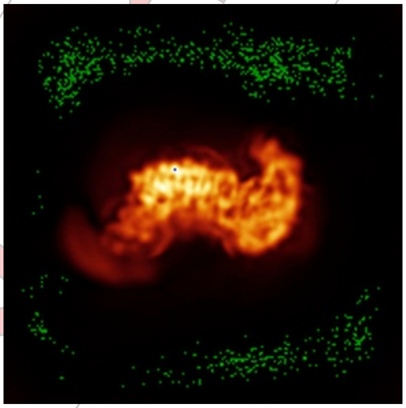
Y

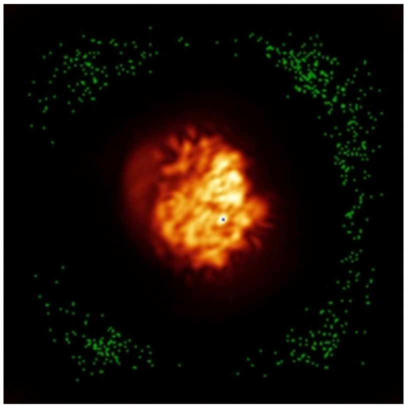
Z

The images above show the map standard deviation projections with false color in three orthogonal directions. Minimum values are shown in green, max in blue, and dark to light orange shades represent small to large values respectively.

For Manuscript Review

#### 6.5 Orthogonal surface views (i)

##### 6.5.1 Primary map

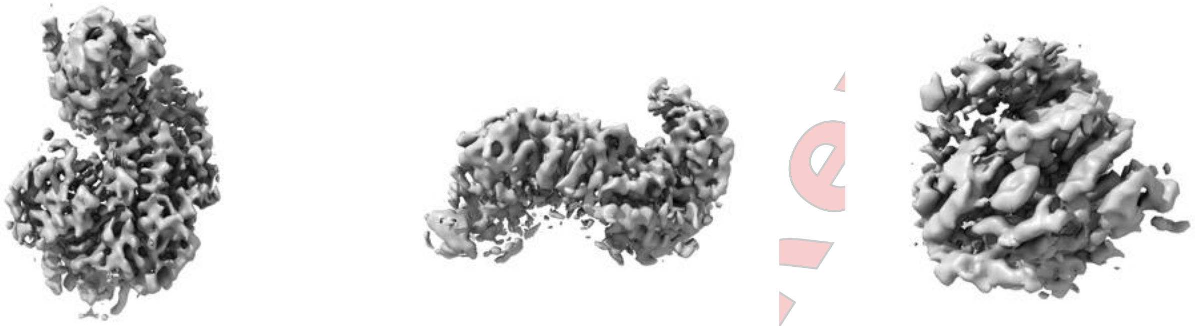

X Y Z

The images above show the 3D surface view of the map at the recommended contour level 0.651. These images, in conjunction with the slice images, may facilitate assessment of whether an appropriate contour level has been provided.

##### 6.5.2 Raw map

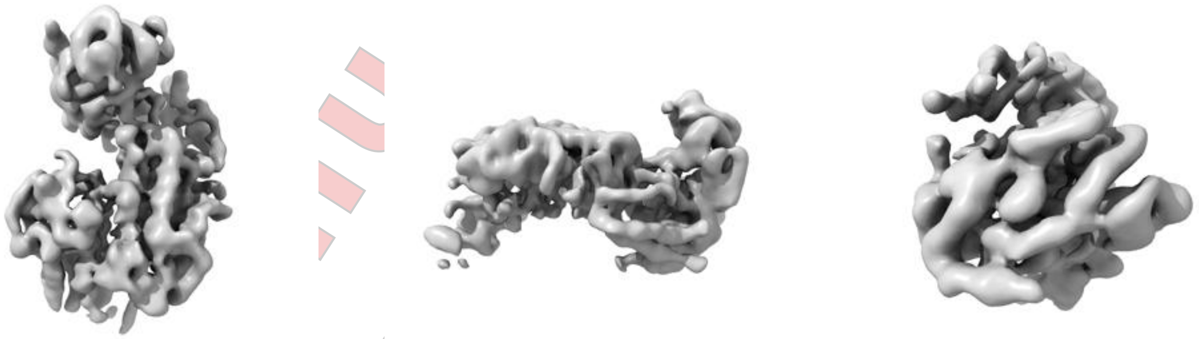

X Y Z

These images show the 3D surface of the raw map. The raw map's contour level was selected so that its surface encloses the same volume as the primary map does at its recommended contour level.

#### 6.6 Mask visualisation (i)

This section was not generated. No masks/segmentation were deposited.

For Manuscript Review

### 7 Map analysis (i)

This section contains the results of statistical analysis of the map.

#### 7.1 Map-value distribution (i)

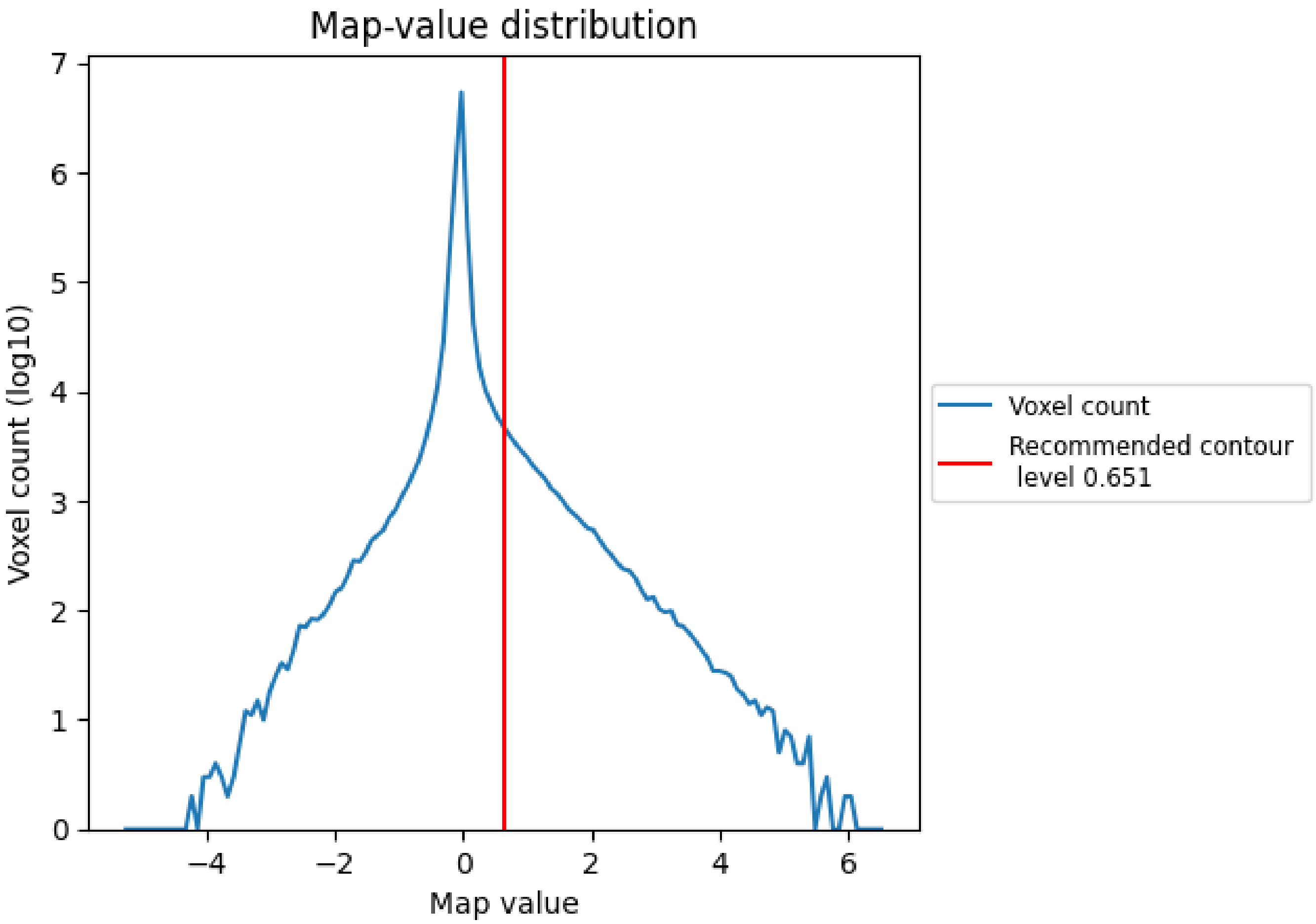


The map-value distribution is plotted in 128 intervals along the x-axis. The y-axis is logarithmic. A spike in this graph at zero usually indicates that the volume has been masked.

For Manuscript Review

#### 7.2 Volume estimate (i)

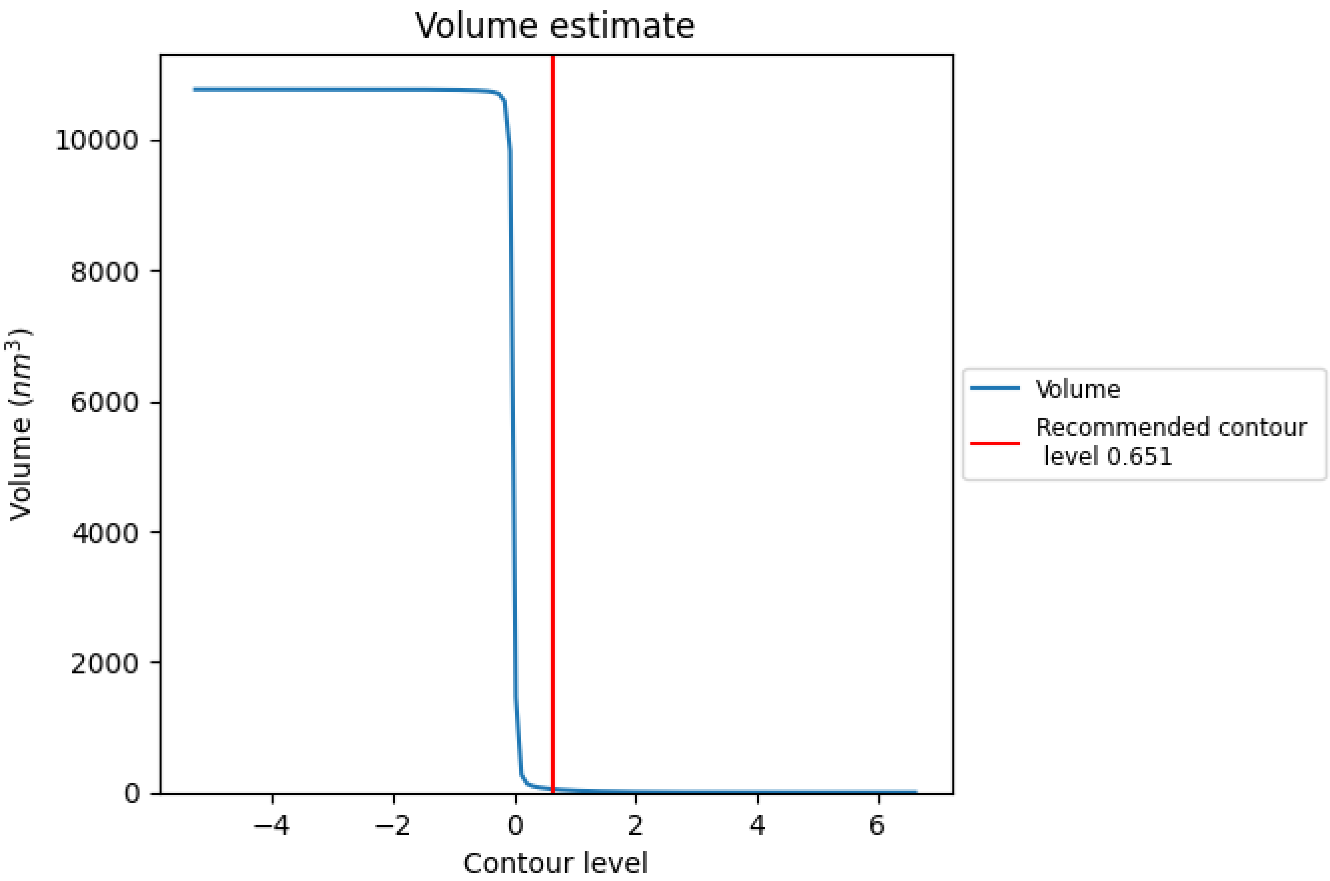


The volume at the recommended contour level is 49 $nm^3$; this corresponds to an approximate mass of 44 kDa.

The volume estimate graph shows how the enclosed volume varies with the contour level. The recommended contour level is shown as a vertical line and the intersection between the line and the curve gives the volume of the enclosed surface at the given level.

For Manuscript Review

#### 7.3 Rotationally averaged power spectrum (i)

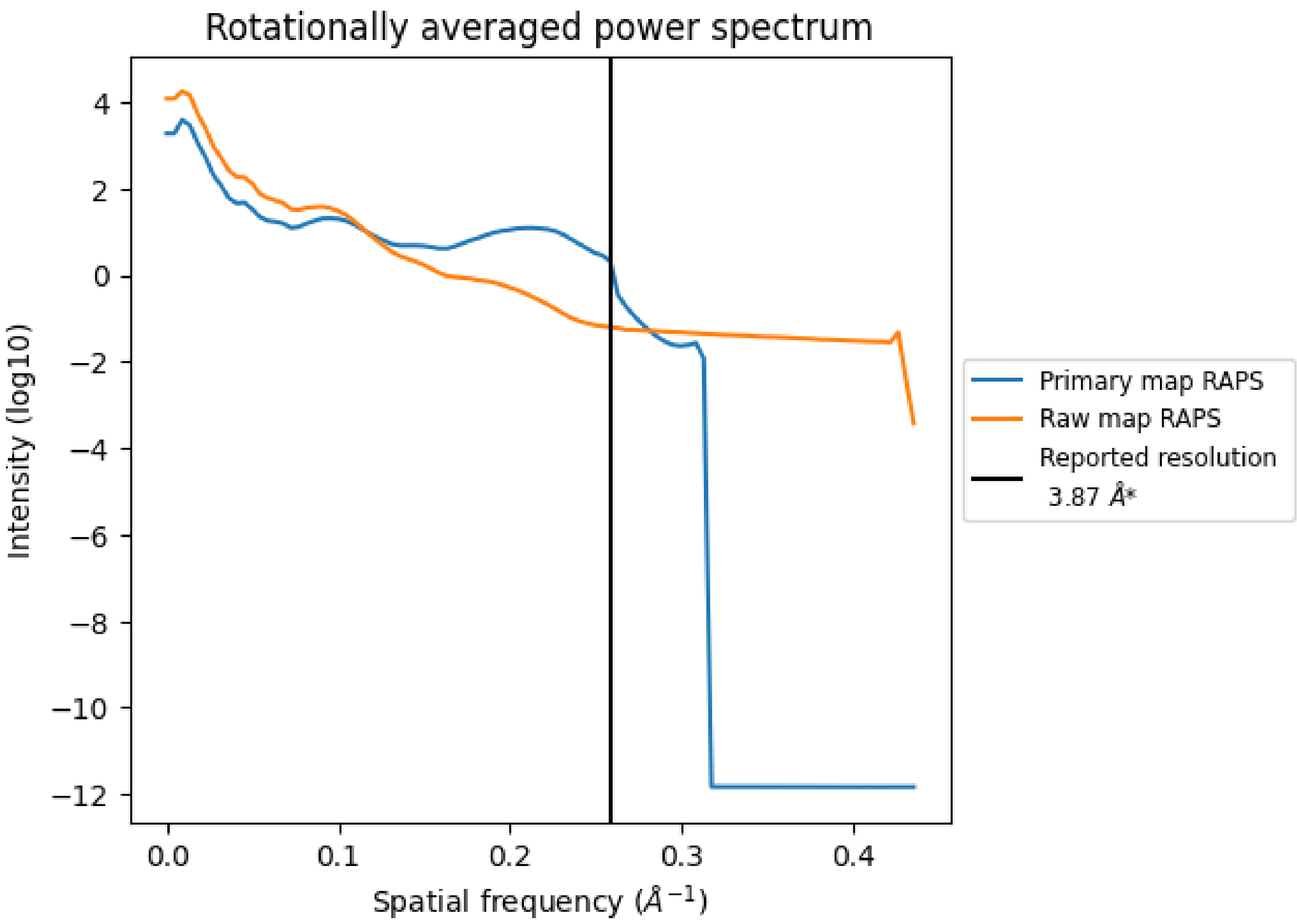


*Reported resolution corresponds to spatial frequency of 0.258 Å$^{-1}$

For Manuscript Review

### 8 Fourier-Shell correlation ⓘ

Fourier-Shell Correlation (FSC) is the most commonly used method to estimate the resolution of single-particle and subtomogram-averaged maps. The shape of the curve depends on the imposed symmetry, mask and whether or not the two 3D reconstructions used were processed from a common reference. The reported resolution is shown as a black line. A curve is displayed for the half-bit criterion in addition to lines showing the 0.143 gold standard cut-off and 0.5 cut-off.

#### 8.1 FSC ⓘ

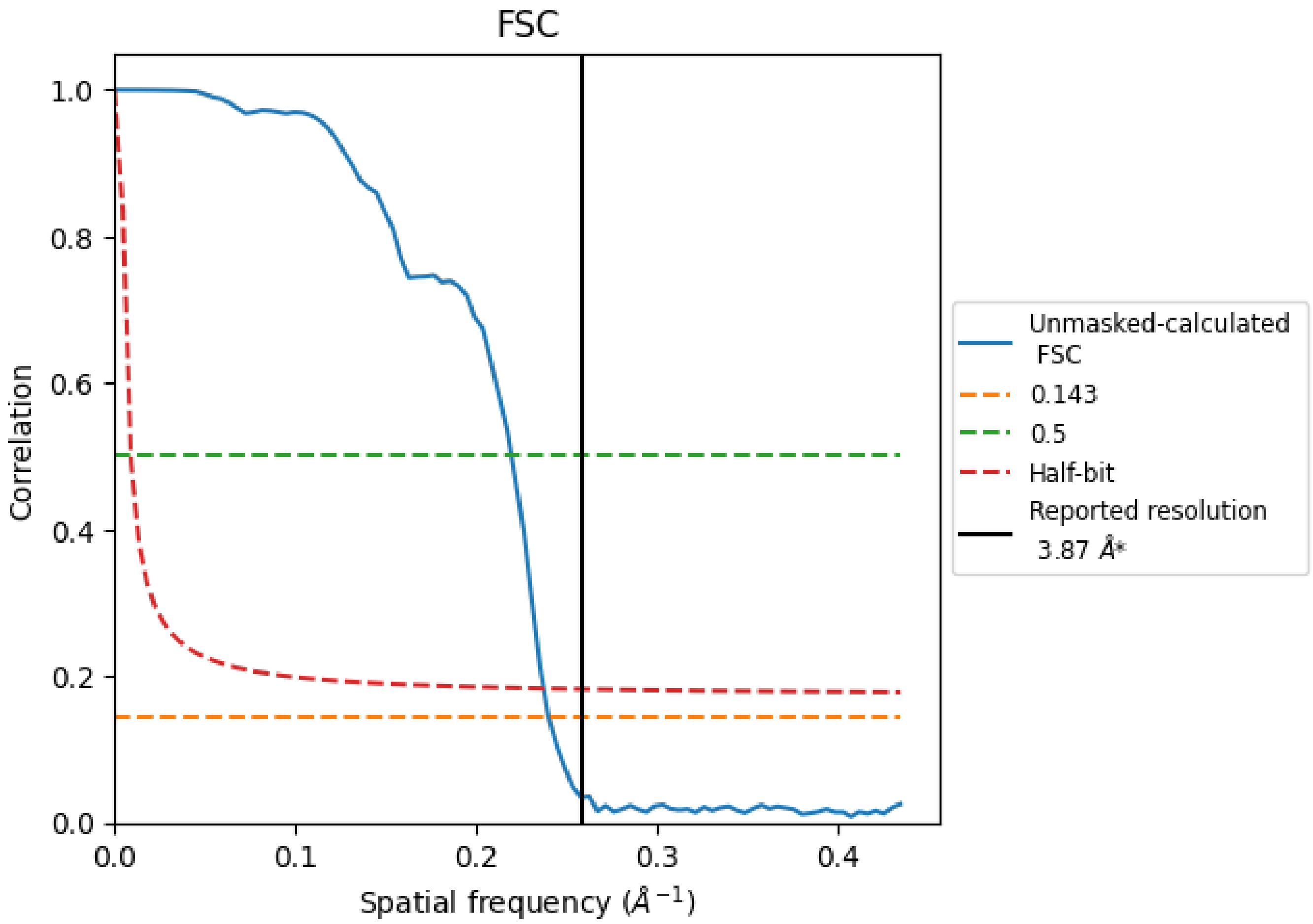


*Reported resolution corresponds to spatial frequency of 0.258 Å$^{-1}$

For Manuscript Review

#### 8.2 Resolution estimates (i)

| Resolution estimate (Å) | Estimation criterion (FSC cut-off) | | |
|---|---|---|---|
| | 0.143 | 0.5 | Half-bit |
| Reported by author | 3.87 | - | - |
| Author-provided FSC curve | - | - | - |
| Unmasked-calculated* | 4.16 | 4.55 | 4.21 |

*Resolution estimate based on FSC curve calculated by comparison of deposited half-maps.

For Manuscript Review

### 9 Map-model fit (i)

This section contains information regarding the fit between EMDB map EMD-47103 and PDB model 9DPD. Per-residue inclusion information can be found in section 3 on page 5.

#### 9.1 Map-model overlay (i)

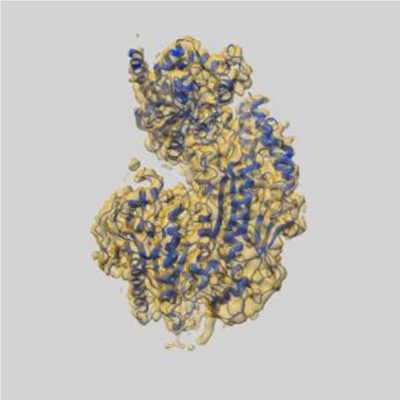

X

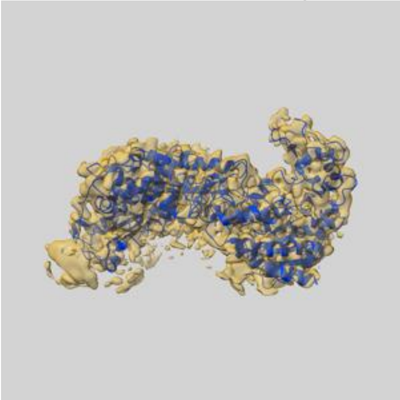

Y

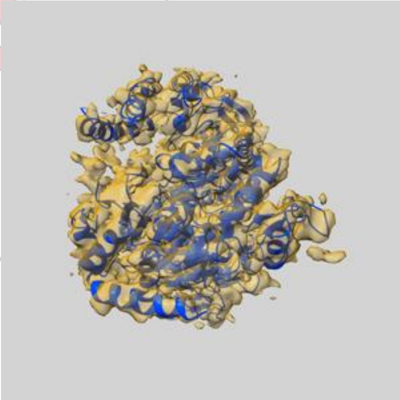

Z

The images above show the 3D surface view of the map at the recommended contour level 0.651 at 50% transparency in yellow overlaid with a ribbon representation of the model coloured in blue. These images allow for the visual assessment of the quality of fit between the atomic model and the map.

For Manuscript Review

#### 9.2 Q-score mapped to coordinate model (i)

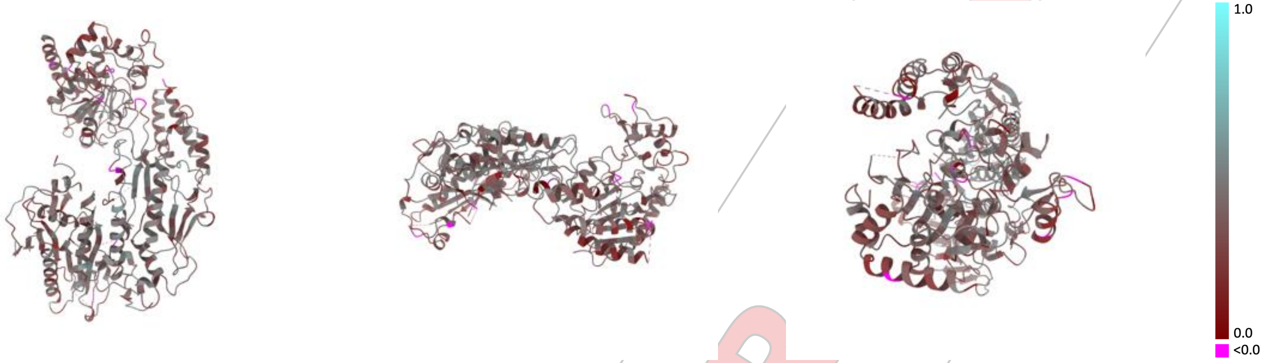


X Y Z

The images above show the model with each residue coloured according its Q-score. This shows their resolvability in the map with higher Q-score values reflecting better resolvability. Please note: Q-score is calculating the resolvability of atoms, and thus high values are only expected at resolutions at which atoms can be resolved. Low Q-score values may therefore be expected for many entries.

#### 9.3 Atom inclusion mapped to coordinate model (i)

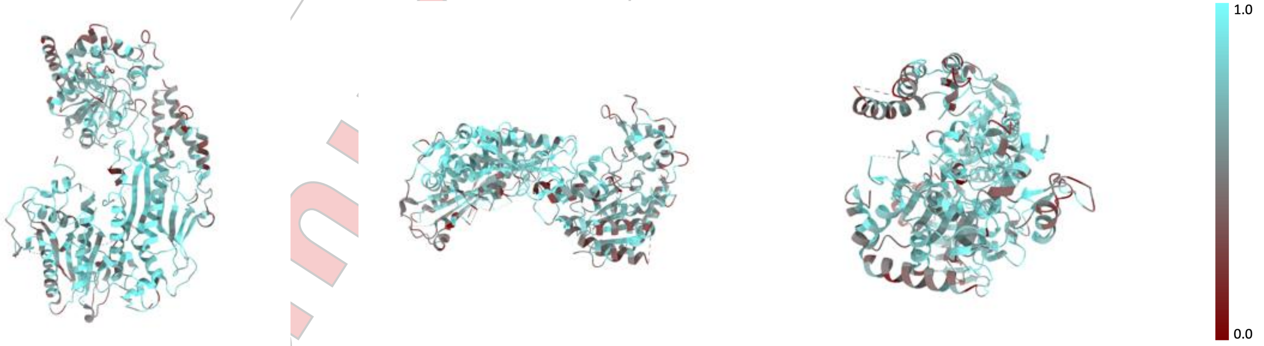


X Y Z

The images above show the model with each residue coloured according to its atom inclusion. This shows to what extent they are inside the map at the recommended contour level (0.651).

For Manuscript Review

#### 9.4 Atom inclusion (i)

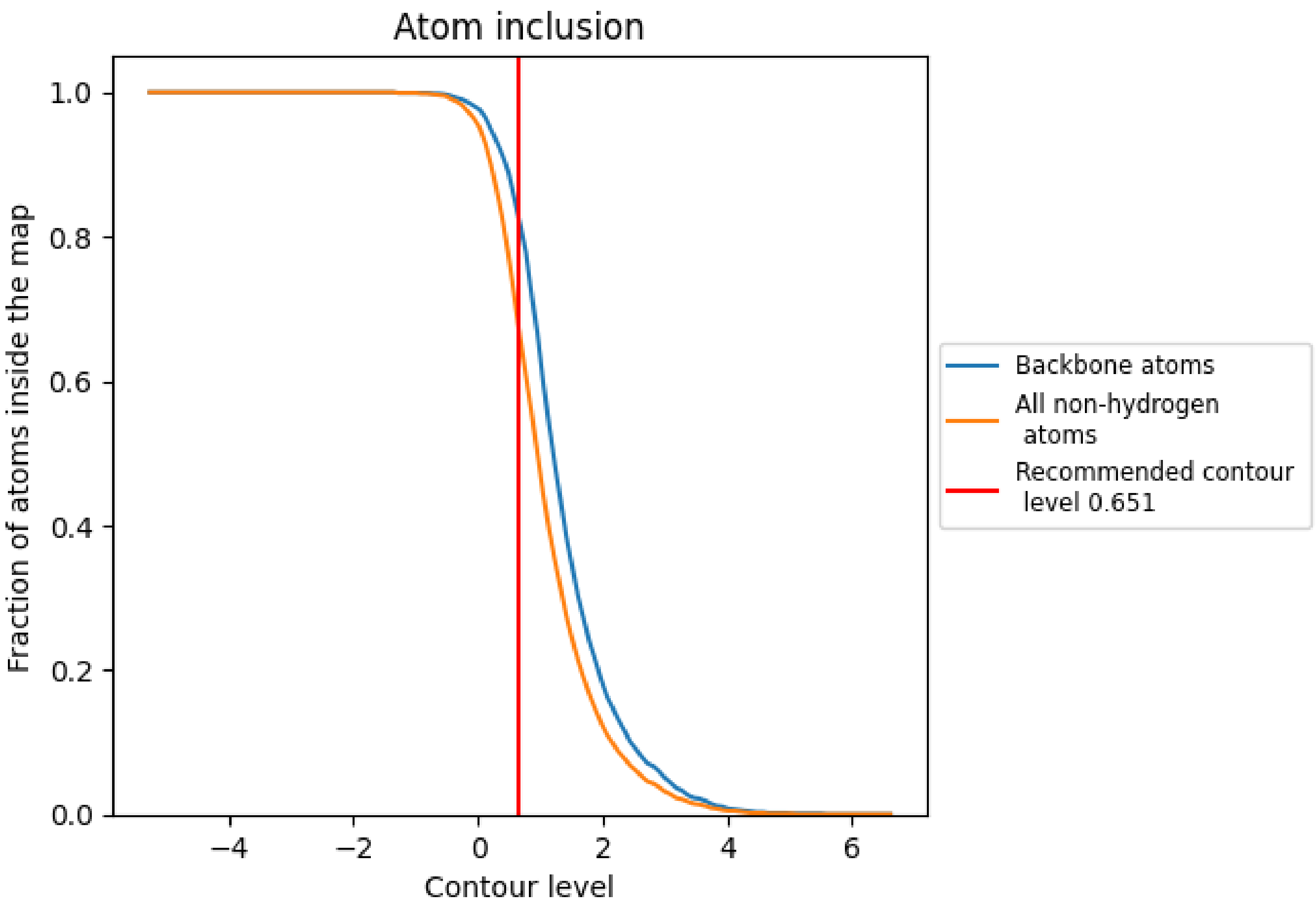


At the recommended contour level, 83% of all backbone atoms, 68% of all non-hydrogen atoms, are inside the map.

For Manuscript Review

#### 9.5 Map-model fit summary (i)

The table lists the average atom inclusion at the recommended contour level (0.651) and Q-score for the entire model and for each chain.

| Chain | Atom inclusion | Q-score |
|---|---|---|
| All | 0.6770 | 0.3600 |
| B | 0.6670 | 0.3520 |
| C | 0.6360 | 0.3420 |
| D | 0.7160 | 0.3800 |

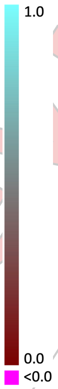


For Manuscript Review
